# Molecular signatures of local adaptation to light in Norway Spruce: role of lignin mediated immunity

**DOI:** 10.1101/2020.01.14.905810

**Authors:** Sonali Sachin Ranade, María Rosario García-Gil

## Abstract

Study of natural variation is an efficient method to elucidate how plants adapt to local climatic conditions, a key process for the evolution of a species. However, it is challenging to determine the genetic basis of adaptive variation especially in forest trees which have large and complex genomes. Norway spruce is a shade tolerant conifer in which the requirement of far-red light for growth increases latitudinally northwards. In the current work, hypocotyl-length followed a latitudinal cline in response to SHADE (low red:far-red ratio). RNA-sequencing revealed differential gene expression in response to SHADE, between a southern and a northern natural population in Sweden. Exome capture included analysis of uniquely large data set (1654 trees) that revealed missense variations in coding regions of nine differentially expressed candidate genes, which followed a latitudinal cline in allele and genotype frequencies. These genes included five transcription factors involved in vital processes like bud-set/bud-flush, lignin pathway and cold acclimation, and other genes that take part in cell-wall remodeling, secondary cell-wall thickening, response to starvation and immunity. Findings from this work primarily suggests that the northern populations of Norway spruce are better adapted towards disease resistance under shade by up-regulation of lignin pathway that is linked to immunity and it forms concrete basis for local adaptation to light quality in Norway spruce, one of the most economically important conifer tree species in Sweden.

## Introduction

Research indicates that local adaptation in conifer populations will be affected by major disturbances due to increase in average temperature, change in precipitation regimes and rise in new biotic stresses, which could compromise forest survival and sustainability at their present locations (Leinonen, 1996, Millar and Stephenson, 2015). Given the prognosis of a rapid climatic change (IPCC, 2019), forest breeders and conservationists are urging for a better understanding of the genetics and genomic basis underlying local adaptation. This is crucial to design effective spatial migration actions to transfer forest trees into their optimal ecological niches or to assist the trees to adapt to the new conditions at their current location (Aitken *et al.*, 2008). Solving the basis of local adaptation at the molecular level has been at the forefront of genetics research aiming for clues on genomic regions of functional relevance (Hoban *et al.*, 2016).

Differences in sensitivity to the spectral quality of light between Norway spruce (*Picea abies* L. Karst.) populations has been demonstrated as early as in 1978 (Ekberg *et al.*, 1978). In the current study, we aimed to identify the genomic signals of local adaptation in response to low Red:Far-red (R:FR) ratio (SHADE) in this species which is shade tolerant. R:FR ratio is considered as a reliable indicator of the degree of shade and low values of it can activate shade-avoidance and shade-tolerance responses (Ranade *et al.*, 2019). In this species, such variation is characterized by an increase in requirement of FR (lower R:FR) with latitude in order to maintain growth (Clapham *et al.*, 1998, Mølmann *et al.*, 2006). Similar latitudinal response has been described in *Salix penta nd ra* L. and Scots pine (*Pinus sylvestris* L.) (Juntilla and Kaurin, 1985, Clapham *et al.*, 2002, Mølmann *et al.*, 2006). This latitudinal variation has been suggested to be an adaptive response to the prolonged end-of-day (EOD) FR-enriched light condition (twilight) (Juntilla and Kaurin, 1985), which characterizes the northern latitudes during the summer solstice (Nilsen, 1985). Thus, trees in the north may have adopted a light system that utilizes FR as a primary cue for growth regulation, while at the southern latitudes duration of darkness may be a more critical factor (Juntilla and Kaurin, 1985, Clapham *et al.*, 1998). In other words, northern trees have optimized growth under low R:FR ratios. EOD FR-enriched and shade tolerance are both triggered by low R:FR ratio and by the existence of common molecular components regulating both mechanisms (Johnson *et al.*, 1994, Muller-Moule *et al.*, 2016).

Twilight is a consequence of the 24-hour cycle during which light changes as a function of the Earth’s rotation in both intensity and spectral composition, being characterized by a daily reduction in R:FR at dawn and dusk due to increase in FR light following atmospheric refraction (Smith, 1981). The length of the twilight periods and the extent of R:FR reduction increases towards the northern latitudes. As a proxy to twilight exposure, in the current work, seedlings were treated with two contrasting values of bi-chromatic R:FR treatments representing day-light (R:FR = 1.2) or twilight period or SHADE (R:FR = 0.2). Thus, we first studied the effect of SHADE or low R:FR on morphological trait (hypocotyl length) in seedlings of Norway spruce sampled from natural populations across four latitudes in Sweden. RNA sequencing (RNA-Seq) analysis was conducted in seedlings originating from two populations farthest apart (south, 56°2’N and north, 67°2’N) as the phenotypic (hypocotyl length) difference in response to SHADE was the highest in these two latitudes, leading to identification of differentially expressed genes (DEGs) under SHADE. In the second step with the aid of exome capture, we estimated changes in the allele/genotype frequency with reference to the DEGs in six populations latitudinally distributed across Sweden (1654 trees). Given the described experimental approach, this study aimed to answer the following specific questions: (i) Do we observe a phenotypic cline in the Norway spruce seedlings in response to low R:FR or SHADE? (ii) Knowing that Norway spruce seem to be adapted to the local light quality conditions (Clapham *et al.*, 1998), what are the genomic signs of local adaptation for the exposure to SHADE?

## Results

### Effect of SHADE on seedling morphology

There was a significant increase in the length of hypocotyl (*p-value*<0.01, Fig. 1) towards the northern latitudes in response to SHADE.

**Fig. 1.**
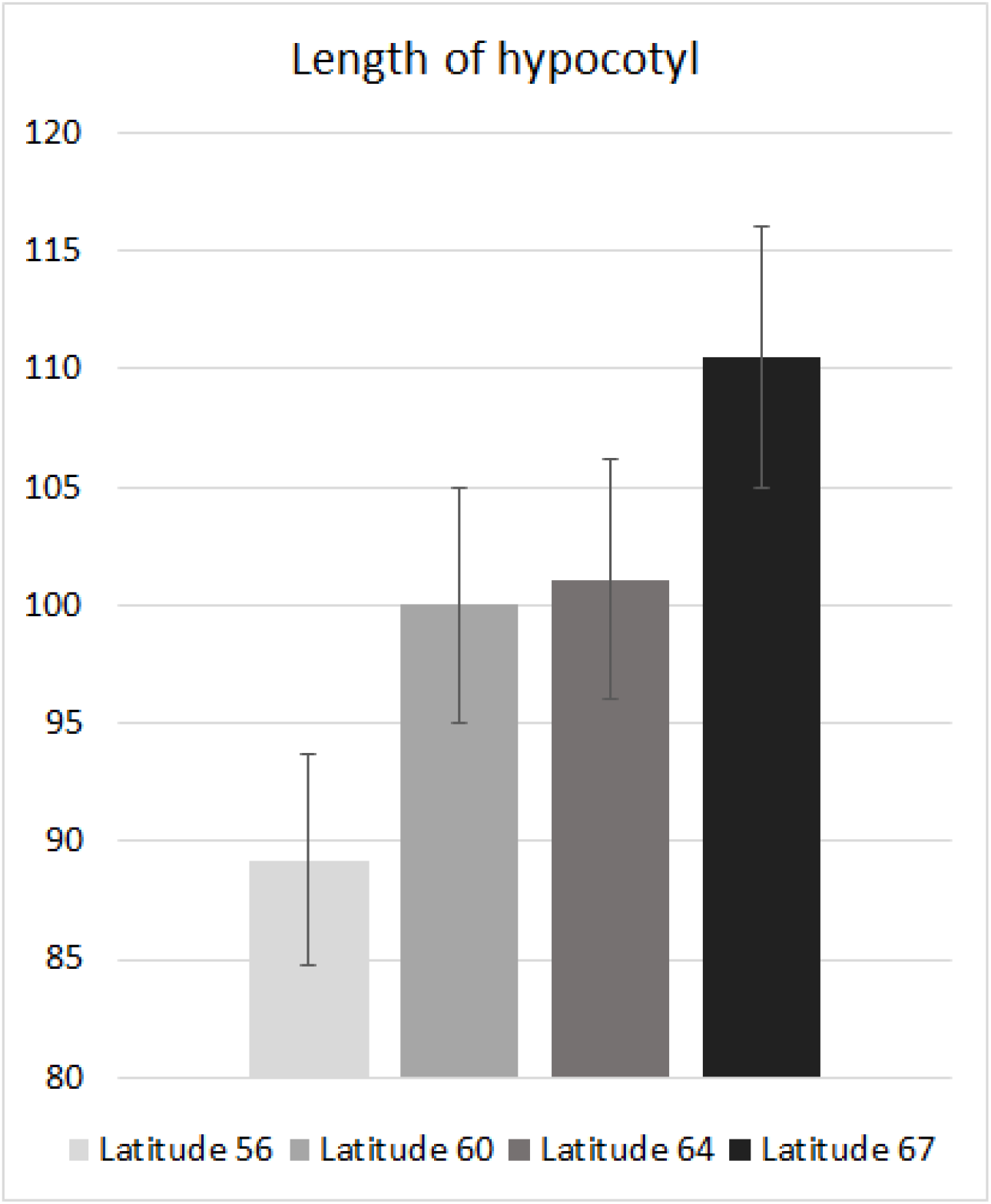
Bar plot with mean and ±SE of the proportion of change of hypocotyl length in response to SHADE as compared to SUN condition (*p-value*<0.01) across four latitudes in Sweden that shows clinal variation.

### Differentially expressed genes (DEGs) and latitudinal cline in SNPs

RNA-Seq analysis showed differential gene expression in response to SHADE in the seedling from the southern (56°2’N) and northern populations (67°2’N) of Norway spruce. Since we compared the expression data between the two latitudes in response to SHADE using SUN as the control, same set of genes that were up-regulated at latitude 67°2’N, were also down-regulated at latitude 56°2’N and vice-versa. 141 genes were found to be up-regulated in the population of Norway spruce at latitude 67°2’N (SHADE-67>SHADE-56) and the same genes were down-regulated at latitude 56°2’N in response to SHADE (Supplementary Table S1, Supplementary Material online). Likewise, 132 genes were detected to be up-regulated at latitude 56°2’N (SHADE-67<SHADE-56) and the same set of genes were found to be down-regulated at 67°2’N, in response to SHADE (Supplementary Table S2, Supplementary Material online).

Gene ontology (GO) analysis revealed that there were significantly (*p-value*=0.05) higher number of defense related genes (15 genes) that were up-regulated in the northern population under SHADE as compared to the southern population (6 genes). The metabolic pathway maps (Supplementary Fig.S1-S5, Supplementary Material online) that were constructed with reference to the DEGs in the two populations showed that higher number of genes were up-regulated in cell wall at latitude 67°2’N as compared to latitude 56°2’N (Supplementary Fig.S1, Supplementary Material online). Likewise, higher number of genes involved in the phenylpropanoid pathway were up-regulated at latitude 67°2’N (Supplementary Fig.S1-S3, Supplementary Material online). Differential expression of equal number of transcription factors was detected in both latitudes (10 and eight TFs were up-regulated in the north and south, respectively; *p-value*>0.05. Supplementary Fig.S4, Supplementary Material online).

Two light intensity/shade responsive genes were detected to be up-regulated under SHADE in the northern population – *DRACULA2 (DRA2)* involved in hypocotyl elongation and control of shade induced gene expression (Gallemi *et al.*, 2016) and *PHYTOCHROME RAPIDLY REGULATED 2* (*PAR2*) that encodes a protein which is up-regulated under shade and acts as negative regulator of shade avoidance response (Roig-Villanova *et al.*, 2006). One light-regulated gene was detected to be up-regulated in the southern population - *TEOSINTE BRANCHED 1, CYCLOIDEA AND PCF TRANSCRIPTION FACTOR 2* (*TCP2*); *TCP2* is a positive regulator of photomorphogenesis and is involved in response to blue light (He *et al.*, 2016). Cell wall related genes - *CELLULOSE SYNTHASE A4* (*CESA4*) involved in secondary cell wall biosynthesis (Endler and Persson, 2011) and *CINNAMOYL COA REDUCTASE* (*CCR2*) involved in lignin biosynthesis (Wadenback *et al.*, 2008) were found to be up-regulated in the northern trees. Likewise, immunity specific genes were detected to be up-regulated at the northern latitude - (*RESISTANT TO P. SYRINGAE 2* (*RPS2*) (Kunkel *et al.*, 1993) and *L-TYPE LECTIN RECEPTOR KINASE S.4* (*LECRK-S.4*) (Wang and Bouwmeester, 2017)).

We considered bi-allelic SNPs for the study. Latitudinal cline in SNP represents increase in frequency of either allele along with decrease in the frequency of the other with latitude, which also reflects in the genotype frequencies of the particular SNP, meaning that there is increase/decrease of either of the homozygotes and the corresponding change in the frequency of heterozygotes with latitude. RNA-Seq analysis revealed a total of 273 genes that were differentially regulated in response to SHADE between the two populations from two latitudes (56°2’N and 67°2’N), out of which 56 genes were detected with at least one missense SNP. We focused only on the missense mutations as those would have an effect on the protein conformation, which may contribute towards the differential regulation of the gene expression in response to SHADE or low R:FR. Missense mutations following a latitudinal cline for allele frequencies across the six Norway spruce populations were detected in 46 out of the 56 genes. In this article, we have discussed only nine such relevant genes (Table 1) that showed statistically significant and precise latitudinal cline in the missense nucleotide variations, and which are also well characterized in the literature. Since it is beyond the scope of this article to discuss all the remaining DEGs that showed cline, we have included the details regarding the allele frequencies of these missense mutations in the supplementary material (Supplementary Table S3, Supplementary Material online). Nevertheless, we have considered all the 56 genes to estimate genetic diversity across the populations included in the study. These genes were differentially regulated in response to SHADE and therefore might be novel regulators in this context.

**Table 1.**
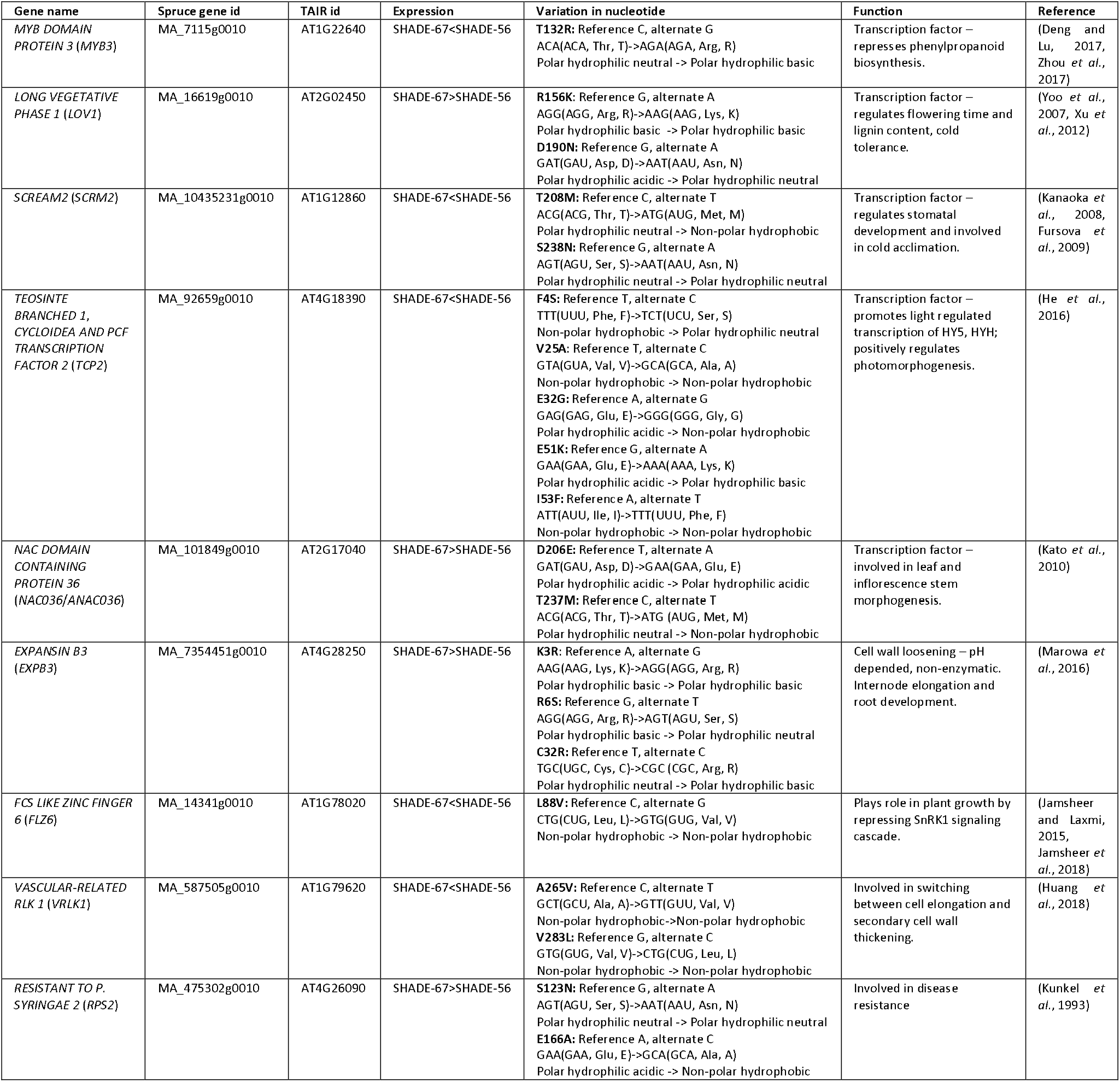
Differentially expressed genes in response to SHADE in the southern and northern Norway spruce populations, which showed latitudinal cline in SNPs of the genes differentially regulated under SHADE

The allele frequencies and genotype frequencies of SNPs in nine DEGs followed a latitudinal cline (Fig. 2; Table 2; Supplementary Fig. S6-S13, Supplementary Material online) which included five transcription factors involved in photomorphogenesis (*TCP2*), lignin pathway (*MYB DOMAIN PROTEIN 3, MYB3; LONG VEGETATIVE PHASE 1, LOV1*), stomatal development and cold acclimation (*SCREAM 2, SCRM2*) and, leaf and inflorescence stem morphogenesis (*NAC DOMAIN CONTAINING PROTEIN 36, NAC036*) (Table 1). Other DEGs genes that showed cline in sequence variations were – *EXPANSIN B3* (*EXPB3*) that controls the cell wall remodeling, *FCS LIKE ZINC FINGER 6* (*FLZ6*) which is involved in response to starvation and *VASCULAR-RELATED RLK 1* (*VRLK1*) that regulates secondary cell wall thickening and a gene potentially involved in plant immunity (*RESISTANT TO P. SYRINGAE 2* (*RPS2*)).

**Fig. 2.**
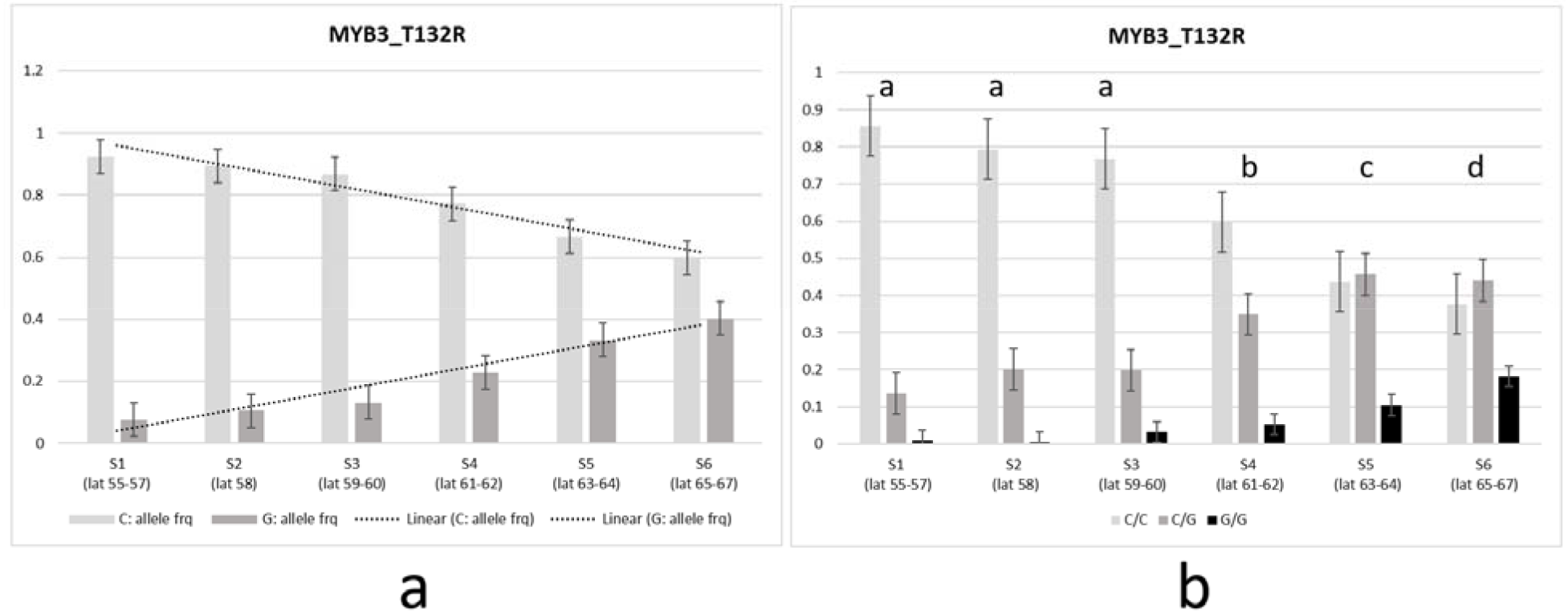
Cline with reference to variation in allele and genotype frequencies of SNP in the *MYB3* gene in Norway spruce populations across Sweden.

a. allele frequencies of T132R
b. genotype frequencies of T132R, Tukey’s *post-hoc* categorization is indicated above the bars.

**Table 2.**
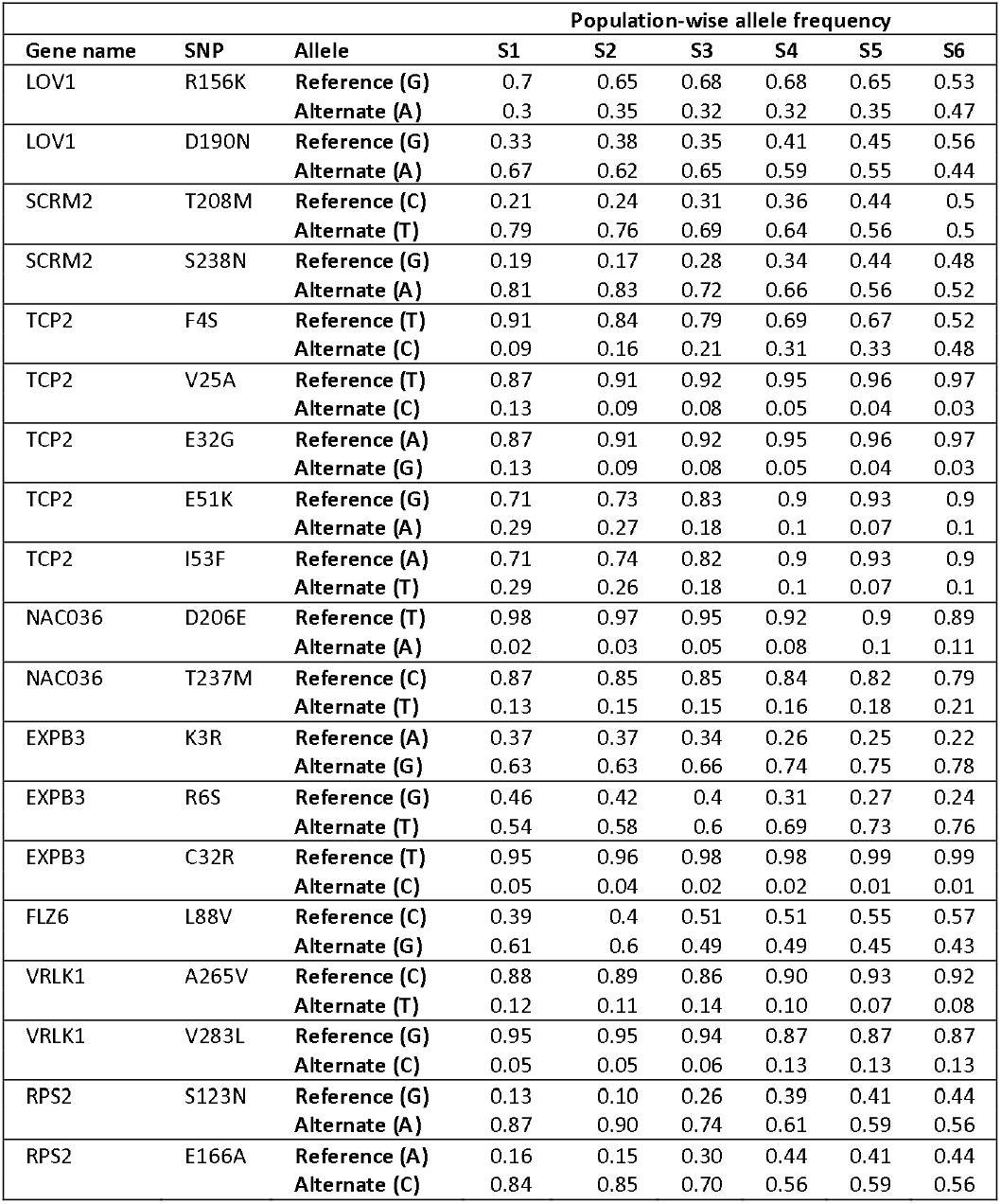
Population-wise allele frequency of SNPs showing latitudinal cline in candidate genes differentially regulated under SHADE

The allele frequencies of the missense SNPs for the nine candidate genes were found to be significantly different across all the populations included in the study except in case of few SNPs; however, the genotype frequencies were observed to be significantly different in all populations (Supplementary Table S4, Supplementary Material online), which was further subjected to Tukey’s *post-hoc*(Supplementary Table S5, Supplementary Material online). The location of the missense mutations in the nine candidate genes with reference to a particular domain of the respective gene is included in Supplementary Material (Supplementary Fig.S14-S22, Supplementary Material online).

Majority of the ecotypes were found to be in HWE with reference to the missense SNPs detected in the DEGs, with few exceptions in case of *LOV1* (D190N), *TCP2* (F4S), *TCP2* (T518K), *EXPB3* (R6S), *FLZ6* (L88V) and *VRLK1* (V283L) where it showed deviation from HWE (Supplementary Table S6, Supplementary Material online). Pairwise *F_ST_* estimates for all the six populations (Table 3), calculated considering both the synonymous + missense SNPs of the 56 DEGs, suggest that differentiation increases with the geographic distance. Cline across the latitudes was also observed in few of the synonymous substitutions with reference to the nine candidate genes – TCP2, *LOV1, FLZ6* and *NAC036* (Supplementary Table S7, Supplementary Material online). The distribution of R^2^ and *F_ST_* across the nine candidate genes is represented in Fig. 3.

**Fig. 3.**
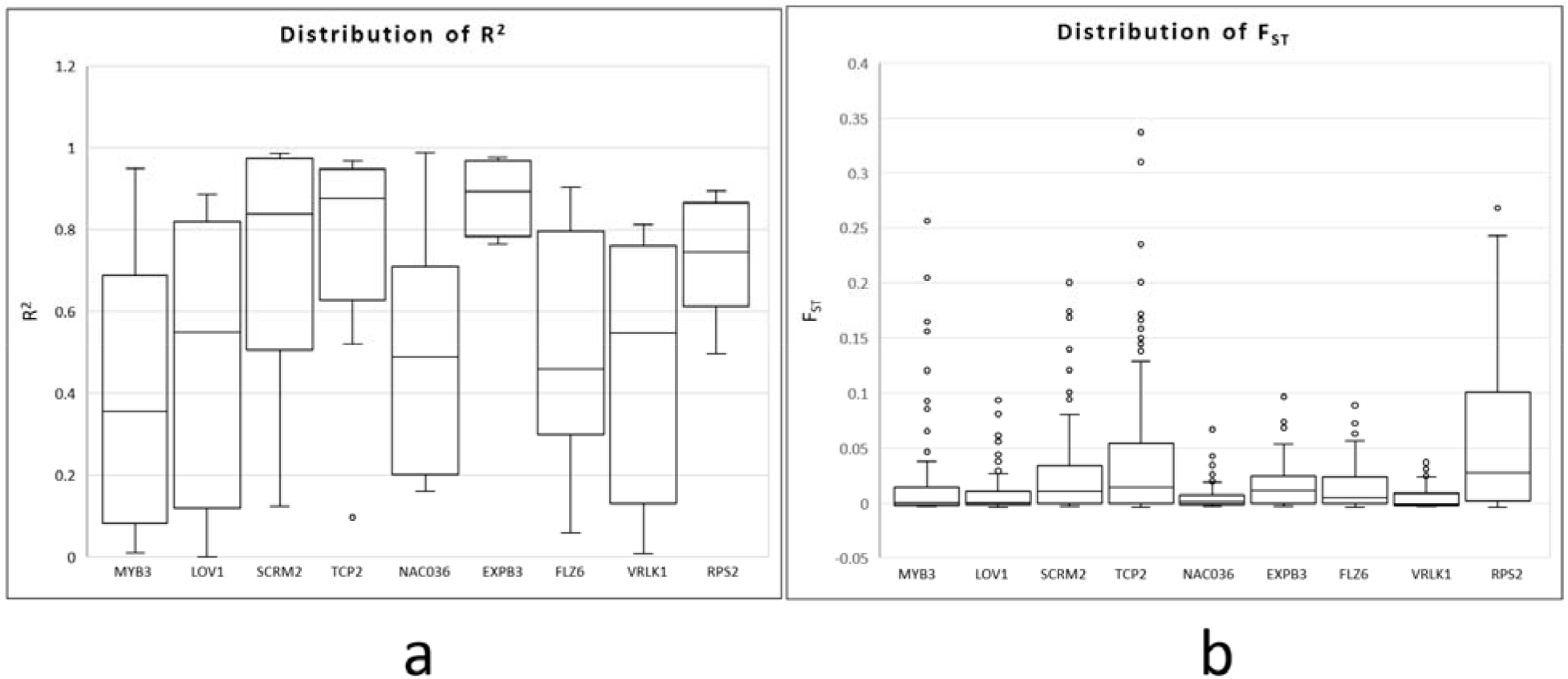
The distribution of (a) R^2^ with reference to allele frequencies and (b) *F_ST_*, across the nine candidate genes showing cline in SNPs and which are differentially regulated in response to SHADE in southern and northern Norway spruce populations Sweden. R^2^ and *F_ST_*-was calculated considering the missense and the synonymous SNPs detected in the particular candidate gene.

**Table 3.** Pairwise *F_ST_* estimates for all populations across Sweden involved the analysis for detection of latitudinal cline in SNPs of the genes differentially regulated under SHADE.

| Population | S2 | S3 | S4 | S5 | S6 |
| --- | --- | --- | --- | --- | --- |
| S1 | -2.87E-05 | 0.003961 | 0.016808 | 0.014853 | 0.029082 |
| S2 |  | 0.00115 | 0.005008 | 0.000645 | 0.024402 |
| S3 |  |  | 0.002349 | 0.004307 | 0.009299 |
| S4 |  |  |  | -0.00014 | -0.00074 |
| S5 |  |  |  |  | 0.001578 |

### Types of the missense polymorphisms in the nine candidate genes

Majority of the missense polymorphisms in the candidate genes resulted in alteration to the amino acid with similar chemical property (e.g. polar/hydrophilic to polar/hydrophilic), with a few exceptions (Table 1). MYB3, LOV1, EXPB3, FLZ6 and VRLK1 showed alterations to the amino acid with similar chemical property whereas one of the polymorphisms from SCRM2, NAC036 and RPS2, and two from TCP2 showed changes in the amino acids that resulted in change in the chemical property, e.g. from polar/hydrophilic to non-polar/hydrophobic and vice-versa.

## Discussion

Higher FR content (lower R:FR ratio) is known to trigger both EOD-FR and response to vegetative shade. In addition, these FR-related processes have been shown to share common molecular components (Johnson *et al.*, 1994, Muller-Moule *et al.*, 2016). These two facts encourage the speculation about a latitudinal cline for the level of shade tolerance (equivalent to that for EOD-FR). Our study has revealed a latitudinal cline for hypocotyl elongation in response to shade, which indicates that previously reported cline for growth regulation by FR-enrich light (Clapham *et al.*, 1998) directly translates into a cline for shade tolerance.

### Latitudinal clines in the allele and genotype frequencies of the DEGs

Pairwise *F_ST_* estimates (Table 3) increase with geographic (latitudinal) distance between ecotypes, although in general the values are low, which agree with previous reports in Norway spruce (Chen *et al.*, 2016). This trend corresponds with significant latitudinal change in the allele/genotype frequencies of missense polymorphisms that suggests the involvement of the DEGs in the ecotype’s adaptation to the local light conditions. Our results together with previous studies in population differentiation in Norway spruce suggest that natural selection and to a less extent isolation by distance (Chen *et al.*, 2014) to be the main force shaping the observed latitudinal cline at the DGEs. As compared to previous studies (Pyhajarvi *et al.*, 2007, Chen *et al.*, 2014, Kallman *et al.*, 2014, Chen *et al.*, 2016), we applied transcriptomic analysis to identify strong candidate genes to explain the genetic basis of latitudinal adaptation.It is relevant to highlight that exome capture does not aim to sequence the entire coding sequence of the gene; therefore, it is possible that we have missed some SNPs that are under selection for the simple reason that they are located in the non-sequenced exons. This could explain why some of the missense SNPs in differentially expressed genes do not exhibit a cline. Moreover, some of the neutral SNPs showing a latitudinal cline may be in linkage disequilibrium with SNPs under selection and therefore, they may appear associated to the adaptive cline. The four genes with the highest *F_ST_* values (SCRM2, TCP2, EXP3, RPS2) also exhibit higher and less dispersed R^2^ values as a result of a steeper latitudinal cline (Fig. 3). As expected from the global *F_ST_* estimates (including 56 DEGs, Table 3), the *F_ST_* values are low and show different levels of dispersion upwards, which can be explained by an increase in differentiation with pair-wise geographic distance.

### Alteration in protein conformation

The missense polymorphism T208M in the *SCRM2* gene involves a change in the amino acid from polar/hydrophilic to non-polar/hydrophobic similar to that of the polymorphisms in *TCP2* (E32G), *NAC036* (T237M) and *RPS2* (E166A) (Table 1). Likewise, there is a change in the amino acid from non-polar/hydrophobic to polar/hydrophilic in *TCP2* gene (F4S). These polymorphisms are likely to alter the protein folding and conformation, consequently influencing their binding-capacity/interaction, leading to alteration in their mode of action. Although the missense polymorphisms reported in this study were not located in the specific conserved domains of the genes (Supplementary Fig.S14-S22, Supplementary Material online), e.g. the DNA binding domain of the particular TF, it will still affect the mode of action of the protein by altering its conformation.

### Cell wall and lignin pathway

Comparison of the expression pattern (RNA-Seq) under SHADE treatment between the southern and northern Norway spruce populations where SUN was used as control treatment lead to identification of DEGs in response to SHADE. Cline was observed with reference to variation in allele and genotype frequencies of SNPs in nine DEGs, of which two genes represented transcription factors related to lignin - *MYB3* involved in lignin biosynthesis (Zhou *et al.*, 2017) and *LOV1* which regulates flowering time and, lignin content and composition (Xu *et al.*, 2012). MYB3 represses cinnamate 4-hydroxylase (C4H) gene expression, which catalyzes the second step of the main phenylpropanoid pathway leading to the synthesis of lignin and pigments (Liu *et al.*, 2018). *LOV1* controls flowering time within the photoperiod pathway in *Arabidopsis thaliana* (*A. thaliana*) by negatively regulating expression of *CONSTANS (CO)* which is a floral promoter (Yoo *et al.*, 2007). Overexpression of *LOV1* leads to delayed flowering time and increased lignin content in Switchgrass (Xu *et al.*, 2012). One gene involved in cell wall synthesis (*VRLK1*) also showed cline in the detected SNPs, while other two genes - *CESA4* and *CCR2* were differentially regulated under SHADE, although did not show cline with reference to SNPs. *VRLK1* encodes leucine-rich repeat receptor-like kinase which is expressed specifically in cells undergoing secondary cell wall thickening (Torii, 2004). Secondary cell wall contains additional lignin as compared to the primary cell wall. *VRLK1* regulates secondary cell wall thickening; down-regulation of *VRLK1* promotes secondary cell wall thickening and up-regulation of *VRLK1* inhibits it in *A. thaliana* (Huang *et al.*, 2018). *CESA4* is up-regulated in the northern trees, suggesting enhanced secondary cell wall biosynthesis. In the context on differential expression between the southern and northern Norway spruce populations in response to SHADE, *MYB3* was down-regulated at latitude 67°2’N as compared to latitude 56°2’N and *LOV1* was found to be up-regulated at latitude 67°2’N as compared to latitude 56 °2’N. Furthermore, up-regulation of *CCR2* at latitude 67°2’N supports the increase in lignin biosynthesis as suppression of CCR results in reduction in lignin content in Norway spruce (Wadenback *et al.*, 2008). This suggests that the lignin pathway was enhanced at latitude 67°2’N as compared to latitude 56°2’N under SHADE, which may be co-related with the SNPs detected in the respective DEG. Additionally, *VRLK1* expression was lower in the northern trees than the southern ones under SHADE, which again indicates that secondary cell wall thickening (lignin deposition) was enhanced in the northern populations. Also, higher number of cell wall genes were up-regulated at latitude 67°2’N (Supporting Material online 2, Fig.S1). This phenomenon can be attributed due to the variation in the SNPs in the respective genes observed in this study that follows a latitudinal cline across Sweden. Although shade stress decreases lignin leading to weak stem in many of the plant species (Wang *et al.*, 2012, Wu *et al.*, 2017, Hussain *et al.*, 2019, Liu *et al.*, 2019b), the northern spruce populations appear to have adapted to SHADE or low R:FR ratio or in other words higher requirement of FR to maintain the growth and other regular plant processes (Clapham *et al.*, 1998, Clapham *et al.*, 2002, Ranade and García-Gil, 2013), which may correspond to the observed up-regulation of cell wall related genes. It is worth mentioning that none of the ecotypes deviated from HWE in case of *MYB3*, which suggests that the particular genetic variation in *MYB3* gene (T132R) will remain constant (i.e., not evolving) in entire population from one generation to the next in the absence of selecting forces.

### Cold tolerance

*SCRM2* encodes *INDUCER OF CBF EXPRESSION 2* (*ICE2*) which participates in the response to deep freezing; overexpression of *ICE2* results in increased tolerance to deep freezing stress after cold acclimation (Fursova *et al.*, 2009). Gain of function of *LOV1* gene confers cold tolerance in *A. thaliana*(Yoo *et al.*, 2007). *SCRM2* was down-regulated and *LOV1* was up-regulated in response to SHADE at latitude 67°2’N as compared to latitude 56°2’N, and this coupled with the clinal variation in the respective gene sequences conveys that the trees at different latitudes may have different molecular mechanisms to adapt to the cold environmental conditions prevailing in the northern latitudes.

### Photomorphogenesis and bud-burst

*TCP2* is a transcription factor that positively regulates photomorphogenesis; it promotes light regulated transcription of photomorphogenesis related genes e.g. *ELONGATED HYPOCOTYL 5* (*HY5*), *HY5* homolog (*HYH*) in *A. thaliana* (He *et al.*, 2016). *TCP2* controls cell-proliferation/division and growth, including floral meristem (Martin-Trillo and Cubas, 2010). *TCP2* is also associated with bud-burst, it was found to be up-regulated from quiescent bud to burst bud in tea (Liu *et al.*, 2019a). *TCP2* was detected to be down-regulated at latitude 67°2’N as compared to latitude 56°2’N in response to SHADE, suggesting that SHADE diminishes cell-division related activities in northern tree populations as compared to the southern ones, instead the northern trees seem to invest higher resources in cell wall thickening. This might influence the timing of bud-burst and bud-set under SHADE, the two traits that are known to follow a steep latitudinal cline in Norway spruce (Sogaard *et al.*, 2008, Chen *et al.*, 2012). Considering that bud-set and bud-burst are known to be closely linked responses and that the northern trees are more frost resistant (Westin *et al.*, 2000, Calleja-Rodriguez *et al.*, 2019, Sebastian-Azcona *et al.*, 2019), it warrants further research to verify the possible role of the interaction between phenology and cold tolerance related DEGs in adaptation to light quality.

### Stem development

*NAC036* is a transcription factor that regulates leaf and inflorescence stem morphogenesis. Overexpression of NAC036 resulted in a dwarf phenotype in *A. thaliana* (Kato *et al.*, 2010). This gene was found to be associated with the shade tolerant response in Norway spruce were it was reported to be up-regulated under SHADE conditions (Ranade *et al.*, 2019). In the current work, *NAC036* was up-regulated in response to SHADE in the northern trees as compared to the southern ones. Expansins comprise of a large gene family that codes for cell wall proteins which mediate the pH depended, non-enzymatic cell wall loosening and extension that plays vital role in plant cell growth and development (Marowa *et al.*, 2016). Differential expression of three expansins in response to SHADE was detected in both populations - *EXPANSIN-LIKE A1* (*EXLA1*) and *EXPANSIN A8* (*EXPA8*) that was down-regulated at latitude 67°2’N, and *EXPB3* which was up-regulated at latitude 67°2’N as compared to latitude 56°2’N (Supplementary Table S1-S2, Supplementary Material online). Out of these, only *EXPB3* showed clinal variation in the sequence. *EXPB3* in rice has been reported to be involved in stem/internode elongation as well as root development (Marowa *et al.*, 2016). Thus, gene responsible was stem development was detected to be differentially regulated in response to SHADE in both the populations.

### Regulation of plant growth

Biotic and abiotic stresses lead to energy deficit in the plant cell that triggers the transcription of genes enabling the plants to withstand and survive under low-energy conditions. The SNF1-RELATED KINASE1 (*SnRK1*) signalling cascade is activated to combat low-energy often by stopping the plant growth involving a reduction in ribosomal protein synthesis, and a simultaneous accumulation of protective metabolites or defense compounds coupled with tuning of the metabolic processes in response to starvation (Baena-Gonzalez *et al.*, 2007, Wurzinger *et al.*, 2018). *SnRK1* signalling cascade involves interaction of *FLZ* gene family members with the kinase subunits. Expression of *FLZ* genes is both positively and negatively regulated by energy deficit as well as energy-rich conditions (Jamsheer and Laxmi, 2015). Starvation or low energy induces expression of *FLZ6, SnRK1* also induces *FLZ6* during energy starvation. Through repression of *SnRK1, FLZ6* promotes target of rapamycin or TOR signalling pathway that induces growth in favourable conditions. *FLZ6* mutants show inhibition and reduced seedling growth under favourable growth conditions due to enhanced *SnRK1* activity that confirms role of *FLZ6* in plant growth by regulating *SnRK1* signalling (Jamsheer *et al.*, 2018). *FLZ6* was found to be up-regulated under SHADE at latitude 67°2’N as compared to latitude 56°2’N, which may indicate that the trees in the northern latitudes are better adapted to shade conditions by withstanding the shade stress and continue to grow better compared to the southern populations.

### Shade responsive genes

Norway spruce is a shade tolerant conifer species. It is interesting that in our earlier work we reported that *SCRM2, NAC036* and *FLZ6* were responsible for the shade tolerant response in Norway spruce (Ranade *et al.*, 2019) and our current finding suggest that missense polymorphisms in these genes show a latitudinal cline in response to shade. *DRA2* and *PAR2* are the shade responsive genes that were up-regulated under SHADE, in the northern population.*DRA2* is involved in hypocotyl elongation, which along with *EXP3* (up-regulated in north) contributes to cell elongation, leading to elongation of the hypocotyls in the northern seedlings, which are the longest in response to SHADE among all the four latitudes. It may be concluded that although Norway spruce is shade tolerant, there is a difference in the hypocotyl elongation that follows a latitudinal cline.

### Plant defense

Up-regulation of higher number of defense-related genes was observed in the northern population under SHADE. Likewise, MapMan network indicates higher expression of genes at latitude 67°2’N that are included in the phenylpropanoid pathway involved in defense (Deng and Lu, 2017) (Supplementary Fig.S1-S3, Supplementary Material online). MYB3 represses phenylpropanoid biosynthesis and downregulation of *MYB3* (with cline in SNP) in the north indicates that phenylpropanoid pathway is enhanced at latitude 67°2’N. Enhanced secondary cell wall biosynthesis is indicated by up-regulation of *CESA4* in the northern latitude. Cell wall plays active role in plant immunity (Bacete *et al.*, 2018); lignin forms a barrier and restricts pathogen entry thus conferring disease resistance in plants (Lee *et al.*, 2019). Plant immunity specific genes *RPS2* and *LECRK-S.4*, were detected to be up-regulated in the northern trees; where *RPS2* also showed latitudinal cline in SNPs. We propose that northern trees might be better equipped for disease resistant mechanisms under shade as compared to the southern ones.

## Conclusions

Understanding adaptation to local climatic conditions is one of the crucial factors in the forest breeding and conservation. Our study on natural variation along a latitudinal cline has provided insights into the genomic basis for local adaptation to shade in Norway spruce - one of the most economically important conifer tree species in Sweden. Integration of knowledge on local adaption into forest tree breeding programs aims for a sustainable forestry. Climate change would lead to an increase in the mean temperatures especially in the northern latitudes, but it will have no effect or negligible effect on the light quality. Therefore, in the context of climate change, populations in the northern latitudes will continue to receive higher amount of FR light or low R:FR ratio which is equivalent to shade. Our results would contribute to efficient design of the programs for assisted migration as a solution to mitigate the mal-adaptation of present populations to their local conditions as a consequence of climatic change.

## Material and methods

Norway spruce seeds were collected from 70 unrelated trees from natural populations from four different latitudes across Sweden – Pellonhuhta (67°2’N, 22°1’E), Hallen (64°3’N, 14°6’E), Sör Amsberg (60°3’N, 15°2’E) and the southernmost population between latitude 56° and 58° (designated as 56°). These populations were selected based on previous studies in Norway spruce (Clapham *et al.*, 1998) in Sweden that reported an adaptive latitudinal cline in response to light quality. To ensure low consanguinity and to capture a representation of the population diversity, trees were collected at a distance of minimum 50 meters from each other. Cones were dried with warm air to force release the seeds. Sound seeds were separated from the empty seeds by flotation. The percentage of germination was obtained by germinating soaked seeds on paper discs on a warm bench with controlled humidity and temperature. The percentage of germination was of 98% on a batch of 200 seeds (five seeds per tree).

### Seed germination and light treatment

Stratified seeds (soaked in water at 4 °C overnight) were sown on water-saturated sterile vermiculite in growth boxes (Saveen Werner) and maintained at a constant temperature of 22°C in Percival (LED-30 Elite series) growth cabinets. 70 sound seeds per treatment and latitude (one seed per sampled tree) were germinated under two light treatments – Treatment A represented shade-like (SHADE) conditions having R:FR ratio of 0.2 and total light intensity of 36 μmol m^-2^s^-1^ (R, 6 μmol m^-2^s^-1^: FR, 30 μmol m^-2^s^-1^) and Treatment B (control) represented SUN-like (SUN) conditions having R:FR ratio equal to 1.2 and a total light intensity of 65 μmol m^-2^s^-1^ (R, 35 μmol m^-2^s^-1^: FR, 30 μmol m^-2^s^-1^). We considered only the R and FR light qualities in this experiment, as these are the two main responsible elements that plants use to determine shade conditions and respond accordingly. Plants sense the shade as decrease in R (or decrease in R:FR); R is primarily absorbed by the pigments and FR is reflected thereby decreasing the R:FR ratio. To mimic the natural conditions, FR intensity was kept constant under SHADE and SUN conditions and R intensity was lowered under SHADE. This experimental design was able to trigger the shade avoidance and the shade tolerance response in Scots pine and Norway spruce respectively, in our earlier work (Ranade *et al.*, 2019).

### Seedling morphological measurements

Hypocotyl was used as a model system to study the cline under the SHADE conditions in Norway spruce. Length of hypocotyl was scored at the mm precision for each seedling at the seedling developmental stage where hypocotyl was fully developed (Ranade *et al.*, 2013). Seedling morphology data under SHADE was converted into percentage of change with respect to SUN (100%). Analysis of variance (ANOVA) was applied to estimate the significance of the SHADE treatment effect. All statistical analysis was conducted using R software, version 3.5.2 (R Development Core Team, 2015).

### RNA sequencing (RNA-Seq)

RNA sequencing was performed with the samples from latitude 67° 2’ N and 56° 2’ N. Three biological replicates were prepared for each of the light treatments - A (SHADE) and B (SUN), for RNA extraction. The biological replicates were prepared by pooling three seedlings per sample to reduce variation between replicates and to increase the statistical power of the analysis (i.e. increased sensitivity to detect genes that were differentially expressed between conditions). Isolation of total RNA, RNA-Seq and pre-processing of RNA-Seq data was carried as described in our previous work (Ranade *et al.*, 2019). The RNA-Seq data was deposited to the ENA and is accessible under the accession number RJEB19683.

### Differential expression analysis

Statistical analysis of single-gene differential expression between the two latitudes in response to SHADE was determined using SUN as control. Analysis was performed using the Bioconductor (v3.3; (Gentleman *et al.*, 2004)) DESeq2 package (v1.12.0; (Love *et al.*, 2014)).

FDR adjusted *p-values* were used to assess significance; a common threshold of 1% was used throughout. For the data quality assessment (QA) and visualisation, the read counts were normalised using a variance stablishing transformation as implemented in DESeq2. The metabolic pathway maps were constructed with the DEGs using MapMan 3.5.1R2.

### Estimation of nucleotide variation following latitudinal cline

A total of 1654 individuals (unrelated parents) originating from different latitudes across Sweden, were included in this study for analysis of the SNP variation following a latitudinal cline. Details regarding the DNA extraction, exome capture, genotyping and SNP annotation have been previously described in Baison et al. (Baison *et al.*, 2019). For the current analysis, trees were divided into six populations, S1-S6. S1 comprised of 245 trees from latitudes 55-57, S2 with 213 trees from latitude 58, S3 with 187 trees from latitudes 59-60, S4 with 213 trees from latitudes 59-60, S5 with 573 trees from latitudes and S6 with 223 trees from latitudes 59-60. This categorisation of the population was done taking into consideration the fact that the quality of light differs latitude-wise from south to north; during the growth season (summer), the northern latitudes receive more amount of FR as compared to the southern ones.

The vcf file was filtered using settings; --min-alleles 2 --max-alleles 2 --maf 0.01 --remove-indels --minQ. 10 --max-missing 0.9. Allele frequencies, genotype frequencies and Hardy-Weinberg equilibrium (HWE) *p-values* were determined using SNPassoc statistical package (Gonzalez *et al.*, 2007). ANNOVA and Tukey’s *post-hoc* (Bonferroni *p-values)* was applied to determine the statistical significance in the difference in the allele and genotype frequencies across the populations included in the study. Genetic diversity among the six different populations was estimated using DnaSP 6 (Rozas *et al.*, 2017). Allele frequencies in each population were calculated and then regressed on population latitude. R^2^ of the linear regression were computed as the proportion of total variance of latitude explained by frequency of each marker (Berry and Kreitman, 1993).

## Acknowledgements

The authors thank the personal from Wallenberg greenhouse at SLU, Umeå for help with the handling of plants. We would also like to acknowledge the team at SLU’s försöksparker and Skogforsk for providing the seed collection for the four ecotypes involved in this study. We acknowledge Linghua Zhou for the comments with data representation. We acknowledge the UPSC bioinformatics facility (https://bioinfomatics.upsc.se) for technical support with regards to the RNA-Seq data pre-processing and analyses, and the support from Science for Life Laboratory (SciLifeLab), the Knut and Alice Wallenberg Foundation, the National Genomics Infrastructure funded by the Swedish Research Council, and Uppsala Multidisciplinary Centre for Advanced Computational Science for assistance with massively parallel sequencing and access to the UPPMAX computational infrastructure. We acknowledge Swedish Foundation for Strategic Research (SSF) for their support with reference to the exome capture project. This work was supported by the Kempe Foundation (JCK-1311) and Kungl. Skogs-och Lantbruksakademien (KSLA-H14-0150-ADA). We also acknowledge Swedish Research Council (VR) and Swedish Governmental Agency for Innovation Systems (VINNOVA) for their support.

## Author Contributions

SSR contributed with experiment performance, data collection, data analysis and interpretation, and manuscript writing. MRGG contributed with experimental design, data analysis and interpretation, and manuscript writing. Both authors read and approved the manuscript.

## Supporting Information

**All supporting tables and figures have been included in Supporting Information.pdf**

**Table S1-S2 Gene expression in response to SHADE in Norway Spruce**

**Table S3 Allele frequencies of missense mutations in the DEGs in response to SHADE in Norway spruce**

**Table S4 Allele and genotype frequencies of missense mutations in the nine candidate DEGs in response to SHADE in Norway spruce**

**Table S5 Tukey’s post-hoc test for genotype frequencies of missense mutations in the nine candidate DEGs in response to SHADE in Norway spruce**

**Table S6 Hardy Weinberg Equilibrium (HWE) for genotype frequencies of missense mutations in the nine candidate DEGs in response to SHADE in Norway spruce**

**Table S7 Allele frequencies of synonymous mutations in the DEGs in response to SHADE in Norway spruce**

**Fig.S1-S5 Metabolic pathway maps in response to SHADE in Norway Spruce**

**Fig.S6-S13 Cline with reference to variation in allele and genotype frequencies of SNPs in the candidate genes in Norway spruce populations across Sweden**

**Fig.S14-S22 Domains of the candidate genes with missense polymorphism sites in the Norway spruce**

