## Supplementary Material online for "Molecular signatures of local adaptation to light in Norway Spruce: role of lignin mediated immunity"

**Table S1 Genes up-regulated at latitude 67°2´N and down-regulated at latitude 56°2´N in response to SHADE in Norway spruce**

| Gene_ID | TAIR | Primary Gene Symbol | Gene Model Description | Expression | baseMean | log2FoldChange | lfcSE | stat | pvalue | padj |
|---|---|---|---|---|---|---|---|---|---|---|
| MA_93016g0010 | AT1G01490 | | Heavy metal transport/detoxification superfam | shade67>shade56 | 171.99156 | 19.31080501 | 3.3123768 | 5.8298938 | 5.55E-09 | 4.05E-06 |
| MA_10436915g0010 | AT1G01630 | | Sec14p-like phosphatidylinositol transfer famil | shade67>shade56 | 181.74323 | 0.890249942 | 0.2256632 | 3.9450378 | 7.98E-05 | 0.00785557 |
| MA_9801637g0010 | AT1G02080 | | transcription regulator;(source:Araport11) | shade67>shade56 | 213.33803 | 24.54497744 | 4.3990602 | 5.5795957 | 2.41E-08 | 1.52E-05 |
| MA_11144g0010 | AT1G02205 | ECERIFERUM 1 (CER1) | Expression of the CER1 gene associated with p | shade67>shade56 | 108.46666 | 2.068137632 | 0.5135865 | 4.0268533 | 5.65E-05 | 0.00616581 |
| MA_10107572g0010 | AT1G02800 | CELLULASE 2 (CEL2) | Encodes a protein with similarity to endo-1,4-b | shade67>shade56 | 139.88385 | 8.882139059 | 1.4882152 | 5.9683163 | 2.40E-09 | 1.91E-06 |
| MA_4984597g0010 | AT1G03230 | | Eukaryotic aspartyl protease family protein;(so | shade67>shade56 | 411.38787 | 21.9613859 | 4.8673892 | 4.5119436 | 6.42E-06 | 0.00109619 |
| MA_388691g0010 | AT1G03600 | (PSB27) | PSB27 is a chloroplast lumen localized protein | shade67>shade56 | 742.76598 | 0.648168009 | 0.1545924 | 4.192755 | 2.76E-05 | 0.00353428 |
| MA_45121g0010 | AT1G05670 | | Pentatricopeptide repeat (PPR-like) superfamil | shade67>shade56 | 184.58984 | 23.3255113 | 4.6756254 | 4.9887468 | 6.08E-07 | 0.00015401 |
| MA_5724086g0010 | AT1G10170 | NF-X-LIKE 1 (NFXL1) | Encodes AtNFXL1, a homologue of the putative | shade67>shade56 | 39.044387 | 21.84528163 | 5.5538932 | 3.9333277 | 8.38E-05 | 0.00803983 |
| MA_12361g0010 | AT1G10390 | DRACULA2 (DRA2) | DRA2 is a homolog of mammalian nucleoporin | shade67>shade56 | 95.148831 | 2.635868046 | 0.5869981 | 4.4904199 | 7.11E-06 | 0.00116898 |
| MA_640601g0010 | AT1G10640 | | Pectin lyase-like superfamily protein;(source:A | shade67>shade56 | 120.66502 | 1.686576144 | 0.3424242 | 4.9253994 | 8.42E-07 | 0.00020244 |
| MA_10430253g0010 | AT1G10740 | | alpha/beta-Hydrolases superfamily protein;(so | shade67>shade56 | 43.357063 | 19.20864968 | 4.2782117 | 4.4898782 | 7.13E-06 | 0.00116898 |
| MA_10432704g0020 | AT1G11290 | CHLORORESPIRATORY REDUCTI | Pentatricopeptide Repeat Protein containing th | shade67>shade56 | 388.93827 | 2.036870563 | 0.4773711 | 4.2668492 | 1.98E-05 | 0.00270169 |
| MA_500971g0010 | AT1G12050 | FUMARYLACETOACETATE HYDR | Encodes a fumarylacetoacetase that converts | shade67>shade56 | 137.11908 | 30 | 6.0065614 | 4.9945381 | 5.90E-07 | 0.00015401 |
| MA_10435025g0010 | AT1G13520 | | hypothetical protein (DUF1262);(source:Arapo | shade67>shade56 | 139.92554 | 2.605127449 | 0.6168562 | 4.223233 | 2.41E-05 | 0.00318496 |
| MA_15382g0010 | AT1G13570 | | F-box/RNI-like superfamily protein;(source:Ara | shade67>shade56 | 63.129848 | 19.75701806 | 4.5405589 | 4.3512304 | 1.35E-05 | 0.00200323 |
| MA_179350g0020 | AT1G23740 | ALKENAL/ONE OXIDOREDUCTAS | AOR is an alkenal/one oxidoreductase that act | shade67>shade56 | 467.58936 | 25.98001962 | 4.9570437 | 5.2410309 | 1.60E-07 | 7.51E-05 |
| MA_470965g0010 | AT1G25440 | B-BOX DOMAIN PROTEIN 15 (BE | B-box type zinc finger protein with CCT domair | shade67>shade56 | 455.31386 | 1.121153225 | 0.2626287 | 4.2689661 | 1.96E-05 | 0.00268966 |
| MA_89572g0010 | AT1G26760 | SET DOMAIN PROTEIN 35 (SDG3 | SET domain protein 35;(source:Araport11) | shade67>shade56 | 42.735581 | 20.65022444 | 4.4404534 | 4.6504765 | 3.31E-06 | 0.00060152 |
| MA_129800g0010 | AT1G27170 | | transmembrane receptors / ATP binding protei | shade67>shade56 | 61.243749 | 23.9648196 | 5.8925615 | 4.0669613 | 4.76E-05 | 0.00538972 |
| MA_190687g0010 | AT1G28570 | | SGNH hydrolase-type esterase superfamily pro | shade67>shade56 | 221.37196 | 19.19121621 | 4.3179928 | 4.4444762 | 8.81E-06 | 0.00140178 |
| MA_78595g0010 | AT1G34300 | | lectin protein kinase family protein;(source:Ara | shade67>shade56 | 236.56516 | 26.19644897 | 4.5192601 | 5.7966234 | 6.77E-09 | 4.69E-06 |
| MA_9827161g0010 | AT1G50510 | | indigoidine synthase A family protein;(source:A | shade67>shade56 | 350.65976 | 24.79895514 | 5.8899002 | 4.2104203 | 2.55E-05 | 0.00332946 |
| MA_27434g0010 | AT1G55790 | | ferredoxin-fold anticodon-binding domain prot | shade67>shade56 | 35.061281 | 22.4663389 | 5.5747433 | 4.0300222 | 5.58E-05 | 0.00613067 |
| MA_119176g0010 | AT1G56145 | | Leucine-rich repeat transmembrane protein kir | shade67>shade56 | 212.29721 | 11.61665618 | 2.5818513 | 4.4993514 | 6.82E-06 | 0.00114018 |
| MA_242575g0010 | AT1G56720 | | Protein kinase superfamily protein;(source:Ara | shade67>shade56 | 144.14445 | 23.65612586 | 5.7870066 | 4.0878 | 4.35E-05 | 0.00502182 |
| MA_650284g0010 | AT1G59740 | NRT1/ PTR FAMILY 4.3 (NPF4.3) | Major facilitator superfamily protein;(source:A | shade67>shade56 | 117.29519 | 24.4485448 | 4.7248031 | 5.1745109 | 2.29E-07 | 0.00010074 |
| MA_1734g0010 | AT1G65450 | GLAUCE (GLC) | Contains dual transcription units and alternativ | shade67>shade56 | 67.713142 | 2.31918862 | 0.5679846 | 4.0831891 | 4.44E-05 | 0.00509479 |
| MA_52897g0010 | AT1G66920 | | Protein kinase superfamily protein;(source:Ara | shade67>shade56 | 214.04324 | 22.16541059 | 4.3348556 | 5.1132985 | 3.17E-07 | 0.0001264 |
| MA_9239567g0010 | AT1G69930 | GLUTATHIONE S-TRANSFERASE | Encodes glutathione transferase belonging to t | shade67>shade56 | 48.887639 | 6.951129764 | 1.4925065 | 4.6573532 | 3.20E-06 | 0.00058802 |
| MA_129851g0010 | AT1G72470 | EXOCYST SUBUNIT EXO70 FAMI | A member of EXO70 gene family, putative exoc | shade67>shade56 | 68.886724 | 23.1219094 | 5.8916095 | 3.9245489 | 8.69E-05 | 0.00824038 |
| MA_553396g0010 | AT1G74310 | HEAT SHOCK PROTEIN 101 (HSP | Encodes ClpB1, which belongs to the Casein lyt | shade67>shade56 | 229.3189 | 24.41459867 | 4.582808 | 5.3274321 | 9.96E-08 | 5.34E-05 |
| MA_88684g0010 | AT1G74460 | | GDSL-motif esterase/acyltransferase/lipase. Er | shade67>shade56 | 198.59424 | 0.780591597 | 0.1767749 | 4.4157374 | 1.01E-05 | 0.00158375 |
| MA_180495g0010 | AT1G78060 | | Glycosyl hydrolase family protein;(source:Arap | shade67>shade56 | 104.55114 | 1.492501596 | 0.3435133 | 4.3448148 | 1.39E-05 | 0.00205549 |
| MA_108157g0010 | AT1G78800 | | UDP-Glycosyltransferase superfamily protein;(s | shade67>shade56 | 288.92213 | 24.91890254 | 4.5419299 | 5.4864128 | 4.10E-08 | 2.44E-05 |
| MA_10348903g0010 | AT1G78895 | | Reticulon family protein;(source:Araport11) | shade67>shade56 | 95.841286 | 17.97643928 | 3.2894734 | 5.464838 | 4.63E-08 | 2.72E-05 |
| MA_10429874g0010 | AT1G80160 | GLYOXYLASE I 7 (GLYI7) | Vicinal oxygen chelate (VOC) superfamily mem | shade67>shade56 | 161.27715 | 23.12495978 | 5.8609333 | 3.9456105 | 7.96E-05 | 0.00785557 |
| MA_395055g0010 | AT1G80820 | CINNAMOYL COA REDUCTASE (C | Encodes an cinnamoyl CoA reductase isoform. | shade67>shade56 | 40.899254 | 30 | 6.0094577 | 4.992131 | 5.97E-07 | 0.00015401 |
| MA_19208g0030 | AT2G01850 | ENDOXYLOGLUCAN TRANSFERA | EXGT-A3 has homology to xyloglucan endotran | shade67>shade56 | 124.3984 | 22.68203697 | 4.60999 | 4.9201922 | 8.65E-07 | 0.00020673 |
| MA_16619g0010 | AT2G02450 | LONG VEGETATIVE PHASE 1 (LO | NAC domain containing protein 35;(source:Ara | shade67>shade56 | 76.088784 | 22.14299267 | 4.3836582 | 5.0512589 | 4.39E-07 | 0.00015401 |
| MA_78676g0020 | AT2G13600 | SLOW GROWTH 2 (SLO2) | Encodes a pentatricopeptide repeat protein be | shade67>shade56 | 196.9592 | 21.58842311 | 4.6065824 | 4.6864294 | 2.78E-06 | 0.0005233 |
| MA_101849g0010 | AT2G17040 | NAC DOMAIN CONTAINING PRO | Member of the NAC transcription factor family | shade67>shade56 | 461.71542 | 26.84616555 | 5.2685729 | 5.0955289 | 3.48E-07 | 0.00013233 |
| MA_7493784g0010 | AT2G21730 | CINNAMYL ALCOHOL DEHYDRO | cinnamyl alcohol dehydrogenase homolog 2;(s | shade67>shade56 | 143.36126 | 22.94004279 | 4.4972198 | 5.1009388 | 3.38E-07 | 0.00013122 |
| MA_455g0010 | AT2G22250 | ASPARTATE AMINOTRANSFERAS | Encodes a prokaryotic-type plastidic aspartate | shade67>shade56 | 154.31616 | 21.73531843 | 4.5262512 | 4.8020575 | 1.57E-06 | 0.00033066 |
| MA_7330598g0010 | AT2G22590 | | UDP-Glycosyltransferase superfamily protein;(s | shade67>shade56 | 341.19687 | 20.19612958 | 4.4945833 | 4.4934376 | 7.01E-06 | 0.00116312 |
| MA_177197g0020 | AT2G24130 | | Leucine-rich receptor-like protein kinase family | shade67>shade56 | 107.67522 | 26.02330665 | 5.8956843 | 4.4139586 | 1.01E-05 | 0.00158504 |
| MA_10427649g0020 | AT2G26700 | PINOID2 (PID2) | Member of AGC VIIIa Kinase gene family. Enco | shade67>shade56 | 145.97226 | 3.016886255 | 0.6021253 | 5.0103961 | 5.43E-07 | 0.00015401 |
| MA_20782g0010 | AT2G27610 | | Tetratricopeptide repeat (TPR)-like superfamily | shade67>shade56 | 185.7094 | 1.647532558 | 0.361988 | 4.5513464 | 5.33E-06 | 0.00093994 |
| MA_906876g0010 | AT2G29380 | HIGHLY ABA-INDUCED PP2C GEN | highly ABA-induced PP2C protein 3;(source:Ara | shade67>shade56 | 101.08365 | 2.557108004 | 0.6497897 | 3.9352856 | 8.31E-05 | 0.0080187 |
| MA_116552g0010 | AT2G32010 | CVP2 LIKE 1 (CVL1) | Encodes an inositol polyphosphate 5?-phospha | shade67>shade56 | 203.68622 | 1.477916508 | 0.3769251 | 3.9209823 | 8.82E-05 | 0.00831233 |
| MA_161036g0010 | AT2G33150 | PEROXISOMAL 3-KETOACYL-COA | Encodes an organellar (peroxisome, glyoxysom | shade67>shade56 | 62.413835 | 22.23258589 | 5.6952758 | 3.9036891 | 9.47E-05 | 0.008787 |
| MA_10432854g0010 | AT2G34930 | | disease resistance family protein / LRR family p | shade67>shade56 | 246.28107 | 19.21591367 | 4.4020052 | 4.3652638 | 1.27E-05 | 0.00191227 |
| MA_10434829g0020 | AT2G35730 | | Heavy metal transport/detoxification superfam | shade67>shade56 | 257.51116 | 1.652742878 | 0.419015 | 3.9443524 | 8.00E-05 | 0.00785557 |
| MA_26811g0010 | AT2G36750 | UDP-GLUCOSYL TRANSFERASE 7 | UDP-glucosyl transferase 73C1;(source:Araport | shade67>shade56 | 642.45689 | 20.92464441 | 3.062855 | 6.831745 | 8.39E-12 | 2.09E-08 |
| MA_8644998g0010 | AT2G36830 | GAMMA TONOPLAST INTRINSIC | Encodes a tonoplast intrinsic protein, which fu | shade67>shade56 | 57.502335 | 22.85053915 | 5.8929248 | 3.8776228 | 0.00010548 | 0.00957963 |
| MA_15920g0010 | AT2G38300 | | myb-like HTH transcriptional regulator family p | shade67>shade56 | 61.559852 | 20.26489734 | 3.19972 | 6.3333346 | 2.40E-10 | 2.99E-07 |
| MA_41286g0020 | AT2G39210 | | Major facilitator superfamily protein;(source:A | shade67>shade56 | 265.38725 | 9.774556004 | 2.4734683 | 3.9517612 | 7.76E-05 | 0.00778006 |
| MA_10429137g0020 | AT2G44460 | BETA GLUCOSIDASE 28 (BGLU28 | beta glucosidase 28;(source:Araport11) | shade67>shade56 | 610.9023 | 11.28373972 | 2.8598257 | 3.945604 | 7.96E-05 | 0.00785557 |
| MA_9544285g0010 | AT2G45660 | AGAMOUS-LIKE 20 (AGL20) | Controls flowering and is required for CO to pr | shade67>shade56 | 59.14217 | 24.48104131 | 5.4303222 | 4.5082115 | 6.54E-06 | 0.00110231 |
| MA_9957507g0010 | AT2G45670 | LYSOPHOSPHATIDYLETHANOLAI | Encodes an acyl-CoA: lysophosphatidylethanol | shade67>shade56 | 77.389898 | 1.950904293 | 0.3984553 | 4.8961686 | 9.77E-07 | 0.000226 |
| MA_10431662g0010 | AT3G01160 | | pre-rRNA-processing ESF1-like protein;(source: | shade67>shade56 | 39.915146 | 21.71071847 | 5.5445025 | 3.9157199 | 9.01E-05 | 0.00843675 |
| MA_187159g0010 | AT3G01190 | | Peroxidase superfamily protein;(source:Arapor | shade67>shade56 | 72.076916 | 23.3149957 | 5.5560126 | 4.196354 | 2.71E-05 | 0.00349982 |
| MA_9076332g0010 | AT3G04140 | | Ankyrin repeat family protein;(source:Araport1 | shade67>shade56 | 16.847116 | 3.1738832 | 0.8031452 | 3.9518174 | 7.76E-05 | 0.00778006 |
| MA_8035959g0010 | AT3G04720 | PATHOGENESIS-RELATED 4 (PR4 | Encodes a protein similar to the antifungal chit | shade67>shade56 | 137.62157 | 25.66154289 | 5.4483365 | 4.7099776 | 2.48E-06 | 0.00048095 |
| MA_101858g0010 | AT3G11980 | MALE STERILITY 2 (MS2) | Similar to fatty acid reductases. | shade67>shade56 | 21.755398 | 30 | 6.0094222 | 4.9921605 | 5.97E-07 | 0.00015401 |
| MA_9825276g0010 | AT3G12390 | | Nascent polypeptide-associated complex (NAC | shade67>shade56 | 64.092653 | 25.42688374 | 5.896436 | 4.3122462 | 1.62E-05 | 0.00230278 |
| MA_85286g0010 | AT3G13080 | ATP-BINDING CASSETTE C3 (ABC | encodes an ATP-dependent MRP-like ABC trans | shade67>shade56 | 68.252939 | 25.17279858 | 5.8945496 | 4.2705211 | 1.95E-05 | 0.00267964 |
| MA_10432070g0010 | AT3G17611 | RHOMBOID-LIKE PROTEIN 14 (R | RHOMBOID-like protein 14;(source:Araport11) | shade67>shade56 | 308.39347 | 9.553088024 | 1.4137652 | 6.7571957 | 1.41E-11 | 3.31E-08 |
| MA_10208460g0010 | AT3G18090 | NUCLEAR RNA POLYMERASE D2 | Encodes a subunit of RNA polymerase IV (aka P | shade67>shade56 | 358.56828 | 25.8970636 | 5.8899104 | 4.3968519 | 1.10E-05 | 0.00170264 |
| MA_9208522g0010 | AT3G19640 | MAGNESIUM TRANSPORTER 4 ( | Transmembrane magnesium transporter. One | shade67>shade56 | 73.279952 | 30 | 5.5599977 | 5.3956857 | 6.83E-08 | 3.85E-05 |
| MA_10342550g0010 | AT3G21000 | | Gag-Pol-related retrotransposon family protein | shade67>shade56 | 43.666787 | 30 | 6.0073801 | 4.9938575 | 5.92E-07 | 0.00015401 |
| MA_226036g0010 | AT3G27970 | | Exonuclease family protein;(source:Araport11) | shade67>shade56 | 173.81697 | 10.14610409 | 1.5841105 | 6.4049221 | 1.50E-10 | 2.20E-07 |
| MA_9736190g0010 | AT3G43660 | | The gene encodes a putative nodulin-like21 pr | shade67>shade56 | 72.609617 | 7.897492333 | 1.4719195 | 5.3654377 | 8.08E-08 | 4.50E-05 |
| MA_218285g0010 | AT3G44050 | | P-loop containing nucleoside triphosphate hyd | shade67>shade56 | 82.801515 | 23.49464132 | 5.6822669 | 4.1347303 | 3.55E-05 | 0.00429705 |
| MA_301190g0010 | AT3G47570 | | Leucine-rich repeat protein kinase family prote | shade67>shade56 | 247.21192 | 23.80674082 | 5.8189401 | 4.0912504 | 4.29E-05 | 0.00497479 |
| MA_551352g0010 | AT3G50740 | UDP-GLUCOSYL TRANSFERASE 7 | UGT72E1 is an UDPG:coniferyl alcohol glucosyl | shade67>shade56 | 44.297926 | 9.692525974 | 1.8236437 | 5.3149229 | 1.07E-07 | 5.51E-05 |
| MA_9132531g0010 | AT3G51520 | DIACYLGLYCEROL ACYLTRANSFE | Encodes a functional acyl-CoA:diacylglycerol ac | shade67>shade56 | 100.09837 | 1.27190695 | 0.3212249 | 3.9595523 | 7.51E-05 | 0.00761091 |
| MA_10430424g0010 | AT3G54420 | HOMOLOG OF CARROT EP3-3 CI | encodes an EP3 chitinase that is expressed dur | shade67>shade56 | 69.439419 | 1.904793455 | 0.4851331 | 3.9263317 | 8.63E-05 | 0.00822125 |
| MA_96570g0010 | AT3G58850 | PHY RAPIDLY REGULATED 2 (PAI | Encodes PHYTOCHROME RAPIDLY REGULATED2 | shade67>shade56 | 151.15644 | 25.46565052 | 4.6671481 | 5.4563622 | 4.86E-08 | 2.82E-05 |
| MA_227006g0010 | AT3G59630 | | diphthamide synthesis DPH2 family protein;(sc | shade67>shade56 | 285.32735 | 23.04775626 | 3.6101531 | 6.3841493 | 1.72E-10 | 2.43E-07 |
| MA_10435788g0030 | AT3G60850 | | hypothetical protein;(source:Araport11) | shade67>shade56 | 44.862537 | 22.27784827 | 4.2893835 | 5.1937179 | 2.06E-07 | 9.18E-05 |
| MA_129478g0010 | AT3G61460 | BRASSINOSTEROID-RESPONSIVE | Encodes a novel ring finger protein and forms a | shade67>shade56 | 269.27653 | 25.02047576 | 4.7458892 | 5.2720312 | 1.35E-07 | 6.64E-05 |
| MA_14234g0010 | AT3G62550 | | Adenine nucleotide alpha hydrolases-like supe | shade67>shade56 | 91.451935 | 2.124069347 | 0.5318995 | 3.9933657 | 6.51E-05 | 0.00684088 |
| MA_90455g0010 | AT4G00290 | (MSL1) | Encodes a non-selective mechanosensitive ion | shade67>shade56 | 78.092407 | 23.01422025 | 4.5550231 | 5.0524925 | 4.36E-07 | 0.00015401 |
| MA_62903g0010 | AT4G00880 | SMALL AUXIN UPREGULATED RI | SAUR-like auxin-responsive protein family;(sou | shade67>shade56 | 186.56969 | 30 | 6.0096019 | 4.9920112 | 5.98E-07 | 0.00015401 |
| MA_155037g0010 | AT4G01130 | | GDSL-motif esterase/acyltransferase/lipase. Er | shade67>shade56 | 73.867381 | 30 | 6.0075272 | 4.9937352 | 5.92E-07 | 0.00015401 |
| MA_10435309g0010 | AT4G02780 | GA REQUIRING 1 (GA1) | Catalyzes the conversion of geranylgeranyl pyr | shade67>shade56 | 1484.8314 | 21.66426694 | 5.0131962 | 4.3214481 | 1.55E-05 | 0.00223228 |

| | | | | | | | | | | |
|---|---|---|---|---|---|---|---|---|---|---|
| MA_10427046g0010 | AT4G13430 | ISOPROPYL MALATE ISOMERASE | Encodes a methylthioalkylmalate isomerase in | shade67>shade56 | 98.540445 | 1.006114198 | 0.2302775 | 4.3691388 | 1.25E-05 | 0.00188536 |
| MA_4231108g0010 | AT4G16970 | | Protein kinase superfamily protein;(source:Ara | shade67>shade56 | 132.69181 | 23.24712591 | 5.8316618 | 3.9863638 | 6.71E-05 | 0.00702839 |
| MA_10426824g0010 | AT4G17785 | MYB DOMAIN PROTEIN 39 (MYB | Encodes a putative transcription factor (MYB39 | shade67>shade56 | 74.263519 | 2.386550787 | 0.5503552 | 4.3363827 | 1.45E-05 | 0.0021182 |
| MA_10433704g0010 | AT4G18550 | DAD1-LIKE SEEDING ESTABLISHM | DSEL is cytosolic acylhydrolase that shows pref | shade67>shade56 | 190.61376 | 2.085062526 | 0.3396734 | 6.1384327 | 8.33E-10 | 8.82E-07 |
| MA_10428179g0010 | AT4G19450 | | Major facilitator superfamily protein;(source:A | shade67>shade56 | 415.5138 | 2.267466224 | 0.5375758 | 4.2179469 | 2.47E-05 | 0.00323386 |
| MA_12346g0010 | AT4G23290 | CYSTEINE-RICH RLK (RECEPTOR- | Encodes a cysteine-rich receptor-like protein ki | shade67>shade56 | 22.587358 | 21.92887916 | 3.1855846 | 6.883785 | 5.83E-12 | 1.64E-08 |
| MA_178762g0010 | AT4G23440 | | Disease resistance protein (TIR-NBS class);(sou | shade67>shade56 | 94.75764 | 30 | 5.8085164 | 5.16483 | 2.41E-07 | 0.000105 |
| MA_498754g0010 | AT4G24130 | | DUF538 family protein (Protein of unknown fu | shade67>shade56 | 45.012723 | 2.359532685 | 0.5523136 | 4.2720886 | 1.94E-05 | 0.00266954 |
| MA_475302g0010 | AT4G26090 | RESISTANT TO P. SYRINGAE 2 (R | Encodes a plasma membrane protein with leu | shade67>shade56 | 69.501895 | 18.84260247 | 3.8365974 | 4.9112795 | 9.05E-07 | 0.00021201 |
| MA_5382714g0010 | AT4G26330 | UNFERTILIZED EMBRYO SAC 17 | Subtilisin-like serine endopeptidase family prot | shade67>shade56 | 217.52282 | 22.4460546 | 3.6951903 | 6.0743975 | 1.24E-09 | 1.20E-06 |
| MA_9399920g0010 | AT4G26740 | SEED GENE 1 (ATS1) | Encodes caleosin, a 27-kDa protein found with | shade67>shade56 | 122.09961 | 9.820074023 | 2.017926 | 4.8664193 | 1.14E-06 | 0.00025446 |
| MA_551848g0020 | AT4G26970 | ACONITASE 2 (ACO2) | Encodes an aconitase that can catalyze the cor | shade67>shade56 | 60.353095 | 24.89660816 | 5.8941381 | 4.2239608 | 2.40E-05 | 0.00318463 |
| MA_7354451g0010 | AT4G28250 | EXPANSIN B3 (EXPB3) | putative beta-expansin/allergen protein. Nami | shade67>shade56 | 143.08438 | 1.612388608 | 0.3690315 | 4.3692442 | 1.25E-05 | 0.00188536 |
| MA_9930140g0010 | AT4G30170 | | Peroxidase family protein;(source:Araport11) | shade67>shade56 | 62.037677 | 21.58697676 | 4.2647119 | 5.0617667 | 4.15E-07 | 0.00015287 |
| MA_36985g0020 | AT4G33300 | ADR1-LIKE 1 (ADR1-L1) | Encodes a member of the ADR1 family nucleot | shade67>shade56 | 45.027778 | 8.91149957 | 2.2191578 | 4.0157124 | 5.93E-05 | 0.00639851 |
| MA_501572g0010 | AT4G33720 | (ATCAPE3) | CAP (Cysteine-rich secretory proteins, Antigen | shade67>shade56 | 178.30396 | 19.21367128 | 4.4740714 | 4.294449 | 1.75E-05 | 0.00244747 |
| MA_15237g0010 | AT4G34100 | ECERIFERUM 9 (CER9) | Encodes a protein involved in cuticular wax bio | shade67>shade56 | 34.446299 | 20.08009195 | 4.3147584 | 4.6538161 | 3.26E-06 | 0.0005944 |
| MA_10427611g0010 | AT4G38190 | CELLULOSE SYNTHASE LIKE D4 ( | encodes a gene similar to cellulose synthase | shade67>shade56 | 70.581856 | 9.382576657 | 1.4742493 | 6.3643081 | 1.96E-10 | 2.62E-07 |
| MA_27312g0010 | AT5G02900 | CYTOCHROME P450, FAMILY 96 | member of CYP96A | shade67>shade56 | 41.060459 | 24.59198292 | 5.1543598 | 4.7711033 | 1.83E-06 | 0.00037641 |
| MA_10425930g0010 | AT5G04895 | ABA OVERLY SENSITIVE 6 (ABO6 | DEA(D/H)-box RNA helicase family protein;(sou | shade67>shade56 | 167.21642 | 30 | 6.0068133 | 4.9943287 | 5.90E-07 | 0.00015401 |
| MA_10238446g0010 | AT5G05170 | CONSTITUTIVE EXPRESSION OF | Encodes a cellulose synthase isomer. CESA3 mu | shade67>shade56 | 166.18441 | 24.65722666 | 5.8904557 | 4.1859625 | 2.84E-05 | 0.00363066 |
| MA_9328561g0010 | AT5G05850 | PLANT INTRACELLULAR RAS GRO | Encodes PIRL1, a member of the Plant Intracell | shade67>shade56 | 56.744221 | 30 | 6.0076142 | 4.9936629 | 5.92E-07 | 0.00015401 |
| MA_422924g0010 | AT5G06360 | | Ribosomal protein S8e family protein;(source:A | shade67>shade56 | 99.352722 | 19.00542484 | 4.5153843 | 4.2090382 | 2.56E-05 | 0.00333958 |
| MA_9786598g0010 | AT5G07580 | (ERF106) | encodes a member of the ERF (ethylene respor | shade67>shade56 | 57.692637 | 21.18347183 | 4.4742683 | 4.7345109 | 2.20E-06 | 0.00043835 |
| MA_6332848g0010 | AT5G08570 | | Pyruvate kinase family protein;(source:Araport | shade67>shade56 | 61.577376 | 7.314099244 | 1.5687699 | 4.6623147 | 3.13E-06 | 0.00057784 |
| MA_10432569g0010 | AT5G09970 | CYTOCHROME P450, FAMILY 78 | member of CYP78A | shade67>shade56 | 399.65958 | 10.03243663 | 1.4394621 | 6.9695734 | 3.18E-12 | 9.61E-09 |
| MA_8066361g0010 | AT5G17540 | | HXXXD-type acyl-transferase family protein;(so | shade67>shade56 | 70.935852 | 23.08690862 | 5.8129366 | 3.971643 | 7.14E-05 | 0.00729666 |
| MA_10192245g0010 | AT5G17920 | METHIONINE SYNTHESIS 1 (ATM | Encodes a cytosolic cobalamin-independent m | shade67>shade56 | 120.13048 | 23.46768982 | 5.8905826 | 3.9839336 | 6.78E-05 | 0.00708316 |
| MA_10428851g0010 | AT5G18980 | | ARM repeat superfamily protein;(source:Arapo | shade67>shade56 | 300.34056 | 19.39280001 | 3.3221509 | 5.8374231 | 5.30E-09 | 3.94E-06 |
| MA_960556g0010 | AT5G19930 | PLASMA MEMBRANE GLUCOSE- | PGR is putative plasma membrane glucose- res | shade67>shade56 | 111.16564 | 24.59536423 | 4.3200391 | 5.6933199 | 1.25E-08 | 8.24E-06 |
| MA_121630g0010 | AT5G20480 | EF-TU RECEPTOR (EFR) | Encodes a predicted leucine-rich repeat recept | shade67>shade56 | 26.649066 | 22.00904044 | 5.479014 | 4.016971 | 5.90E-05 | 0.00638073 |
| MA_41345g0010 | AT5G20950 | (BGLC1) | Encodes a beta-glucosidase involved in xyloglu | shade67>shade56 | 64.774986 | 9.855500114 | 1.8217342 | 5.409955 | 6.30E-08 | 3.61E-05 |
| MA_10437273g0010 | AT5G24080 | | Protein kinase superfamily protein;(source:Ara | shade67>shade56 | 128.58433 | 25.06420688 | 5.8916771 | 4.2541718 | 2.10E-05 | 0.00281904 |
| MA_132981g0010 | AT5G24760 | | GroES-like zinc-binding dehydrogenase family | shade67>shade56 | 11339.414 | 1.617442195 | 0.4114815 | 3.9307775 | 8.47E-05 | 0.00810721 |
| MA_10434521g0010 | AT5G27100 | GLUTAMATE RECEPTOR 2.1 (GL | member of Putative ligand-gated ion channel s | shade67>shade56 | 195.80201 | 27.11057913 | 5.3083363 | 5.1071706 | 3.27E-07 | 0.00012934 |
| MA_10307058g0010 | AT5G33406 | | hAT dimerization domain-containing protein / | shade67>shade56 | 104.57168 | 23.22641523 | 4.4124719 | 5.2638103 | 1.41E-07 | 6.79E-05 |
| MA_8760260g0010 | AT5G36930 | | Disease resistance protein (TIR-NBS-LRR class) | shade67>shade56 | 35.361848 | 23.57751423 | 5.8966228 | 3.9984776 | 6.38E-05 | 0.00672822 |
| MA_494813g0010 | AT5G39320 | UDP-GLUCOSE DEHYDROGENAS | UDP-glucose 6-dehydrogenase family protein;( | shade67>shade56 | 118.97449 | 21.89497984 | 4.3769355 | 5.0023537 | 5.66E-07 | 0.00015401 |
| MA_183130g0020 | AT5G44030 | CELLULOSE SYNTHASE A4 (CESA | Encodes a cellulose synthase involved in secon | shade67>shade56 | 130.7649 | 18.39774898 | 4.2813283 | 4.2972058 | 1.73E-05 | 0.00244001 |
| MA_10428737g0010 | AT5G44380 | (ATBBE28) | FAD-binding Berberine family protein;(source: | shade67>shade56 | 93.64181 | 23.02795435 | 4.4935545 | 5.1246635 | 2.98E-07 | 0.00012247 |
| MA_986218g0010 | AT5G44700 | GASSHO 2 (GSO2) | Encodes GASSHO2 (GSO2), a putative leucine-r | shade67>shade56 | 165.23896 | 21.02946968 | 4.4369235 | 4.7396512 | 2.14E-06 | 0.0004294 |
| MA_10319538g0010 | AT5G45370 | USUALLY MULTIPLE ACIDS MOV | nodulin MtN21-like transporter family protein | shade67>shade56 | 17.447841 | 14.92039355 | 3.6351066 | 4.1045272 | 4.05E-05 | 0.00477606 |
| MA_90634g0010 | AT5G46610 | | aluminum activated malate transporter family | shade67>shade56 | 64.971654 | 7.108339214 | 1.802015 | 3.9446614 | 7.99E-05 | 0.00785557 |
| MA_83446g0020 | AT5G50400 | PURPLE ACID PHOSPHATASE 27 | purple acid phosphatase 27;(source:Araport11 | shade67>shade56 | 317.71611 | 21.33455177 | 3.4619624 | 6.1625602 | 7.16E-10 | 7.77E-07 |
| MA_321195g0010 | AT5G51780 | | basic helix-loop-helix (bHLH) DNA-binding supe | shade67>shade56 | 168.47283 | 12.79658274 | 2.427048 | 5.2724885 | 1.35E-07 | 6.64E-05 |
| MA_39500g0010 | AT5G53470 | ACYL-COA BINDING PROTEIN 1 ( | Encodes an acyl-CoA binding protein that is loc | shade67>shade56 | 109.56587 | 1.789931901 | 0.3968894 | 4.509901 | 6.49E-06 | 0.00109794 |
| MA_766568g0010 | AT5G54930 | METABOLIC NETWORK MODULA | AT hook motif-containing protein;(source:Arap | shade67>shade56 | 81.183942 | 10.29709703 | 2.5140508 | 4.0958189 | 4.21E-05 | 0.00493173 |
| MA_10426326g0020 | AT5G55090 | MITOGEN-ACTIVATED PROTEIN | member of MEKK subfamily | shade67>shade56 | 233.8183 | 9.34552367 | 1.3981422 | 6.6842442 | 2.32E-11 | 4.82E-08 |
| MA_10428846g0020 | AT5G55250 | IAA CARBOXYLMETHYLTRANSFE | Encodes an enzyme which specifically converts | shade67>shade56 | 48.710255 | 17.97497551 | 3.8356595 | 4.6862803 | 2.78E-06 | 0.0005233 |
| MA_10426320g0010 | AT5G56960 | | basic helix-loop-helix (bHLH) DNA-binding fami | shade67>shade56 | 94.262969 | 23.53511908 | 5.4073855 | 4.3524027 | 1.35E-05 | 0.00199954 |
| MA_10434319g0010 | AT5G58360 | OVATE FAMILY PROTEIN 3 (OFP | ovate family protein 3;(source:Araport11) | shade67>shade56 | 177.21754 | 1.223426997 | 0.2913975 | 4.1984811 | 2.69E-05 | 0.00347772 |
| MA_875703g0010 | AT5G60700 | | glycosyltransferase family protein 2;(source:Ar | shade67>shade56 | 566.80733 | 2.251919099 | 0.4671589 | 4.8204559 | 1.43E-06 | 0.00030461 |
| MA_92381g0010 | AT5G64140 | RIBOSOMAL PROTEIN S28 (RPS2 | Encodes a putative ribosomal protein S28. | shade67>shade56 | 121.24359 | 23.74769686 | 5.7573079 | 4.1247919 | 3.71E-05 | 0.00443618 |
| MA_261256g0010 | AT5G64570 | BETA-D-XYLOSIDASE 4 (XYL4) | Encodes a beta-d-xylosidase that belongs to fa | shade67>shade56 | 41.297439 | 22.53010886 | 5.4595113 | 4.1267629 | 3.68E-05 | 0.00442333 |

**Table S2 Genes down-regulated at latitude 67$^{o}$2´N and up-regulated at latitude 56$^{o}$2´N in response to SHADE in Norway spruce**

| Gene_ID | TAIR | Primary Gene Symbol | Gene Model Description | Expression | baseMean | log2FoldChange | lfcSE | stat | pvalue | padj |
|---|---|---|---|---|---|---|---|---|---|---|
| MA_10325260g0010 | AT1G04580 | ALDEHYDE OXIDASE 4 (AO4) | Encodes aldehyde oxidase AAO4 preferentially | shade67<shade56 | 81.315393 | -22.88013074 | 5.8918542 | -3.88335 | 0.00010303 | 0.00941731 |
| MA_536645g0010 | AT1G05200 | GLUTAMATE RECEPTOR 3.4 (GL | Encodes a putative glutamate receptor GLR3 w | shade67<shade56 | 272.51679 | -25.55209379 | 4.2803869 | -5.969576 | 2.38E-09 | 1.91E-06 |
| MA_355745g0010 | AT1G05270 | | TraB family protein;(source:Araport11) | shade67<shade56 | 214.99011 | -12.54145848 | 2.4921623 | -5.03236 | 4.84E-07 | 0.00015401 |
| MA_213682g0010 | AT1G06640 | | encodes a protein whose sequence is similar to | shade67<shade56 | 210.29368 | -23.77830423 | 5.8901633 | -4.036952 | 5.42E-05 | 0.00599576 |
| MA_10427341g0010 | AT1G07650 | | Leucine-rich repeat transmembrane protein ki | shade67<shade56 | 2798.077 | -3.098507079 | 0.7826282 | -3.959105 | 7.52E-05 | 0.00761091 |
| MA_496233g0010 | AT1G11180 | SECRETORY CARRIER MEMBRAN | Secretory carrier membrane protein (SCAMP) f | shade67<shade56 | 464.85582 | -4.803988131 | 0.7783166 | -6.17228 | 6.73E-10 | 7.50E-07 |
| MA_10435231g0010 | AT1G12860 | SCREAM 2 (SCRM2) | Encodes ICE2 (Inducer of CBF Expression 2), a t | shade67<shade56 | 1036.0692 | -2.995103713 | 0.7645797 | -3.917321 | 8.95E-05 | 0.00840214 |
| MA_447489g0010 | AT1G15100 | RING-H2 FINGER A2A (RHA2A) | Encodes a putative RING-H2 finger protein RHA | shade67<shade56 | 2475.4907 | -2.089481029 | 0.5191656 | -4.024691 | 5.71E-05 | 0.00619083 |
| MA_1026335g0020 | AT1G15780 | NON-RECOGNITION-OF-BTH 4 (I | mediator of RNA polymerase II transcription su | shade67<shade56 | 34.011685 | -23.05216854 | 5.5165931 | -4.178697 | 2.93E-05 | 0.00369282 |
| MA_10431686g0010 | AT1G18090 | | 5-3 exonuclease family protein;(source:Arapor | shade67<shade56 | 133.77856 | -1.707936457 | 0.3775515 | -4.523718 | 6.08E-06 | 0.00104111 |
| MA_3340g0010 | AT1G20510 | OPC-8:0 COA LIGASE1 (OPCL1) | OPC-8:0 CoA ligase1;(source:Araport11) | shade67<shade56 | 73.538753 | -30 | 6.0072541 | -4.993962 | 5.92E-07 | 0.00015401 |
| MA_7115g0010 | AT1G22640 | MYB DOMAIN PROTEIN 3 (MYB3 | MYB-type transcription factor (MYB3) that rep | shade67<shade56 | 172.44193 | -23.14887515 | 5.8902415 | -3.930038 | 8.49E-05 | 0.00811381 |
| MA_939384g0010 | AT1G26500 | | Pentatricopeptide repeat (PPR) superfamily pr | shade67<shade56 | 76.036931 | -23.46362023 | 5.8912287 | -3.982806 | 6.81E-05 | 0.00709934 |
| MA_15946g0010 | AT1G30220 | INOSITOL TRANSPORTER 2 (INT | Inositol transporter presenting conserved extr | shade67<shade56 | 338.30772 | -19.37093044 | 4.3878454 | -4.414679 | 1.01E-05 | 0.00158504 |
| MA_10433153g0020 | AT1G30290 | | unknown protein | shade67<shade56 | 262.51475 | -27.08410624 | 4.3695575 | -6.198364 | 5.71E-10 | 6.71E-07 |
| MA_9974774g0010 | AT1G32300 | L -GULONO-1,4-LACTONE ( L -GU | D-arabinono-1,4-lactone oxidase family protei | shade67<shade56 | 59.887848 | -20.72426353 | 4.3067491 | -4.812043 | 1.49E-06 | 0.00031613 |
| MA_10432360g0020 | AT1G37130 | NITRATE REDUCTASE 2 (NIA2) | Identified as a mutant resistant to chlorate. En | shade67<shade56 | 1268.3351 | -1.297015908 | 0.25408 | -5.104754 | 3.31E-07 | 0.00012979 |
| MA_19887g0010 | AT1G48500 | JASMONATE-ZIM-DOMAIN PRO | jasmonate-zim-domain protein 4;(source:Arap | shade67<shade56 | 22.796923 | -30 | 6.0105393 | -4.991233 | 6.00E-07 | 0.00015401 |
| MA_86183g0010 | AT1G54200 | BIG GRAIN 3 (BG3) | DNA mismatch repair Msh6-like protein;(sourc | shade67<shade56 | 296.12841 | -3.275875616 | 0.7906427 | -4.143307 | 3.42E-05 | 0.00416318 |
| MA_564894g0010 | AT1G55910 | ZINC TRANSPORTER 11 PRECUR | member of Putative zinc transporter ZIP2 - like | shade67<shade56 | 182.443 | -8.543741989 | 1.4440798 | -5.916392 | 3.29E-09 | 2.58E-06 |
| MA_464469g0010 | AT1G56120 | | Leucine-rich repeat transmembrane protein ki | shade67<shade56 | 43.755794 | -21.51556729 | 4.4004257 | -4.889429 | 1.01E-06 | 0.00023135 |
| MA_401306g0010 | AT1G61065 | | 1,3-beta-glucan synthase component (DUF121 | shade67<shade56 | 232.08101 | -24.50516034 | 5.8901333 | -4.160375 | 3.18E-05 | 0.00389277 |
| MA_16626g0010 | AT1G61250 | SECRETORY CARRIER 3 (SC3) | Encodes a putative secretory carrier membran | shade67<shade56 | 444.83114 | -2.131529526 | 0.5017083 | -4.248544 | 2.15E-05 | 0.00288164 |
| MA_167711g0010 | AT1G64680 | | beta-carotene isomerase D27;(source:Araport1 | shade67<shade56 | 3312.6325 | -1.089032164 | 0.2432031 | -4.477872 | 7.54E-06 | 0.00122246 |
| MA_38922g0010 | AT1G71050 | HEAVY METAL ASSOCIATED ISO | Heavy metal transport/detoxification superfam | shade67<shade56 | 323.46382 | -9.342012204 | 2.2831081 | -4.091796 | 4.28E-05 | 0.00497479 |
| MA_10180304g0010 | AT1G72840 | | Disease resistance protein (TIR-NBS-LRR class); | shade67<shade56 | 165.28471 | -24.62843545 | 5.6158792 | -4.385499 | 1.16E-05 | 0.00178087 |
| MA_10370672g0010 | AT1G77460 | CELLULOSE SYNTHASE INTERACT | Encodes a plasma membrane, microtubule ass | shade67<shade56 | 31.994045 | -4.57047869 | 1.0090288 | -4.529582 | 5.91E-06 | 0.00101675 |
| MA_441084g0010 | AT1G77840 | | Translation initiation factor IF2/IF5;(source:Ara | shade67<shade56 | 367.29826 | -24.06317044 | 5.8899652 | -4.085452 | 4.40E-05 | 0.00505909 |
| MA_14341g0010 | AT1G78020 | FCS LIKE ZINC FINGER 6 (FLZ6) | FCS like zinc finger 6 is induced during energy s | shade67<shade56 | 412.05258 | -11.32909104 | 1.5438017 | -7.338437 | 2.16E-13 | 1.83E-09 |
| MA_587505g0010 | AT1G79620 | VASCULAR-RELATED RLK 1 (VRL | VRLK1 is a LRR kinase involved in switching bet | shade67<shade56 | 16.397498 | -8.007723581 | 1.7295361 | -4.629984 | 3.66E-06 | 0.00065858 |
| MA_8433628g0010 | AT2G01610 | | Plant invertase/pectin methylesterase inhibito | shade67<shade56 | 47.351166 | -22.33584837 | 5.3966668 | -4.138823 | 3.49E-05 | 0.00423322 |
| MA_8300751g0010 | AT2G04570 | | GDSL-motif esterase/acyltransferase/lipase. E | shade67<shade56 | 61.237582 | -18.88374371 | 3.705427 | -5.09624 | 3.46E-07 | 0.00013233 |
| MA_10429950g0020 | AT2G13540 | ABA HYPERSENSITIVE 1 (ABH1) | Encodes a nuclear cap-binding protein that for | shade67<shade56 | 357.93022 | -3.055303903 | 0.7643503 | -3.997256 | 6.41E-05 | 0.0067462 |
| MA_349646g0010 | AT2G14080 | | Disease resistance protein (TIR-NBS-LRR class) | shade67<shade56 | 101.04164 | -11.15758155 | 1.5849578 | -7.039671 | 1.93E-12 | 6.54E-09 |
| MA_17793g0010 | AT2G16850 | PLASMA MEMBRANE INTRINSIC | plasma membrane intrinsic protein 2;(source:A | shade67<shade56 | 62.714376 | -24.34183127 | 5.8938296 | -4.130053 | 3.63E-05 | 0.00437292 |
| MA_8866128g0010 | AT2G17570 | CIS-PRENYLTRANSFERASE 1 (CP | Undecaprenyl pyrophosphate synthetase famil | shade67<shade56 | 486.50293 | -3.685883792 | 0.8846408 | -4.166531 | 3.09E-05 | 0.00383343 |
| MA_5951905g0010 | AT2G18050 | HISTONE H1-3 (HIS1-3) | encodes a structurally divergent linker histone | shade67<shade56 | 236.99984 | -26.56529095 | 5.8901749 | -4.510102 | 6.48E-06 | 0.00109794 |
| MA_346860g0010 | AT2G19130 | | S-locus lectin protein kinase family protein;(so | shade67<shade56 | 207.98178 | -2.253851841 | 0.3374051 | -6.679957 | 2.39E-11 | 4.82E-08 |
| MA_63882g0010 | AT2G19790 | ADAPTOR PROTEIN COMPLEX 4 | Encodes a component of the AP4 complex and | shade67<shade56 | 224.03066 | -21.39560558 | 4.2691118 | -5.011723 | 5.39E-07 | 0.00015401 |
| MA_266310g0010 | AT2G23910 | | NAD(P)-binding Rossmann-fold superfamily pro | shade67<shade56 | 449.84823 | -2.093146683 | 0.4448531 | -4.705253 | 2.54E-06 | 0.00048775 |
| MA_10431396g0010 | AT2G26490 | JINGUBANG (JGB) | JGB contains seven WD40 repeats and is highly | shade67<shade56 | 17.775076 | -24.01944223 | 5.9027062 | -4.069225 | 4.72E-05 | 0.00535193 |
| MA_780853g0010 | AT2G28160 | FER-LIKE IRON DEFICIENCY INDU | Encodes a putative transcription factor that re | shade67<shade56 | 68.760449 | -24.30070684 | 5.5800542 | -4.354923 | 1.33E-05 | 0.00198363 |
| MA_9765350g0010 | AT2G28800 | ALBINO 3 (ALB3) | member of Chloroplast membrane protein ALB | shade67<shade56 | 180.19431 | -26.40437984 | 5.8904643 | -4.482563 | 7.38E-06 | 0.00120048 |
| MA_206815g0010 | AT2G29690 | ANTHRANILATE SYNTHASE 2 (AS | Encode a functional anthranilate synthase pro | shade67<shade56 | 140.64084 | -1.878720942 | 0.4452341 | -4.219625 | 2.45E-05 | 0.00322627 |
| MA_279076g0010 | AT2G30933 | | Carbohydrate-binding X8 domain superfamily | shade67<shade56 | 57.035001 | -19.49922833 | 4.3004765 | -4.534202 | 5.78E-06 | 0.00100058 |
| MA_10436215g0020 | AT2G31750 | UDP-GLUCOSYL TRANSFERASE 7 | Encodes an auxin glycosyltransferase that is lik | shade67<shade56 | 305.02062 | -12.53572629 | 2.8605809 | -4.382231 | 1.17E-05 | 0.00180033 |
| MA_564543g0010 | AT2G31900 | MYOSIN-LIKE PROTEIN XIF (XIF) | Encodes an novel myosin isoform. | shade67<shade56 | 76.765086 | -10.83430494 | 2.767198 | -3.915262 | 9.03E-05 | 0.00843675 |
| MA_1160818g0020 | AT2G32850 | | Protein kinase superfamily protein;(source:Ara | shade67<shade56 | 88.015676 | -21.26675086 | 4.2646005 | -4.98681 | 6.14E-07 | 0.00015463 |
| MA_960018g0020 | AT2G34050 | | ATP synthase F1 complex assembly factor;(sou | shade67<shade56 | 371.48094 | -13.99385857 | 2.3307011 | -6.004141 | 1.92E-09 | 1.66E-06 |
| MA_834070g0010 | AT2G35430 | | Zinc finger C-x8-C-x5-C-x3-H type family protei | shade67<shade56 | 89.347434 | -21.25102673 | 4.3980546 | -4.831915 | 1.35E-06 | 0.00028904 |
| MA_10351640g0010 | AT2G35800 | S-ADENOSYL METHIONINE TRAN | Encodes a predicted calcium-dependent S-ade | shade67<shade56 | 41.524372 | -23.62600107 | 4.2967167 | -5.498617 | 3.83E-08 | 2.31E-05 |
| MA_122962g0010 | AT2G38120 | AUXIN RESISTANT 1 (AUX1) | Encodes an auxin influx transporter. AUX1 resi | shade67<shade56 | 592.19336 | -1.797521412 | 0.3642197 | -4.935267 | 8.00E-07 | 0.00019484 |
| MA_95687g0010 | AT2G38940 | PHOSPHATE TRANSPORTER 1;4 | Encodes Pht1;4, a member of the Pht1 family | shade67<shade56 | 134.32187 | -25.45675205 | 5.893003 | -4.321354 | 1.55E-05 | 0.00223228 |
| MA_10428530g0010 | AT2G39480 | ATP-BINDING CASSETTE B6 (AB | P-glycoprotein 6;(source:Araport11) | shade67<shade56 | 46.515492 | -9.455428442 | 1.7703551 | -5.340979 | 9.24E-08 | 5.08E-05 |
| MA_418372g0010 | AT2G40610 | EXPANSIN A8 (EXPA8) | member of Alpha-Expansin Gene Family. Nami | shade67<shade56 | 61.370234 | -23.02511844 | 5.8923518 | -3.907628 | 9.32E-05 | 0.00866945 |
| MA_634100g0010 | AT2G42360 | | RING/U-box superfamily protein;(source:Arapo | shade67<shade56 | 174.80972 | -23.15469167 | 5.7896012 | -3.999359 | 6.35E-05 | 0.00672822 |
| MA_3206652g0010 | AT2G42850 | CYTOCHROME P450, FAMILY 71 | cytochrome P450, family 718;(source:Araport1 | shade67<shade56 | 42.720765 | -30 | 5.3576359 | -5.599485 | 2.15E-08 | 1.38E-05 |
| MA_253584g0010 | AT2G45550 | CYTOCHROME P450, FAMILY 76 | member of CYP76C | shade67<shade56 | 156.33341 | -3.129882756 | 0.7288422 | -4.294321 | 1.75E-05 | 0.00244747 |
| MA_10427075g0010 | AT2G45560 | CYTOCHROME P450, FAMILY 76 | cytochrome P450 monooxygenase | shade67<shade56 | 59.97644 | -2.642012325 | 0.4197184 | -6.294726 | 3.08E-10 | 3.72E-07 |
| MA_1925g0010 | AT2G48130 | (LTPG15) | Encodes a plasma membrane-localized glycosy | shade67<shade56 | 196.7309 | -1.606474795 | 0.3603114 | -4.458573 | 8.25E-06 | 0.00132264 |
| MA_7555671g0010 | AT3G02470 | S-ADENOSYLMETHIONINE DECA | Encodes a S-adenosylmethionine decarboxylas | shade67<shade56 | 91.999685 | -22.6108394 | 5.6409809 | -4.008317 | 6.12E-05 | 0.00652739 |
| MA_80461g0010 | AT3G04220 | | Disease resistance protein (TIR-NBS-LRR class) | shade67<shade56 | 39.54445 | -30 | 6.0078069 | -4.993503 | 5.93E-07 | 0.00015401 |
| MA_909045g0010 | AT3G07880 | SUPERCENTIPEDE1 (SCN1) | RhoGTPase GDP dissociation inhibitor (RhoGD | shade67<shade56 | 96.913487 | -21.61953728 | 4.3552138 | -4.964059 | 6.90E-07 | 0.00016986 |
| MA_10429484g0010 | AT3G14075 | | Mono-/di-acylglycerol lipase, N-terminal;(sourc | shade67<shade56 | 60.455908 | -23.47161132 | 5.8943013 | -3.982085 | 6.83E-05 | 0.0071034 |
| MA_5681263g0010 | AT3G14360 | OIL BODY LIPASE 1 (ATOBL1) | alpha/beta-Hydrolases superfamily protein;(so | shade67<shade56 | 146.95975 | -21.7894401 | 3.6013222 | -6.0504 | 1.44E-09 | 1.32E-06 |
| MA_403867g0010 | AT3G14820 | | GDSL-motif esterase/acyltransferase/lipase. E | shade67<shade56 | 239.45738 | -26.41920125 | 5.893083 | -4.483087 | 7.36E-06 | 0.00120048 |
| MA_16493g0010 | AT3G16520 | UDP-GLUCOSYL TRANSFERASE 8 | UDP-glucosyl transferase 88A1;(source:Araport | shade67<shade56 | 195.61151 | -27.88664738 | 5.8930942 | -4.732089 | 2.22E-06 | 0.00044153 |
| MA_9959752g0010 | AT3G20630 | UBIQUITIN-SPECIFIC PROTEASE | Encodes a ubiquitin-specific protease. Identica | shade67<shade56 | 65.003028 | -30 | 6.0070734 | -4.994112 | 5.91E-07 | 0.00015401 |
| MA_10428105g0010 | AT3G22060 | | contains Pfam profile: PF01657 Domain of unk | shade67<shade56 | 46.626838 | -18.84022474 | 3.4184677 | -5.511307 | 3.56E-08 | 2.18E-05 |
| MA_10367103g0010 | AT3G22220 | | hAT transposon superfamily;(source:Araport11 | shade67<shade56 | 206.49539 | -23.52400821 | 3.6648839 | -6.418759 | 1.37E-10 | 2.15E-07 |
| MA_51552g0010 | AT3G24503 | ALDEHYDE DEHYDROGENASE 2 | Arabidopsis thaliana aldehyde dehydrogenase | shade67<shade56 | 45.01806 | -22.24427383 | 4.4359404 | -5.014557 | 5.32E-07 | 0.00015401 |
| MA_12314g0010 | AT3G28917 | MINI ZINC FINGER 2 (MIF2) | mini zinc finger 2;(source:Araport11) | shade67<shade56 | 123.11633 | -13.05871064 | 3.2825001 | -3.978282 | 6.94E-05 | 0.0071539 |
| MA_10426995g0010 | AT3G43550 | | GDSL-motif esterase/acyltransferase/lipase. E | shade67<shade56 | 139.89151 | -23.49012238 | 5.4596462 | -4.302499 | 1.69E-05 | 0.00239039 |
| MA_10428913g0020 | AT3G44630 | | Disease resistance protein (TIR-NBS-LRR class) | shade67<shade56 | 41.328588 | -3.626809213 | 0.895874 | -4.048347 | 5.16E-05 | 0.00577497 |
| MA_9820002g0010 | AT3G45970 | EXPANSIN-LIKE A1 (EXLA1) | member of EXPANSIN-LIKE. Naming conventio | shade67<shade56 | 135.13285 | -23.59508938 | 5.8868045 | -4.008132 | 6.12E-05 | 0.00652739 |
| MA_4364416g0010 | AT3G46520 | ACTIN-12 (ACT12) | Member of actin subclass composed of ACT12 | shade67<shade56 | 67.344876 | -19.21428754 | 3.9109052 | -4.913152 | 8.96E-07 | 0.00021201 |
| MA_9934639g0010 | AT3G46960 | (SKI2) | The gene encodes a DExD⁄H box RNA h | shade67<shade56 | 85.589589 | -2.565890682 | 0.6452093 | -3.976835 | 6.98E-05 | 0.00715652 |
| MA_10132652g0010 | AT3G47110 | | Leucine-rich repeat protein kinase family prote | shade67<shade56 | 301.15243 | -19.51797408 | 4.3330031 | -4.504491 | 6.65E-06 | 0.00111734 |
| MA_673539g0010 | AT3G48140 | | B12D protein;(source:Araport11) | shade67<shade56 | 99.159008 | -21.24768603 | 4.3594317 | -4.873958 | 1.09E-06 | 0.00024624 |
| MA_5530991g0010 | AT3G48280 | CYTOCHROME P450, FAMILY 71 | putative cytochrome P450 | shade67<shade56 | 394.25791 | -30 | 4.1804968 | -7.176181 | 7.17E-13 | 3.03E-09 |
| MA_9494886g0010 | AT3G48320 | CYTOCHROME P450, FAMILY 71 | putative cytochrome P450 | shade67<shade56 | 81.257292 | -22.44025384 | 4.4949674 | -4.992306 | 5.97E-07 | 0.00015401 |
| MA_8080485g0010 | AT3G49260 | IQ-DOMAIN 21 (iqd21) | IQ-domain 21;(source:Araport11) | shade67<shade56 | 189.49709 | -24.63962602 | 5.8903625 | -4.183041 | 2.88E-05 | 0.00365557 |
| MA_8912495g0010 | AT3G49990 | | Low temperature viability protein;(source:Ara | shade67<shade56 | 69.482362 | -23.5724293 | 5.8945255 | -3.999038 | 6.36E-05 | 0.00672822 |
| MA_499152g0010 | AT3G50780 | | BTB/POZ domain protein;(source:Araport11) | shade67<shade56 | 181.36179 | -17.95607005 | 4.3590214 | -4.119289 | 3.80E-05 | 0.00453065 |
| MA_7310594g0010 | AT3G51130 | | transmembrane protein;(source:Araport11) | shade67<shade56 | 166.2113 | -22.51264882 | 4.4382384 | -5.072429 | 3.93E-07 | 0.0001471 |
| MA_179915g0020 | AT3G52500 | | Eukaryotic aspartyl protease family protein;(so | shade67<shade56 | 85.077405 | -21.10809367 | 4.2982352 | -4.910874 | 9.07E-07 | 0.00021201 |
| MA_918643g0010 | AT3G57120 | | Protein kinase superfamily protein;(source:Ara | shade67<shade56 | 38.232762 | -23.74454222 | 5.4693731 | -4.341365 | 1.42E-05 | 0.00208079 |

| | | | | | | | | | | |
|---|---|---|---|---|---|---|---|---|---|---|
| MA_10058810g0010 | AT3G61680 | PLASTID LIPASE1 (PLIP1) | PLIP1 encodes a plastid localized phospholipas | shade67<shade56 | 311.26525 | -24.11401178 | 5.8901635 | -4.093946 | 4.24E-05 | 0.00494951 |
| MA_111976g0010 | AT4G04970 | GLUCAN SYNTHASE-LIKE 1 (GSL1 | encodes a gene similar to callose synthase The | shade67<shade56 | 111.36636 | -4.093241089 | 0.7300074 | -5.607123 | 2.06E-08 | 1.34E-05 |
| MA_169781g0010 | AT4G09340 | | SPla/RYanodine receptor (SPRY) domain-conta | shade67<shade56 | 55.161341 | -19.17770461 | 4.0645158 | -4.718325 | 2.38E-06 | 0.00046377 |
| MA_10435813g0010 | AT4G10310 | HIGH-AFFINITY K+ TRANSPORTE | encodes a sodium transporter (HKT1) expresse | shade67<shade56 | 325.95917 | -22.18232556 | 3.1536734 | -7.033806 | 2.01E-12 | 6.54E-09 |
| MA_179643g0010 | AT4G10500 | DMR6-LIKE OXYGENASE 1 (DLO1 | 2-oxoglutarate (2OG) and Fe(II)-dependent oxy | shade67<shade56 | 377.3487 | -12.92130221 | 2.8225575 | -4.57787 | 4.70E-06 | 0.00083178 |
| MA_58710g0010 | AT4G10550 | | Subtilase family protein;(source:Araport11) | shade67<shade56 | 284.90435 | -1.187277451 | 0.3004241 | -3.952005 | 7.75E-05 | 0.00778006 |
| MA_423032g0010 | AT4G10780 | | LRR and NB-ARC domains-containing disease re | shade67<shade56 | 327.37919 | -15.04072897 | 3.6782994 | -4.089044 | 4.33E-05 | 0.00500861 |
| MA_92659g0010 | AT4G18390 | TEOSINTE BRANCHED 1, CYCLOI | TEOSINTE BRANCHED 1, cycloidea and PCF trar | shade67<shade56 | 81.598303 | -22.31837131 | 4.4340774 | -5.033374 | 4.82E-07 | 0.00015401 |
| MA_456473g0010 | AT4G23140 | CYSTEINE-RICH RLK (RECEPTOR- | Arabidopsis thaliana receptor-like protein kina | shade67<shade56 | 98.792615 | -19.52952353 | 4.4650202 | -4.373894 | 1.22E-05 | 0.00185801 |
| MA_945672g0010 | AT4G23340 | | 2-oxoglutarate (2OG) and Fe(II)-dependent oxy | shade67<shade56 | 172.16112 | -24.69937417 | 5.8902701 | -4.19325 | 2.75E-05 | 0.00353428 |
| MA_128616g0010 | AT4G26050 | PLANT INTRACELLULAR RAS GRO | Encodes PIRL8, a member of the Plant Intracell | shade67<shade56 | 96.865151 | -25.06625578 | 5.8910747 | -4.254955 | 2.09E-05 | 0.00281814 |
| MA_17530g0010 | AT4G26890 | MITOGEN-ACTIVATED PROTEIN | member of MEKK subfamily | shade67<shade56 | 338.17932 | -24.49086995 | 5.8898861 | -4.158123 | 3.21E-05 | 0.00391345 |
| MA_31254g0010 | AT4G28200 | | U3 small nucleolar RNA-associated-like protein | shade67<shade56 | 114.00041 | -20.96888821 | 4.4917221 | -4.668341 | 3.04E-06 | 0.0005661 |
| MA_5708g0010 | AT4G32300 | S-DOMAIN-2 5 (SD2-5) | S-domain-2 5;(source:Araport11) | shade67<shade56 | 205.36017 | -12.85435167 | 2.6516443 | -4.847691 | 1.25E-06 | 0.0002739 |
| MA_96242g0010 | AT4G34160 | CYCLIN D3;1 (CYCD3;1) | encodes a cyclin D-type protein involved in the | shade67<shade56 | 28.327684 | -30 | 6.0098127 | -4.991836 | 5.98E-07 | 0.00015401 |
| MA_13722g0010 | AT4G37520 | | Peroxidase superfamily protein;(source:Arapor | shade67<shade56 | 129.64975 | -21.47274279 | 4.4832093 | -4.789592 | 1.67E-06 | 0.00034841 |
| MA_10357974g0010 | AT4G38070 | | transcription factor bHLH131-like protein;(sour | shade67<shade56 | 147.69388 | -2.410789851 | 0.6004615 | -4.014895 | 5.95E-05 | 0.00640439 |
| MA_351763g0010 | AT5G02230 | | Haloacid dehalogenase-like hydrolase (HAD) su | shade67<shade56 | 75.443171 | -18.97876163 | 3.3107345 | -5.732493 | 9.90E-09 | 6.76E-06 |
| MA_19871g0020 | AT5G02960 | | Ribosomal protein S12/S23 family protein;(sou | shade67<shade56 | 188.10599 | -25.08343565 | 5.8905345 | -4.258261 | 2.06E-05 | 0.00278748 |
| MA_96757g0010 | AT5G05340 | PEROXIDASE 52 (PRX52) | Encodes a protein with sequence similarity to p | shade67<shade56 | 157.26187 | -1.263494658 | 0.3037252 | -4.159993 | 3.18E-05 | 0.00389277 |
| MA_10280064g0010 | AT5G07990 | TRANSPARENT TESTA 7 (TT7) | Required for flavonoid 3' hydroxylase activity. | shade67<shade56 | 591.00163 | -2.52383258 | 0.6503736 | -3.880589 | 0.0001042 | 0.0094839 |
| MA_10308567g0040 | AT5G08020 | RPA70-KDA SUBUNIT B (RPA70B | Encodes a homolog of Replication Protein A. rp | shade67<shade56 | 235.74736 | -1.878923288 | 0.3928419 | -4.782899 | 1.73E-06 | 0.0003567 |
| MA_93721g0020 | AT5G09810 | ACTIN 7 (ACT7) | Member of Actin gene family.Mutants are defe | shade67<shade56 | 86.773394 | -30 | 5.4063976 | -5.548981 | 2.87E-08 | 1.79E-05 |
| MA_9106g0010 | AT5G13700 | POLYAMINE OXIDASE 1 (PAO1) | Encodes a protein with polyamine oxidase acti | shade67<shade56 | 133.92391 | -24.89671027 | 5.8905269 | -4.226568 | 2.37E-05 | 0.00315788 |
| MA_73048g0010 | AT5G15780 | | Pollen Ole e 1 allergen and extensin family pro | shade67<shade56 | 3059.7929 | -2.976854627 | 0.682619 | -4.360931 | 1.30E-05 | 0.00193675 |
| MA_7491716g0010 | AT5G17680 | | disease resistance protein (TIR-NBS-LRR class); | shade67<shade56 | 155.4035 | -21.7115806 | 5.5174828 | -3.935052 | 8.32E-05 | 0.0080187 |
| MA_8256257g0010 | AT5G20680 | TRICHOME BIREFRINGENCE-LIKE | Encodes a member of the TBL (TRICHOME BIRE | shade67<shade56 | 223.31806 | -22.76446823 | 4.6127576 | -4.93511 | 8.01E-07 | 0.00019484 |
| MA_95738g0010 | AT5G20700 | | senescence-associated family protein, putative | shade67<shade56 | 104.26097 | -4.201242853 | 0.89282 | -4.705588 | 2.53E-06 | 0.00048775 |
| MA_19854g0020 | AT5G21960 | | encodes a member of the DREB subfamily A-5 | shade67<shade56 | 36.547943 | -21.52738551 | 5.5151606 | -3.903311 | 9.49E-05 | 0.008787 |
| MA_10434800g0010 | AT5G23230 | NICOTINAMIDASE 2 (NIC2) | nicotinamidase 2;(source:Araport11) | shade67<shade56 | 23.253605 | -16.06965962 | 3.6305816 | -4.426194 | 9.59E-06 | 0.00152023 |
| MA_59443g0010 | AT5G26220 | GAMMA-GLUTAMYL CYCLOTRA | ChaC-like family protein;(source:Araport11) | shade67<shade56 | 522.51603 | -6.937424699 | 0.9367568 | -7.40579 | 1.30E-13 | 1.83E-09 |
| MA_7706481g0010 | AT5G45050 | (WRKY16) | Encodes a member of the WRKY Transcription | shade67<shade56 | 78.771933 | -11.19808893 | 2.3505736 | -4.763981 | 1.90E-06 | 0.0003862 |
| MA_10434773g0030 | AT5G47500 | PECTIN METHYLESTERASE 5 (PM | predicted to encode a pectin methylesterase | shade67<shade56 | 31.715567 | -18.86120083 | 3.5596116 | -5.298668 | 1.17E-07 | 5.95E-05 |
| MA_9957075g0010 | AT5G48930 | HYDROXYCINNAMOYL-COA SHIK | At5g48930 has been shown to encode for the | shade67<shade56 | 22.009697 | -30 | 6.0103004 | -4.991431 | 5.99E-07 | 0.00015401 |
| MA_29890g0010 | AT5G49890 | CHLORIDE CHANNEL C (CLC-C) | member of Anion channel protein family | shade67<shade56 | 191.2228 | -22.09201482 | 3.634185 | -6.078946 | 1.21E-09 | 1.19E-06 |
| MA_838949g0010 | AT5G53350 | CLP PROTEASE REGULATORY SU | CLP protease regulatory subunit CLPX mRNA, n | shade67<shade56 | 206.92373 | -10.40859199 | 2.6518131 | -3.925085 | 8.67E-05 | 0.00824038 |
| MA_652482g0010 | AT5G55490 | GAMETE EXPRESSED PROTEIN 1 | Encodes a transmembrane domain containing | shade67<shade56 | 61.204977 | -24.14644217 | 5.8929358 | -4.097523 | 4.18E-05 | 0.00490917 |
| MA_10153873g0010 | AT5G56150 | UBIQUITIN-CONJUGATING ENZY | ubiquitin-conjugating enzyme 30;(source:Arap | shade67<shade56 | 135.84427 | -20.51991807 | 4.4386793 | -4.622978 | 3.78E-06 | 0.00067834 |
| MA_23239g0010 | AT5G58490 | | NAD(P)-binding Rossmann-fold superfamily pro | shade67<shade56 | 492.44562 | -1.143730454 | 0.2256147 | -5.069397 | 3.99E-07 | 0.00014815 |
| MA_83574g0010 | AT5G59970 | | Histone superfamily protein;(source:Araport11 | shade67<shade56 | 223.28613 | -25.64928987 | 3.5723895 | -7.179869 | 6.98E-13 | 3.03E-09 |
| MA_3003293g0010 | AT5G60390 | (EF1ALPHA) | GTP binding Elongation factor Tu family protei | shade67<shade56 | 52.962014 | -22.97246948 | 5.8933811 | -3.898012 | 9.70E-05 | 0.00894233 |
| MA_183219g0010 | AT5G63960 | GIGANTEA SUPPRESSOR5 (GIS5) | Encodes the catalytic subunits of DNA polymer | shade67<shade56 | 72.570338 | -23.25206085 | 5.7098909 | -4.072243 | 4.66E-05 | 0.00529725 |
| MA_10437110g0010 | AT5G64790 | | O-Glycosyl hydrolases family 17 protein;(sourc | shade67<shade56 | 11.086688 | -30 | 6.0115883 | -4.990362 | 6.03E-07 | 0.00015401 |
| MA_205355g0010 | AT5G64950 | MITOCHONDRIAL TRANSCRIPTIO | mTERF family protein which functions in the re | shade67<shade56 | 69.227533 | -24.0330073 | 5.8934183 | -4.07794 | 4.54E-05 | 0.00519707 |
| MA_10238024g0010 | AT5G66580 | | hypothetical protein;(source:Araport11) | shade67<shade56 | 13.514632 | -30 | 6.0106723 | -4.991122 | 6.00E-07 | 0.00015401 |

**Table S3 Allele frequencies of missense mutations in the DEGs in response to SHADE in Norway spruce**

| | | | | | Population-wise allele frequency | | | | | |
|---|---|---|---|---|---|---|---|---|---|---|
| **ContigID:SnpPosition** | **Gene_ID** | **Gene name** | **Amino acid change** | **Allele** | **S1** | **S2** | **S3** | **S4** | **S5** | **S6** |
| MA_86183:6248 | MA_86183g0010 | BIG GRAIN 3 (BG3) | Ala30Val | **Reference (C)** | C:0.952869 | C:0.95283 | C:0.935135 | C:0.884058 | C:0.864675 | C:0.819005 |
| | | | | **Alternate (T)** | T:0.0471311 | T:0.0471698 | T:0.0648649 | T:0.115942 | T:0.135325 | T:0.180995 |
| MA_10351640:1873 | MA_10351640g0010 | S-ADENOSYL METHIONINE TRANSPORTER-LIKE (SAMTL) | Gln119Lys | **Reference (C)** | C:0.972803 | C:0.971154 | C:0.964865 | C:0.927184 | C:0.921147 | C:0.925676 |
| | | | | **Alternate (A)** | A:0.0271967 | A:0.0288462 | A:0.0351351 | A:0.0728155 | A:0.078853 | A:0.0743243 |
| MA_15920:34530 | MA_15920g0010 | MYB-like HTH transcriptional regulator family protein | Val9Met | **Reference (G)** | G:0.959916 | G:0.954545 | G:0.949153 | G:0.914216 | G:0.869091 | G:0.815421 |
| | | | | **Alternate (A)** | A:0.0400844 | A:0.0454545 | A:0.0508475 | A:0.085784 | A:0.130909 | A:0.184579 |
| MA_20782:9490 | MA_20782g0010 | Tetratricopeptide repeat (TPR)-like superfamily protein | Ala180Val | **Reference (C)** | C:0.212185 | C:0.220588 | C:0.287634 | C:0.355392 | C:0.366667 | C:0.372146 |
| | | | | **Alternate (T)** | T:0.787815 | T:0.779412 | T:0.712366 | T:0.644608 | T:0.633333 | T:0.627854 |
| MA_80461:1299 | MA_80461g0010 | Disease resistance protein | Asp154Ala | **Reference (A)** | A:0.991416 | A:0.995074 | A:0.991279 | A:0.992105 | A:0.985102 | A:0.987805 |
| | | | | **Alternate (C)** | C:0.00858369 | C:0.00492611 | C:0.00872093 | C:0.00789474 | C:0.0148976 | C:0.0121951 |
| MA_227006:3453 | MA_227006g0010 | Diphthamide synthesis DPH2 family protein | Arg21Trp | **Reference (C)** | C:0.946721 | C:0.933962 | C:0.884409 | C:0.866995 | C:0.859929 | C:0.777027 |
| | | | | **Alternate (T)** | T:0.0532787 | T:0.0660377 | T:0.115591 | T:0.133005 | T:0.140071 | T:0.222973 |
| MA_423032:1006 | MA_423032g0010 | Disease resistance protein | Ala51Ser | **Reference (G)** | G:0.981405 | G:0.981221 | G:0.986559 | G:0.973171 | G:0.968582 | G:0.965438 |
| | | | | **Alternate (T)** | T:0.018595 | T:0.0187793 | T:0.0134409 | T:0.0268293 | T:0.0314183 | T:0.0345622 |
| MA_447489:1980 | MA_447489g0010 | RING-H2 FINGER A2A (RHA2A) | Ser92Thr | **Reference (T)** | T:0.995885 | T:0.995283 | T:0.986559 | T:0.983412 | T:0.956714 | T:0.945946 |
| | | | | **Alternate (A)** | A:0.00411523 | A:0.00471698 | A:0.0134409 | A:0.0165877 | A:0.0432862 | A:0.0540541 |
| MA_904305:7873 | MA_904305g0010 | Tetratricopeptide repeat (TPR)-like superfamily protein | Arg55Ser | **Reference (G)** | G:0.973333 | G:0.976923 | G:0.956395 | G:0.937173 | G:0.913424 | G:0.917073 |
| | | | | **Alternate (C)** | C:0.0266667 | C:0.0230769 | C:0.0436047 | C:0.0628272 | C:0.0865759 | C:0.0829268 |
| MA_904305:8387 | MA_904305g0010 | Tetratricopeptide repeat (TPR)-like superfamily protein | Ala227Thr | **Reference (G)** | G:0.971311 | G:0.973934 | G:0.957219 | G:0.949761 | G:0.938163 | G:0.938914 |
| | | | | **Alternate (A)** | A:0.0286885 | A:0.0260664 | A:0.0427807 | A:0.0502392 | A:0.0618375 | A:0.061086 |
| MA_918643:2547 | MA_918643g0010 | Protein kinase superfamily protein | Try87His | **Reference (T)** | T:0.653061 | T:0.601415 | T:0.521505 | T:0.464789 | T:0.367544 | T:0.335586 |
| | | | | **Alternate (C)** | C:0.346939 | C:0.398585 | C:0.478495 | C:0.535211 | C:0.632456 | C:0.664414 |
| MA_945672:6851 | MA_945672g0010 | Oxygenase superfamily protein | Asp71Val | **Reference (A)** | A:0.900826 | A:0.887019 | A:0.852151 | A:0.816425 | A:0.833037 | A:0.718182 |
| | | | | **Alternate (T)** | T:0.0991736 | T:0.112981 | T:0.147849 | T:0.183575 | T:0.166963 | T:0.281818 |
| MA_945672:7014 | MA_945672g0010 | Oxygenase superfamily protein | Ser17Gly | **Reference (A)** | A:0.904564 | A:0.889952 | A:0.860215 | A:0.822967 | A:0.831541 | A:0.726852 |
| | | | | **Alternate (G)** | G:0.0954357 | G:0.110048 | G:0.139785 | G:0.177033 | G:0.168459 | G:0.273148 |
| MA_945672:7341 | MA_945672g0010 | Oxygenase superfamily protein | Arg115Ser | **Reference (C)** | C:0.971074 | C:0.978469 | C:0.959677 | C:0.934466 | C:0.935599 | C:0.936364 |
| | | | | **Alternate (A)** | A:0.0289256 | A:0.0215311 | A:0.0403226 | A:0.065534 | A:0.0644007 | A:0.0636364 |
| MA_10430253:15461 | MA_10430253g0010 | Alpha/beta-Hydrolases superfamily protein | Phe61Tyr | **Reference (T)** | T:0.168103 | T:0.207729 | T:0.265537 | T:0.308081 | T:0.313528 | T:0.375 |
| | | | | **Alternate (A)** | A:0.831897 | A:0.792271 | A:0.734463 | A:0.691919 | A:0.686472 | A:0.625 |
| MA_10432704:73817 | MA_10432704g0020 | CHLORORESPIRATORY REDUCTION22 (CRR22) | Ala219Val | **Reference (C)** | C:0.997908 | C:0.992857 | C:0.991848 | C:0.9875 | C:0.979204 | C:0.96789 |
| | | | | **Alternate (T)** | T:0.00209205 | T:0.00714286 | T:0.00815217 | T:0.0125 | T:0.0207957 | T:0.0321101 |
| MA_96757:12081 | MA_96757g0010 | PEROXIDASE 52 (PRX52) | Ala55Thr | **Reference (G)** | G:0.993878 | G:0.992891 | G:0.997312 | G:0.978673 | G:0.975177 | G:0.975225 |
| | | | | **Alternate (A)** | A:0.00612245 | A:0.007109 | A:0.00268817 | A:0.021327 | A:0.0248227 | A:0.0247748 |
| MA_90634:8021 | MA_90634g0010 | Aluminum activated malate transporter family protein | Pro43Ser | **Reference (C)** | C:0.997925 | C:0.987923 | C:0.986486 | C:0.981043 | C:0.97331 | C:0.958904 |
| | | | | **Alternate (T)** | T:0.00207469 | T:0.0120773 | T:0.0135135 | T:0.0189573 | T:0.0266904 | T:0.0410959 |
| MA_89572:19598 | MA_89572g0010 | SET DOMAIN PROTEIN 35 (SDG35) | Leu35Ser | **Reference (A)** | A:0.738589 | A:0.722222 | A:0.765027 | A:0.77451 | A:0.852415 | A:0.844037 |
| | | | | **Alternate (G)** | G:0.261411 | G:0.277778 | G:0.234973 | G:0.22549 | G:0.147585 | G:0.155963 |
| MA_8433628:1321 | MA_8433628g0010 | Plant invertase/pectin methylesterase inhibitor superfamily protein | Met85Leu | **Reference (T)** | T:0.970464 | T:0.953202 | T:0.980447 | T:0.987113 | T:0.996176 | T:0.995327 |
| | | | | **Alternate (A)** | A:0.0295359 | A:0.046798 | A:0.0195531 | A:0.0128866 | A:0.00382409 | A:0.0046729 |
| MA_83446:30566 | MA_83446g0020 | purple acid phosphatase 27;(source:Araport11) | Trp38Ser | **Reference (G)** | G:0.868313 | G:0.848341 | G:0.892473 | G:0.929612 | G:0.974245 | G:0.979638 |
| | | | | **Alternate (C)** | C:0.131687 | C:0.151659 | C:0.107527 | C:0.0703883 | C:0.0257549 | C:0.020362 |
| MA_8344:30649 | MA_83446g0020 | purple acid phosphatase 27;(source:Araport11) | Glu66Lys | **Reference (G)** | G:0.865702 | G:0.841346 | G:0.895161 | G:0.919903 | G:0.975979 | G:0.972727 |
| | | | | **Alternate (A)** | A:0.134298 | A:0.158654 | A:0.104839 | A:0.0800971 | A:0.0240214 | A:0.0272727 |
| MA_8344:30651 | MA_83446g0020 | purple acid phosphatase 27;(source:Araport11) | Glu66Asp | **Reference (G)** | G:0.865702 | G:0.841346 | G:0.895161 | G:0.92029 | G:0.975133 | G:0.972851 |
| | | | | **Alternate (T)** | T:0.134298 | T:0.158654 | T:0.104839 | T:0.0797101 | T:0.0248668 | T:0.0271493 |
| MA_8344:30830 | MA_83446g0020 | purple acid phosphatase 27;(source:Araport11) | Leu126His | **Reference (T)** | T:0.880753 | T:0.885167 | T:0.905405 | T:0.871287 | T:0.857923 | T:0.818605 |
| | | | | **Alternate (A)** | A:0.119247 | A:0.114833 | A:0.0945946 | A:0.128713 | A:0.142077 | A:0.181395 |
| MA_58710:10558 | MA_58710g0010 | Subtilase family protein | Cys24Tyr | **Reference (G)** | G:0.761224 | G:0.755924 | G:0.737838 | G:0.71327 | G:0.738011 | G:0.717489 |
| | | | | **Alternate (A)** | A:0.238776 | A:0.244076 | A:0.262162 | A:0.28673 | A:0.261989 | A:0.282511 |
| MA_5587:58879 | MA_5587g0010 | glycosyltransferase family protein 2 | Cys294Ser | **Reference (A)** | A:0.932735 | A:0.957286 | A:0.921512 | A:0.959459 | A:0.954373 | A:0.987923 |
| | | | | **Alternate (T)** | T:0.0672646 | T:0.0427136 | T:0.0784884 | T:0.0405405 | T:0.0456274 | T:0.0120773 |
| MA_5587:58921 | MA_5587g0010 | glycosyltransferase family protein 2 | Llu280Lys | **Reference (C)** | C:0.886076 | C:0.90625 | C:0.879121 | C:0.935 | C:0.933028 | C:0.962441 |

| | | | | | | | | | | |
|---|---|---|---|---|---|---|---|---|---|---|
| | | | | **Alternate (T)** | T:0.113924 | T:0.09375 | T:0.120879 | T:0.065 | T:0.0669725 | T:0.0375587 |
| MA_5587:58936 | MA_5587g0010 | glycosyltransferase family protein 2 | Gln275Lys | **Reference (G)**<br>**Alternate (T)** | G:0.866109<br>T:0.133891 | G:0.900474<br>T:0.0995261 | G:0.863388<br>T:0.136612 | G:0.921182<br>T:0.0788177 | G:0.925551<br>T:0.0744485 | G:0.949074<br>T:0.0509259 |
| MA_5587:58946 | MA_5587g0010 | glycosyltransferase family protein 2 | Gln271His | **Reference (C)**<br>**Alternate (A)** | C:0.983402<br>A:0.0165975 | C:0.973934<br>A:0.0260664 | C:0.978261<br>A:0.0217391 | C:0.987745<br>A:0.0122549 | C:0.99543<br>A:0.00457038 | C:0.997706<br>A:0.00229358 |
| MA_5587:58965 | MA_5587g0010 | glycosyltransferase family protein 2 | Ala265Val | **Reference (G)**<br>**Alternate (A)** | G:0.968487<br>A:0.0315126 | MG:0.966667<br>A:0.0333333 | G:0.983784<br>A:0.0162162 | G:0.97<br>A:0.03 | G:0.979204<br>A:0.0207957 | G:0.990868<br>A:0.00913242 |
| MA_5587:59010 | MA_5587g0010 | glycosyltransferase family protein 2 | Ile250Thr | **Reference (A)**<br>**Alternate (G)** | A:0.968619<br>G:0.0313808 | A:0.968447<br>G:0.0315534 | A:0.989011<br>G:0.010989 | A:0.9675<br>G:0.0325 | A:0.974074<br>G:0.0259259 | A:0.979167<br>G:0.0208333 |
| MA_5587:59031 | MA_5587g0010 | glycosyltransferase family protein 2 | Pro243Leu | **Reference (G)**<br>**Alternate (A)** | G:0.968354<br>A:0.0316456 | G:0.968293<br>A:0.0317073 | G:0.986111<br>A:0.0138889 | G:0.972081<br>A:0.0279188 | G:0.975791<br>A:0.0242086 | G:0.981132<br>A:0.0188679 |
| MA_52897:17848 | MA_52897g0010 | Protein kinase superfamily protein | Gly194Arg | **Reference (C)**<br>**Alternate (T)** | C:0.57377<br>T:0.42623 | C:0.561905<br>T:0.438095 | C:0.601093<br>T:0.398907 | C:0.629268<br>T:0.370732 | C:0.590909<br>T:0.409091 | C:0.625<br>T:0.375 |
| MA_52897:18331 | MA_52897g0010 | Protein kinase superfamily protein | Val33Ile | **Reference (C)**<br>**Alternate (T)** | C:0.89959<br>T:0.10041 | C:0.899522<br>T:0.100478 | C:0.948925<br>T:0.0510753 | C:0.939904<br>T:0.0600962 | C:0.949558<br>T:0.0504425 | C:0.958904<br>T:0.0410959 |
| MA_510832:1492 | MA_510832g0010 | hAT dimerization domain-containing protein / transposase-like protein | Ala52Thr | **Reference (C)**<br>**Alternate (T)** | C:0.985656<br>T:0.0143443 | C:0.971564<br>T:0.028436 | C:0.973118<br>T:0.0268817 | C:0.898551<br>T:0.101449 | C:0.919183<br>T:0.0808171 | C:0.910314<br>T:0.0896861 |
| MA_4984597:670 | MA_4984597g0010 | Eukaryotic aspartyl protease family protein | Val224Met | **Reference (G)**<br>**Alternate (A)** | G:0.983673<br>A:0.0163265 | G:0.971698<br>A:0.0283019 | G:0.97027<br>A:0.0297297 | G:0.952381<br>A:0.047619 | G:0.944938<br>A:0.0550622 | G:0.950673<br>A:0.0493274 |
| MA_38922:6655 | MA_38922g0010 | HEAVY METAL ASSOCIATED ISOPRENYLATED PLANT PROTEIN 20 (HIPP20) | Phe2Cys | **Reference (T)**<br>**Alternate (G)** | T:0.946721<br>G:0.0532787 | T:0.940758<br>G:0.0592417 | T:0.943243<br>G:0.0567568 | T:0.969484<br>G:0.0305164 | T:0.97242<br>G:0.0275801 | T:0.974886<br>G:0.0251142 |
| MA_31254:22109 | MA_31254g0010 | U3 small nucleolar RNA-associated-like protein | Glu30Asp | **Reference (T)**<br>**Alternate (G)** | T:0.724066<br>G:0.275934 | T:0.758294<br>G:0.241706 | T:0.76087<br>G:0.23913 | T:0.806763<br>G:0.193237 | T:0.814286<br>G:0.185714 | T:0.809417<br>G:0.190583 |
| MA_1925:30754 | MA_1925g0010 | LTPG15 | Pro6Leu | **Reference (G)**<br>**Alternate (A)** | G:0.836735<br>A:0.163265 | G:0.866197<br>A:0.133803 | G:0.897849<br>A:0.102151 | G:0.930622<br>A:0.069378 | G:0.941441<br>A:0.0585586 | G:0.928241<br>A:0.0717593 |
| MA_17793:51035 | MA_17793g0010 | PLASMA MEMBRANE INTRINSIC PROTEIN 2;8 (PIP2;8) | Gly37Glu | **Reference (G)**<br>**Alternate (A)** | G:0.958678<br>A:0.0413223 | G:0.913043<br>A:0.0869565 | G:0.932432<br>A:0.0675676 | G:0.874408<br>A:0.125592 | G:0.855098<br>A:0.144902 | G:0.772936<br>A:0.227064 |
| MA_17793:51051 | MA_17793g0010 | PLASMA MEMBRANE INTRINSIC PROTEIN 2;8 (PIP2;8) | Lys42Asn | **Reference (G)**<br>**Alternate (T)** | G:0.966805<br>T:0.033195 | G:0.978049<br>T:0.0219512 | G:0.991892<br>T:0.00810811 | G:0.990431<br>T:0.00956938 | G:0.985714<br>T:0.0142857 | G:0.990826<br>T:0.00917431 |
| MA_169781:13449 | MA_169781g0010 | SPla/RYanodine receptor (SPRY) domain-containing protein | Arg46Ile | **Reference (C)**<br>**Alternate (A)** | C:0.997826<br>A:0.00217391 | C:0.976923<br>A:0.0230769 | C:0.988827<br>A:0.0111732 | C:0.974619<br>A:0.0253807 | C:0.943916<br>A:0.0560837 | C:0.947005<br>A:0.0529954 |
| MA_15382:33084 | MA_15382g0010 | F-box/RNI-like superfamily protein | Leu249Gln | **Reference (T)**<br>**Alternate (A)** | T:0.991632<br>A:0.0083682 | T:0.990476<br>A:0.00952381 | T:0.98913<br>A:0.0108696 | T:0.956731<br>A:0.0432692 | T:0.925267<br>A:0.0747331 | T:0.958716<br>A:0.0412844 |
| MA_15382:33180 | MA_15382g0010 | F-box/RNI-like superfamily protein | Leu281Pro | **Reference (T)**<br>**Alternate (C)** | T:0.958678<br>C:0.0413223 | T:0.976303<br>C:0.0236967 | T:0.9<br>C:0.1 | T:0.865385<br>C:0.134615 | T:0.840708<br>C:0.159292 | T:0.763636<br>C:0.236364 |
| MA_15382:33267 | MA_15382g0010 | F-box/RNI-like superfamily protein | Gly310Glu | **Reference (G)**<br>**Alternate (A)** | G:0.689956<br>A:0.310044 | G:0.711905<br>A:0.288095 | G:0.585165<br>A:0.414835 | G:0.556931<br>A:0.443069 | G:0.567935<br>A:0.432065 | G:0.461187<br>A:0.538813 |
| MA_14234:43020 | MA_14234g0010 | Adenine nucleotide alpha hydrolases-like superfamily protein | Arg25Gly | **Reference (C)**<br>**Alternate (G)** | C:0.985417<br>G:0.0145833 | C:0.995098<br>G:0.00490196 | C:0.975543<br>G:0.0244565 | C:0.977273<br>G:0.0227273 | C:0.962523<br>G:0.0374771 | C:0.956019<br>G:0.0439815 |
| MA_129478:6147 | MA_129478g0010 | BRASSINOSTEROID-RESPONSIVE RING-H2 (BRH1) | Ile64Val | **Reference (A)**<br>**Alternate (G)** | A:0.103306<br>G:0.896694 | A:0.11165<br>G:0.88835 | A:0.145946<br>G:0.854054 | A:0.141791<br>G:0.858209 | A:0.138989<br>G:0.861011 | A:0.334101<br>G:0.665899 |
| MA_116552:11911 | MA_116552g0010 | CVP2 LIKE 1 (CVL1) | Arg141Thr | **Reference (G)**<br>**Alternate (C)** | G:0.993534<br>C:0.00646552 | G:0.978155<br>C:0.0218447 | G:0.988827<br>C:0.0111732 | G:0.967822<br>C:0.0321782 | G:0.969203<br>C:0.0307971 | G:0.969626<br>C:0.0303738 |
| MA_10265740:3728 | MA_10265740g0010 | lectin protein kinase family protein | Phe60Ser | **Reference (T)**<br>**Alternate (C)** | T:0.975207<br>C:0.0247934 | T:0.983254<br>C:0.0167464 | T:0.931319<br>C:0.0686813 | T:0.930952<br>C:0.0690476 | T:0.901596<br>C:0.0984043 | T:0.905405<br>C:0.0945946 |
| MA_10265740:5430 | MA_10265740g0010 | lectin protein kinase family protein | Asn627Lys | **Reference (C)**<br>**Alternate (G)** | C:1<br>G:0 | C:0.992424<br>G:0.00757576 | C:0.980114<br>G:0.0198864 | C:0.976562<br>G:0.0234375 | C:0.958333<br>G:0.0416667 | C:0.960094<br>G:0.0399061 |
| MA_10265740:5624 | MA_10265740g0010 | lectin protein kinase family protein | Arg692His | **Reference (G)**<br>**Alternate (A)** | G:0.959184<br>A:0.0408163 | G:0.966981<br>A:0.0330189 | G:0.882353<br>A:0.117647 | G:0.866029<br>A:0.133971 | G:0.826585<br>A:0.173415 | G:0.830317<br>A:0.169683 |
| MA_10265740:5737 | MA_10265740g0010 | lectin protein kinase family protein | Val730Leu | **Reference (G)**<br>**Alternate (T)** | G:0.971074<br>T:0.0289256 | G:0.966667<br>T:0.0333333 | G:0.895161<br>T:0.104839 | G:0.877404<br>T:0.122596 | G:0.83659<br>T:0.16341 | G:0.844595<br>T:0.155405 |
| MA_10265740:5784 | MA_10265740g0010 | lectin protein kinase family protein | Gln745His | **Reference (A)**<br>**Alternate (C)** | A:0.946502<br>C:0.0534979 | A:0.96875<br>C:0.03125 | A:0.893443<br>C:0.106557 | A:0.855392<br>C:0.144608 | A:0.832423<br>C:0.167577 | A:0.845622<br>C:0.154378 |

| | | | | | | | | | | |
|---|---|---|---|---|---|---|---|---|---|---|
| MA_10238024:5310 | MA_10238024g0010 | hypothetical protein | Glu147Lys | **Reference (G)**<br>**Alternate (A)** | G:0.960905<br>A:0.0390947 | G:0.95<br>A:0.05 | G:0.932796<br>A:0.0672043 | G:0.908213<br>A:0.0917874 | G:0.922662<br>A:0.0773381 | G:0.864679<br>A:0.135321 |
| MA_10208460:670 | MA_10208460g0010 | NUCLEAR RNA POLYMERASE D2B (NRPD2B) | Val31Leu | **Reference (G)**<br>**Alternate (C)** | G:0.954918<br>C:0.045082 | G:0.943128<br>C:0.056872 | G:0.975806<br>C:0.0241935 | G:0.971292<br>C:0.0287081 | G:0.97043<br>C:0.0295699 | G:0.984018<br>C:0.0159817 |
| MA_388691:5378 | MA_388691g0010 | Photosystem II lipoprotein (PSB27) | Cys52Tyr | **Reference (G)**<br>**Alternate (A)** | G:0.246835<br>A:0.753165 | G:0.256098<br>A:0.743902 | G:0.23743<br>A:0.76257 | G:0.275862<br>A:0.724138 | G:0.27808<br>A:0.72192 | G:0.3<br>A:0.7 |

**Table S4 Allele and genotype frequencies of missense mutations in the nine candidate DEGs in response to SHADE in Norway spruce**

| Gene name | SNP | Gene_ID | Reference allele frequency p-value | Alternate allele frequency p-value | Genotype frequency p-value |
|---|---|---|---|---|---|
| MYB3 | T132R | MA_7115g0010 | **C: 2.98E-40** | **G: 1.64E-06** | **4.77E-52** |
| LOV1 | R156K | MA_16619g0010 | G: 0.424181 | A: 0.139508 | **8.94E-07** |
| LOV1 | D190N | MA_16619g0010 | **G: 0.036124** | **A: 0.016444** | **8.78E-11** |
| SCRM2 | T208M | MA_10435231g0010 | **C: 7.00E-11** | **T: 0.017001** | **3.59E-29** |
| SCRM2 | S238N | MA_10435231g0010 | **G: 2.38E-20** | **A: 0.014526** | **2.64E-39** |
| TCP2 | F4S | MA_92659g0010 | **T: 0.000215** | **C: 4.47E-27** | **9.13E-38** |
| TCP2 | V25A | MA_92659g0010 | **T: 6.18E-07** | **C: 7.93E-11** | **1.87E-12** |
| TCP2 | E32G | MA_92659g0010 | **A: 5.11E-07** | **G: 5.81E-11** | **1.31E-12** |
| TCP2 | E51K | MA_92659g0010 | **G: 3.03E-23** | **A: 0.001302** | **6.37E-36** |
| TCP2 | I53F | MA_92659g0010 | **A: 0.001336** | **T: 1.93E-22** | **5.44E-34** |
| NAC036 | D206E | MA_101849g0010 | **T: 8.36E-05** | **A: 3.48E-09** | **1.13E-09** |
| NAC036 | T237M | MA_101849g0010 | C: 0.758596 | T: 0.090354 | **0.008645** |
| EXPB3 | K3R | MA_7354451g0010 | **A: 1.14E-06** | G: 0.358795 | **4.10E-09** |
| EXPB3 | R6S | MA_7354451g0010 | **G: 4.67E-11** | T: 0.317117 | **4.71E-16** |
| EXPB3 | C32R | MA_7354451g0010 | **T: 3.02E-09** | **C: 3.02E-09** | **3.02E-09** |
| FLZ6 | L88V | MA_14341g0010 | **C: 7.35E-06** | G: 0.434854 | **8.07E-10** |
| VRLK1 | A265V | MA_587505g0010 | C: 0.160674 | **T: 0.005841** | **0.001393** |
| VRLK1 | V283L | MA_587505g0010 | **G: 1.85E-07** | **C: 0.020671** | **4.64E-08** |
| RPS2 | S123N | MA_475302g0010 | **G: 1.24E-27** | A: 0.35738 | **1.36E-37** |
| RPS2 | E166A | MA_475302g0010 | **A: 1.43E-22** | C: 0.590028 | **2.69E-29** |

Significant p-values (p-value<0.05) are in bold

**Table S5 Tukey's post-hoc test for genotype frequencies of missense mutations in the nine candidate DEGs in response to SHADE in Norway spruce**

| Gene name | SNP | Gene_ID | Tukey's p-value S1 vs S2 | Bonferroni p-value S1 vs S2 | Tukey's p-value S1 vs S3 | Bonferroni p-value S1 vs S3 | Tukey's p-value S1 vs S4 | Bonferroni p-value S1 vs S4 | Tukey's p-value S1 vs S5 | Bonferroni p-value S1 vs S5 | Tukey's p-value S1 vs S6 | Bonferroni p-value S1 vs S6 | Tukey's p-value S2 vs S3 | Bonferroni p-value S2 vs S3 | Tukey's p-value S2 vs S4 | Bonferroni p-value S2 vs S4 | Tukey's p-value S2 vs S5 | Bonferroni p-value S2 vs S5 | Tukey's p-value S2 vs S6 | Bonferroni p-value S2 vs S6 | Tukey's p-value S3 vs S4 | Bonferroni p-value S3 vs S4 | Tukey's p-value S3 vs S5 | Bonferroni p-value S3 vs S5 | Tukey's p-value S3 vs S6 | Bonferroni p-value S3 vs S6 | Tukey's p-value S4 vs S5 | Bonferroni p-value S4 vs S5 | Tukey's p-value S4 vs S6 | Bonferroni p-value S4 vs S6 | Tukey's p-value S5 vs S6 | Bonferroni p-value S5 vs S6 |
|---|---|---|---|---|---|---|---|---|---|---|---|---|---|---|---|---|---|---|---|---|---|---|---|---|---|---|---|---|---|---|---|---|
| MYB3 T132R | T132R | MA 7115g0010 | 0.9 | 4.438543 | 0.3809 | 0.7933012 | 0.001 | 8.41E-07 | 0.001 | 0.00E+00 | 0.001 | 0.00E+00 | 0.9 | 5.5308979 | 0.001 | 0.0002866 | 0.001 | 0.00E+00 | 0.001 | 0.00E+00 | 0.0142 | 0.016704 | 0.001 | 9.99E-15 | 0.001 | 0.00E+00 | 0.001 | 0.000147 | 0.001 | 1.44E-08 | 0.0372 | 0.047318 |
| LOV1 R156K | R156K | MA 16619g0010 | 0.7743 | 3.0534676 | 0.9 | 9.9629893 | 0.9 | 8.96E+00 | 0.538 | 1.39E+00 | 0.001 | 1.58E-06 | 0.9 | 6.6357244 | 0.9 | 7.0910793 | 0.9 | 1.37E+01 | 0.0014 | 1.46E-03 | 0.9 | 14.12404 | 0.9 | 4.60E+00 | 0.001 | 8.42E-05 | 0.9 | 4.907642 | 0.001 | 5.80E-05 | 0.001 | 5.79E-05 |
| LOV1 D190N | D190N | MA 16619g0010 | 0.6904 | 2.3489015 | 0.9 | 10.477417 | 0.2199 | 3.78E-01 | 0.001 | 4.49E-04 | 0.001 | 1.92E-09 | 0.9 | 5.1359826 | 0.9 | 6.2452656 | 0.1804 | 2.95E-01 | 0.001 | 1.80E-05 | 0.5121 | 1.267055 | 0.0106 | 1.23E-02 | 0.001 | 2.59E-07 | 0.7444 | 2.787098 | 0.001 | 9.97E-04 | 0.0057 | 0.006423 |
| SCRM2 T208M | T208M | MA 10435231g0010 | 0.9 | 6.6801453 | 0.0351 | 0.0444439 | 0.001 | 6.92E-05 | 0.001 | 0.00E+00 | 0.001 | 0.00E+00 | 0.2527 | 0.4531943 | 0.0029 | 0.0031488 | 0.001 | 2.97E-12 | 0.001 | 6.66E-15 | 0.674 | 2.226297 | 0.001 | 1.06E-04 | 0.001 | 1.12E-07 | 0.0466 | 0.060783 | 0.001 | 1.33E-04 | 0.127 | 0.192767 |
| SCRM2 S238N | S238N | MA 10435231g0010 | 0.9 | 6.9390908 | 0.0595 | 0.0798303 | 0.001 | 2.51E-05 | 0.001 | 0.00E+00 | 0.001 | 0.00E+00 | 0.0091 | 0.0104674 | 0.001 | 1.41E-06 | 0.001 | 0.00E+00 | 0.001 | 0.00E+00 | 0.4508 | 1.01988 | 0.001 | 3.44E-07 | 0.001 | 1.45E-08 | 0.0053 | 0.005981 | 0.001 | 1.74E-04 | 0.5127 | 1.270101 |
| TCP2 F4S | F4S | MA 92659g0010 | 0.2461 | 0.4377259 | 0.0032 | 0.00353 | 0.001 | 1.20E-10 | 0.001 | 0.00E+00 | 0.001 | 0.00E+00 | 0.638 | 1.9760238 | 0.001 | 9.49E-05 | 0.001 | 5.59E-08 | 0.001 | 0.00E+00 | 0.0431 | 0.055716 | 0.0014 | 1.50E-03 | 0.001 | 1.03E-13 | 0.9 | 9.955051 | 0.001 | 7.58E-06 | 0.001 | 1.87E-07 |
| TCP2 V25A | V25A | MA 92659g0010 | 0.1101 | 0.1619267 | 0.0083 | 0.0094848 | 0.001 | 2.74E-06 | 0.001 | 2.18E-10 | 0.001 | 8.44E-10 | 0.9 | 5.3953807 | 0.1032 | 0.150322 | 0.0076 | 8.69E-03 | 0.0017 | 1.88E-03 | 0.593 | 1.693677 | 0.2328 | 4.07E-01 | 0.0577 | 7.72E-02 | 0.9 | 11.06309 | 0.7961 | 3.26E+00 | 0.8389 | 3.693338 |
| TCP2 E32G | E32G | MA 92659g0010 | 0.0808 | 0.1127966 | 0.0067 | 0.0075799 | 0.001 | 1.99E-06 | 0.001 | 1.32E-10 | 0.001 | 5.73E-10 | 0.9 | 5.7775225 | 0.1151 | 0.1705974 | 0.0088 | 1.02E-02 | 0.002 | 2.22E-03 | 0.5926 | 1.691056 | 0.2297 | 4.00E-01 | 0.0575 | 7.68E-02 | 0.9 | 10.99938 | 0.7957 | 3.26E+00 | 0.8416 | 3.721841 |
| TCP2 E51K | E51K | MA 92659g0010 | 0.9 | 8.2469537 | 0.001 | 0.0001536 | 0.001 | 3.33E-13 | 0.001 | 0.00E+00 | 0.001 | 4.33E-14 | 0.0026 | 0.002883 | 0.001 | 1.17E-10 | 0.001 | 0.00E+00 | 0.001 | 2.27E-11 | 0.0388 | 0.049652 | 0.001 | 5.35E-05 | 0.0243 | 2.99E-02 | 0.8234 | 3.531849 | 0.9 | 1.37E+01 | 0.8793 | 4.138478 |
| TCP2 I53F | I53F | MA 92659g0010 | 0.7579 | 2.9050206 | 0.001 | 0.0002009 | 0.001 | 2.23E-13 | 0.001 | 0.00E+00 | 0.001 | 2.33E-14 | 0.0309 | 0.0386213 | 0.001 | 7.27E-09 | 0.001 | 0.00E+00 | 0.001 | 1.49E-09 | 0.0279 | 0.034556 | 0.001 | 2.71E-05 | 0.0159 | 1.89E-02 | 0.8167 | 3.463999 | 0.9 | 1.35E+01 | 0.8839 | 4.191659 |
| NAC036 D206E | D206E | MA 101849g0010 | 0.9 | 7.1059258 | 0.5672 | 1.5466731 | 0.0236 | 2.89E-02 | 0.001 | 6.94E-07 | 0.001 | 4.23E-06 | 0.9 | 5.4945844 | 0.1911 | 0.3168462 | 0.001 | 2.10E-04 | 0.001 | 3.24E-04 | 0.732 | 2.680765 | 0.024 | 2.95E-02 | 0.0165 | 1.97E-02 | 0.6241 | 1.885218 | 0.4121 | 8.89E-01 | 0.9 | 6.968164 |
| NAC036 T237M | T237M | MA 101849g0010 | 0.9 | 5.9875681 | 0.9 | 5.2107547 | 0.7967 | 3.27E+00 | 0.0952 | 1.37E-01 | 0.0066 | 7.51E-03 | 0.9 | 13.521499 | 0.9 | 10.60103 | 0.6454 | 2.03E+00 | 0.1132 | 1.67E-01 | 0.9 | 12.16295 | 0.7731 | 3.04E+00 | 0.1842 | 3.03E-01 | 0.9 | 4.470195 | 0.2571 | 4.63E-01 | 0.6065 | 1.77461 |
| EXPB3 K3R | K3R | MA 7354451g0010 | 0.9 | 12.011758 | 0.9 | 4.9009516 | 0.0094 | 1.08E-02 | 0.001 | 2.56E-05 | 0.001 | 1.63E-05 | 0.9 | 7.1191219 | 0.0301 | 0.0375134 | 0.001 | 3.17E-04 | 0.001 | 1.23E-04 | 0.2375 | 0.417652 | 0.0183 | 2.19E-02 | 0.0048 | 5.36E-03 | 0.9 | 8.255559 | 0.7049 | 2.46E+00 | 0.8839 | 4.191597 |
| EXPB3 R6S | R6S | MA 7354451g0010 | 0.8347 | 3.6497194 | 0.4845 | 1.1474042 | 0.001 | 1.97E-04 | 0.001 | 9.30E-11 | 0.001 | 4.42E-10 | 0.9 | 7.9913455 | 0.0251 | 0.0308244 | 0.001 | 3.37E-06 | 0.001 | 1.90E-06 | 0.1702 | 0.273728 | 0.001 | 3.89E-04 | 0.001 | 1.09E-04 | 0.6542 | 2.085785 | 0.2455 | 4.36E-01 | 0.8501 | 3.813265 |
| EXPB3 C32R | C32R | MA 7354451g0010 | 0.9 | 4.6678307 | 0.0052 | 0.0058414 | 0.0036 | 3.97E-03 | ####### | 3.56E-08 | 0.001 | 1.07E-04 | 0.1251 | 0.1892671 | 0.1098 | 0.1614217 | 0.001 | 9.58E-05 | 0.0102 | 1.18E-02 | 0.9 | 14.62198 | 0.7326 | 2.69E+00 | 0.9 | 6.94E+00 | 0.6699 | 2.19717 | 0.9 | 6.41E+00 | 0.9 | 9.185559 |
| FLZ6 L88V | L88V | MA 14341g0010 | 0.9 | 11.374357 | 0.0179 | 0.0214026 | 0.0184 | 2.21E-02 | 0.001 | 5.09E-07 | 0.001 | 7.60E-06 | 0.06 | 0.0806038 | 0.0636 | 0.0860072 | 0.001 | 1.82E-05 | 0.001 | 9.26E-05 | 0.9 | 14.02866 | 0.7154 | 2.54E+00 | 0.6026 | 1.75E+00 | 0.6324 | 1.939177 | 0.5326 | 1.37E+00 | 0.9 | 9.39412 |
| VRLK1 A265V | A265V | MA 587505g0010 | 0.9 | 11.032665 | 0.9 | 5.6330672 | 0.8851 | 4.204986 | 0.044 | 0.05708 | 0.3175 | 0.6161669 | 0.8323 | 3.6237187 | 0.9 | 7.1254889 | 0.175 | 0.283642 | 0.5636 | 1.5273451 | 0.4262 | 0.935191 | 0.0038 | 0.0042099 | 0.0625 | 0.08431 | 0.6479 | 2.042243 | 0.9 | 5.414246 | 0.9 | 10.20483 |
| VRLK1 V283L | V283L | MA 587505g0010 | 0.9 | 13.19395 | 0.9 | 10.859847 | 0.0046 | 5.12E-03 | 0.001 | 8.20E-05 | 0.0027 | 2.91E-03 | 0.9 | 12.620119 | 0.0118 | 0.0137868 | 0.001 | 5.13E-04 | 0.0074 | 8.45E-03 | 0.0313 | 0.03918 | 0.0027 | 3.00E-03 | 0.0213 | 2.58E-02 | 0.9 | 13.29123 | 0.9 | 1.39E+01 | 0.9 | 14.5287 |
| RPS2 S123N | S123N | MA 475302g0010 | 0.9 | 6.875651 | 0.0097 | 0.0112368 | 0.001 | 1.61E-11 | 0.001 | 0.00E+00 | 0.001 | 0.00E+00 | 0.0012 | 0.0012757 | 0.001 | 7.76E-13 | 0.001 | 0.00E+00 | 0.001 | 0.00E+00 | 0.0077 | 0.008806 | 0.001 | 1.25E-05 | 0.001 | 2.90E-05 | 0.9 | 5.664152 | 0.7174 | 2.56E+00 | 0.9 | 6.590226 |
| RPS2 E166A | E166A | MA 475302g0010 | 0.9 | 11.677818 | 0.0045 | 0.0049875 | 0.001 | 6.33E-13 | 0.001 | 3.33E-15 | 0.001 | 3.06E-13 | 0.0023 | 0.0025296 | 0.001 | 3.73E-13 | 0.001 | 3.33E-15 | 0.001 | 1.87E-13 | 0.0038 | 0.004158 | 0.0109 | 1.26E-02 | 0.0032 | 3.51E-03 | 0.9 | 4.398827 | 0.9 | 1.49E+01 | 0.8875 | 4.232984 |

Significant p-values (p-value<0.05) are highlighted

**Table S6 Hardy Weinberg Equilibrium(HWE) for genotype frequencies of missense mutations in the nine candidate DEGs in response to SHADE in Norway spruce**

| Gene name | SNP | Gene_Id | S1 (lat 55-57) HWE p-value | S2 (lat 58) HWE p-value | S3 (lat 59-60) HWE p-value | S4 (lat 61-62) HWE p-value | S5 (lat 63-64) HWE p-value | S6 (lat 65-67) HWE p-value |
|---|---|---|---|---|---|---|---|---|
| MYB3_T132R | T132R | MA_7115g0010 | 0.6305073 | 0.7079489 | 0.09879334 | 0.8456276 | 0.5690967 | 0.2102694 |
| LOV1_R156K | R156K | MA_16619g0010 | 0.5444456 | 0.7613816 | 0.7349385 | 0.8755682 | 0.3548293 | 0.418793 |
| LOV1_D190N | D190N | MA_16619g0010 | 0.5665429 | **0.005432982** | 0.420296 | **0.01394043** | **0.001147745** | **0.02871752** |
| SCRM2_T208M | T208M | MA_10435231g0010 | 0.5657677 | 0.85023 | 0.05895691 | 0.05068066 | 0.4412498 | 0.2792934 |
| SCRM2_S238N | S238N | MA_10435231g0010 | 0.08930325 | 0.2158439 | **0.01860447** | 0.1180562 | 0.4398115 | 0.1794888 |
| TCP2_F4S | F4S | MA_92659g0010 | **0.001170066** | **0.0171661** | **0.003380215** | 0.4064667 | **0.01912685** | **0.000335543** |
| TCP2_V25A | V25A | MA_92659g0010 | 0.7798684 | 0.6814202 | 0.2724744 | 1 | 0.6166626 | 0.1421518 |
| TCP2_E32G | E32G | MA_92659g0010 | 0.7805884 | 0.6788518 | 0.2724744 | 1 | 0.6166373 | 0.1421518 |
| TCP2_E51K | E51K | MA_92659g0010 | 0.3382155 | **0.000678064** | 0.1981032 | 1 | 0.7584974 | 0.1178837 |
| TCP2_I53F | I53F | MA_92659g0010 | 0.3402181 | **0.000714094** | 0.1205355 | 1 | 0.758248 | 0.1163396 |
| NAC036_D206E | D206E | MA_101849g0010 | 0.07351378 | 1 | 0.3554533 | 0.08390553 | 0.05158337 | **0.03029547** |
| NAC036_T237M | T237M | MA_101849g0010 | 0.3934845 | 0.4284897 | 0.5779759 | 0.3071086 | 0.6738657 | 0.8415818 |
| EXPB3_K3R | K3R | MA_7354451g0010 | 0.07548853 | 0.2311453 | 0.1825768 | 1 | **0.000137283** | **0.01557581** |
| EXPB3_R6S | R6S | MA_7354451g0010 | 0.1956392 | 0.4769058 | **0.0299857** | 0.8717671 | **4.78E-06** | **0.01553658** |
| EXPB3_C32R | C32R | MA_7354451g0010 | 1 | 1 | 1 | 1 | 1 | 1 |
| FLZ6_L88V | L88V | MA_14341g0010 | **7.04E-08** | **0.001073109** | 0.05147672 | **0.02159798** | **1.80E-05** | **0.002130486** |
| VRLK1_A265V | A265V | MA_587505g0010 | 0.1128199 | 0.2850072 | 0.338132 | 1 | 0.1760774 | 0.3675938 |
| VRLK1_V283L | V283L | MA_587505g0010 | **0.0179411** | 0.4710266 | **0.001521785** | **0.01064617** | **2.25E-06** | **0.03230442** |
| RPS2_S123N | S123N | MA_475302g0010 | **3.62E-09** | **0.000151837** | **0.000284223** | **2.97E-09** | **7.97E-18** | **3.74E-08** |
| RPS2_E166A | E166A | MA_475302g0010 | **2.82E-10** | **0.000473692** | **0.000236755** | **6.31E-06** | **5.82E-11** | **5.95E-06** |

Significant p-values (p-value<0.05) are highlighted

**Table S7 Allele frequencies of synonymous mutations in the DEGs in response to SHADE in Norway spruce**

| | | | | Population-wise allele frequency | | | | | | |
|---|---|---|---|---|---|---|---|---|---|---|
| **ContigID:SnpPosition** | **Gene_ID** | **Gene name** | **Amino acid change** | **Allele** | **S1** | **S2** | **S3** | **S4** | **S5** | **S6** |
| MA_92659:14474 | MA_92659g0010 | TCP2 | Tyr69Tyr | **Reference (T)** | T:0.964912 | T:0.958115 | T:0.985955 | T:0.992537 | T:0.992767 | T:0.992991 |
| | | | | **Alternate (C)** | C:0.0350877 | C:0.0418848 | C:0.0140449 | C:0.00746269 | C:0.00723327 | C:0.00700935 |
| MA_92659:15344 | MA_92659g0010 | TCP2 | Arg359Arg | **Reference (C)** | C:0.948661 | C:0.8925 | C:0.826705 | C:0.728856 | C:0.70941 | C:0.564286 |
| | | | | **Alternate (A)** | A:0.0513393 | A:0.1075 | A:0.173295 | A:0.271144 | A:0.29059 | A:0.435714 |
| MA_16619:4700 | MA_16619g0010 | LOV1 | Ala34Ala | **Reference (G)** | G:0.979508 | G:0.992925 | G:0.967914 | G:0.954327 | G:0.939367 | G:0.91704 |
| | | | | **Alternate (A)** | A:0.0204918 | A:0.00707547 | A:0.0320856 | A:0.0456731 | A:0.0606327 | A:0.0829596 |
| MA_14341:1032 | MA_14341g0010 | FLZ6 | Phe8Phe | **Reference (T)** | T:0.963265 | T:0.959906 | T:0.938172 | T:0.915094 | T:0.925569 | T:0.849099 |
| | | | | **Alternate (C)** | C:0.0367347 | C:0.0400943 | C:0.061828 | C:0.0849057 | C:0.0744308 | C:0.150901 |
| MA_101849:25612 | MA_101849g0010 | NAC036 | Phe97Phe | **Reference (T)** | T:0.884774 | T:0.850711 | T:0.859459 | T:0.86019 | T:0.847711 | T:0.808559 |
| | | | | **Alternate (C)** | C:0.115226 | C:0.149289 | C:0.140541 | C:0.13981 | C:0.152289 | C:0.191441 |
| MA_101849:25633 | MA_101849g0010 | NAC036 | Thr104Thr | **Reference (T)** | T:0.584362 | T:0.535545 | T:0.58871 | T:0.581731 | T:0.545936 | T:0.513514 |
| | | | | **Alternate (G)** | G:0.415638 | G:0.464455 | G:0.41129 | G:0.418269 | G:0.454064 | G:0.486486 |
| MA_101849:25735 | MA_101849g0010 | NAC036 | Tyr138Tyr | **Reference (T)** | T:0.965021 | T:0.944175 | T:0.951613 | T:0.953431 | T:0.928187 | T:0.884091 |
| | | | | **Alternate (C)** | C:0.0349794 | C:0.0558252 | C:0.0483871 | C:0.0465686 | C:0.0718133 | C:0.115909 |
| MA_101849:25762 | MA_101849g0010 | NAC036 | Phe147Phe | **Reference (C)** | C:0.944215 | C:0.93 | C:0.942935 | C:0.936893 | C:0.927549 | C:0.974654 |
| | | | | **Alternate (T)** | T:0.0557851 | T:0.07 | T:0.0570652 | T:0.0631068 | T:0.0724508 | T:0.0253456 |

**Fig.S1 Metabolism overview with the DEGs in response to shade in the northern and southern populations of Norway spruce. The network map represents the gene regulation in the northern population as compared to the southern one.**

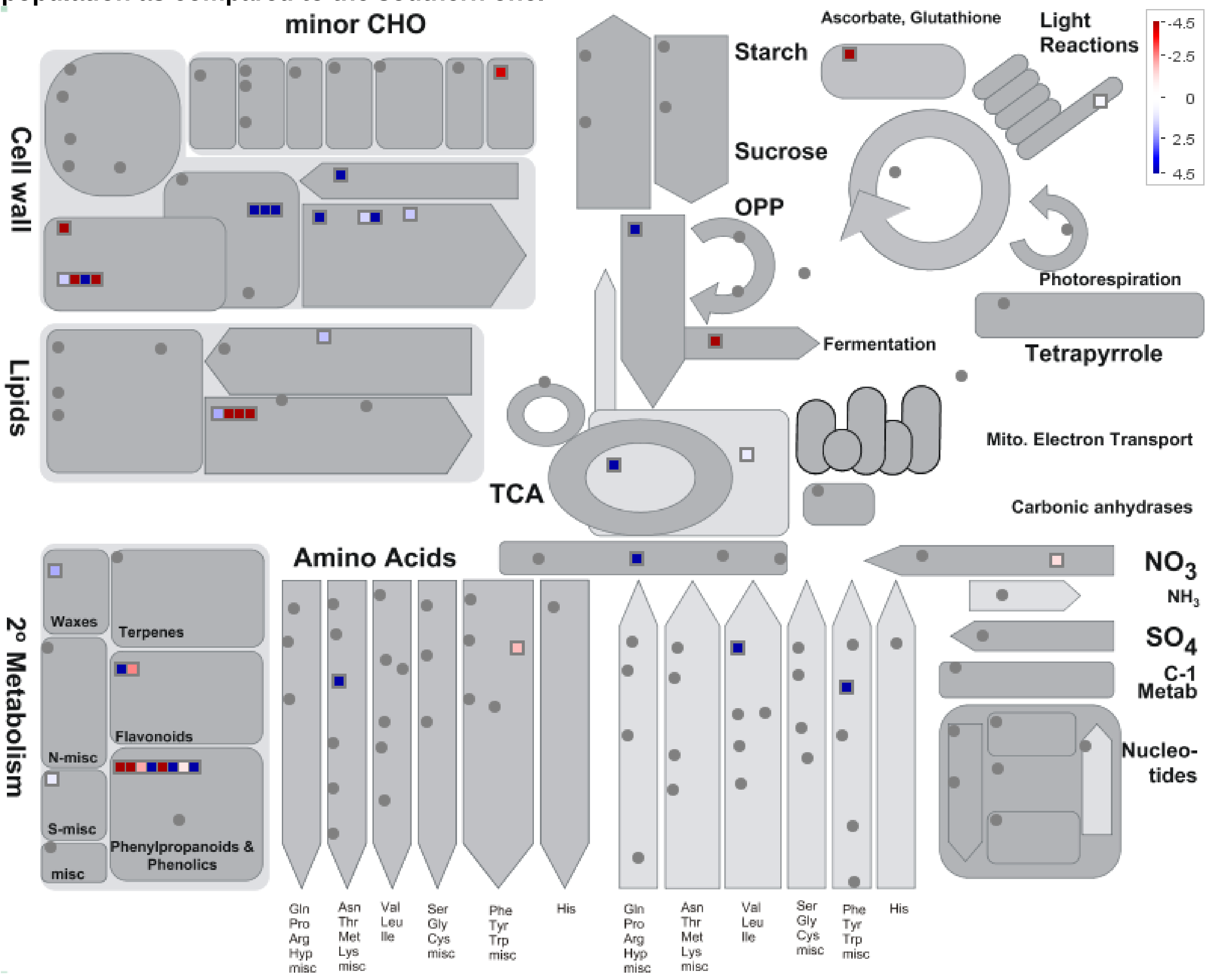

**Fig.S2 Secondary metabolism with the DEGs in response to shade in the northern and southern populations of Norway spruce. The network map represents the gene regulation in the northern population as compared to the southern one.**

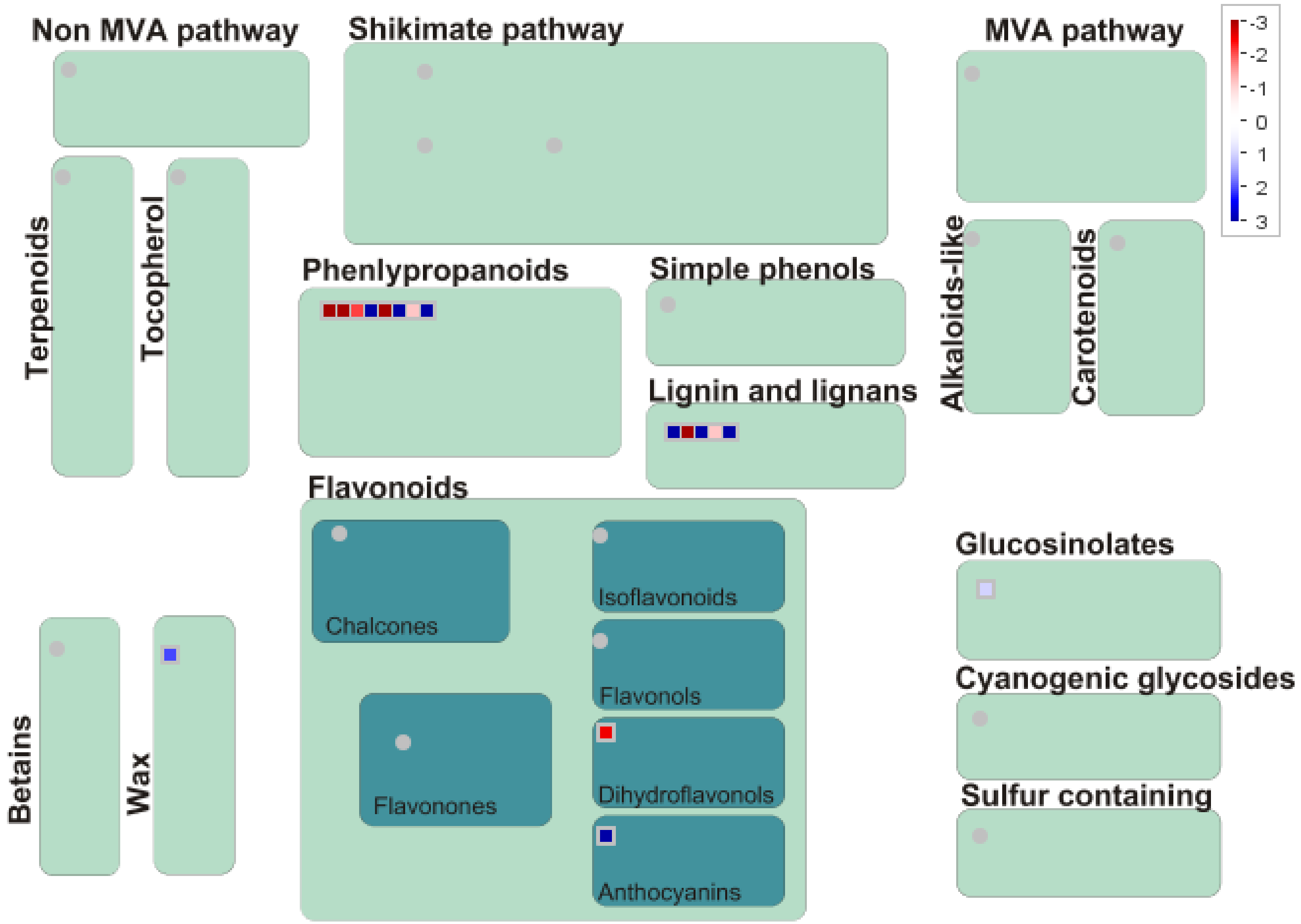

**Fig.S3 Phenylpropanoid pathway with the DEGs in response to shade in the northern and southern populations of Norway spruce. The network map represents the gene regulation in the northern population as compared to the southern one.**

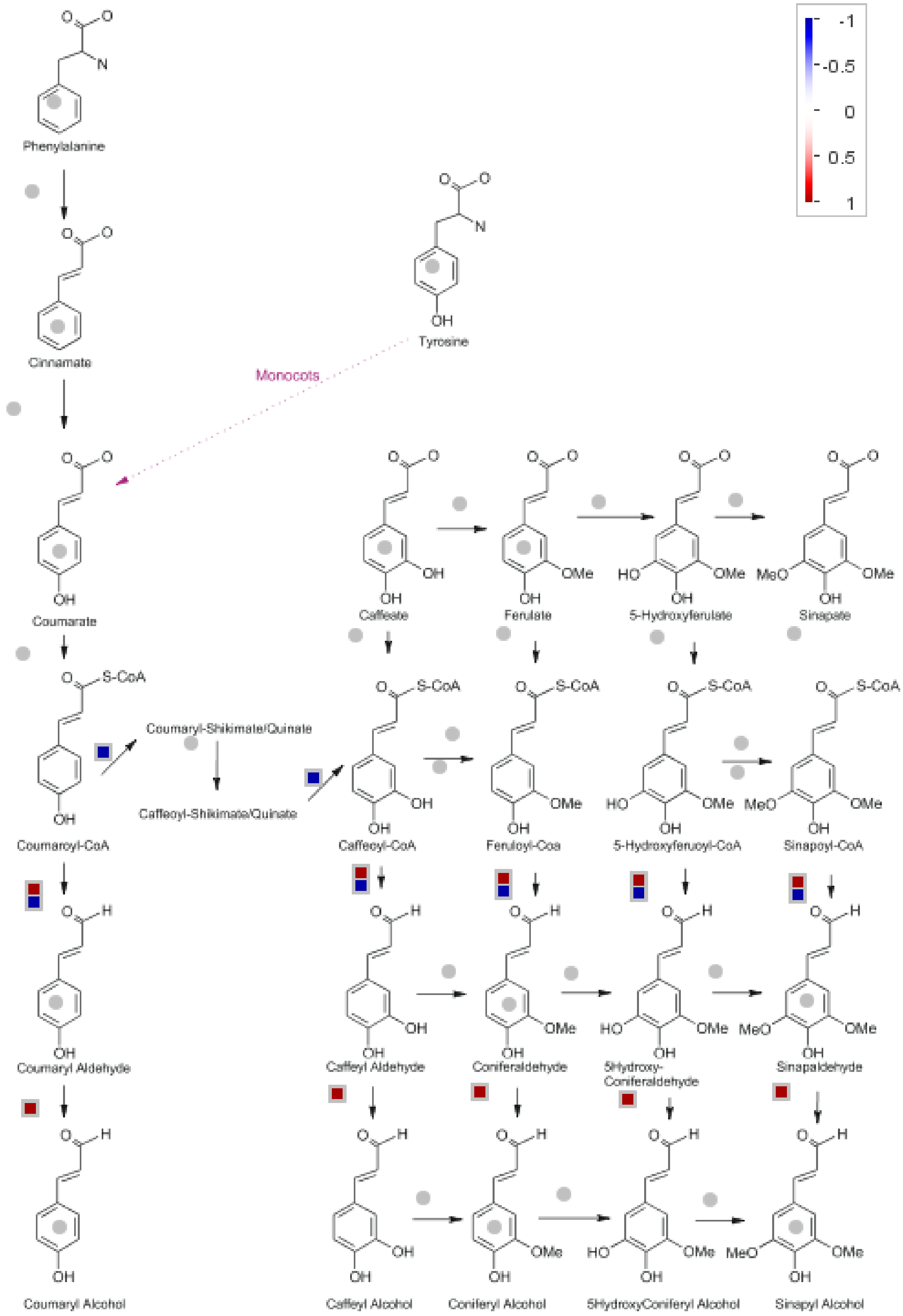

**Fig.S4 Regulation overview with the DEGs in response to shade in the northern and southern populations of Norway spruce. The network map represents the gene regulation in the northern population as compared to the southern one.**

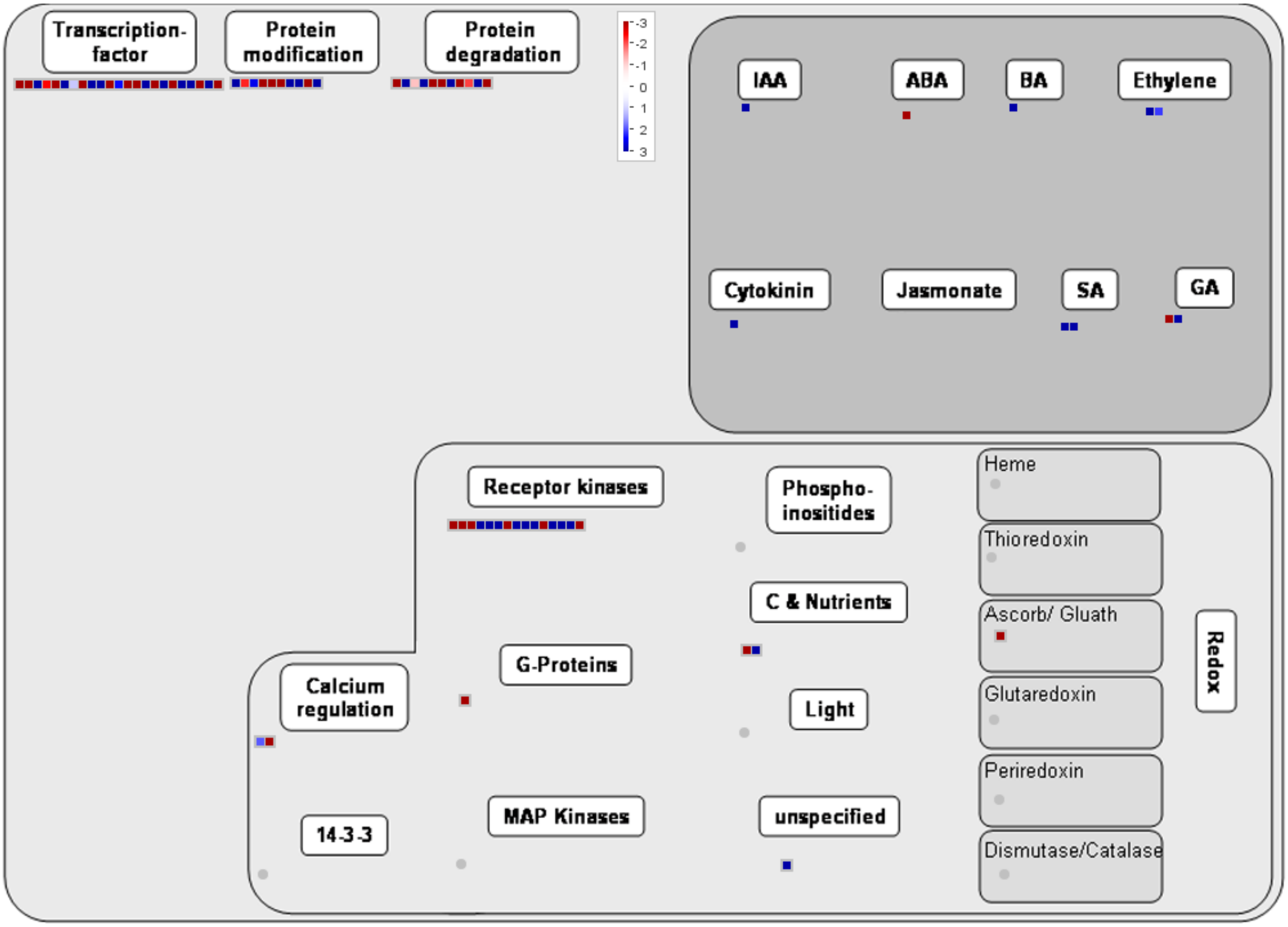

**Fig.S5 Cellular response overview with the DEGs in response to shade in the northern and southern populations of Norway spruce. The network map represents the gene regulation in the northern population as compared to the southern one.**

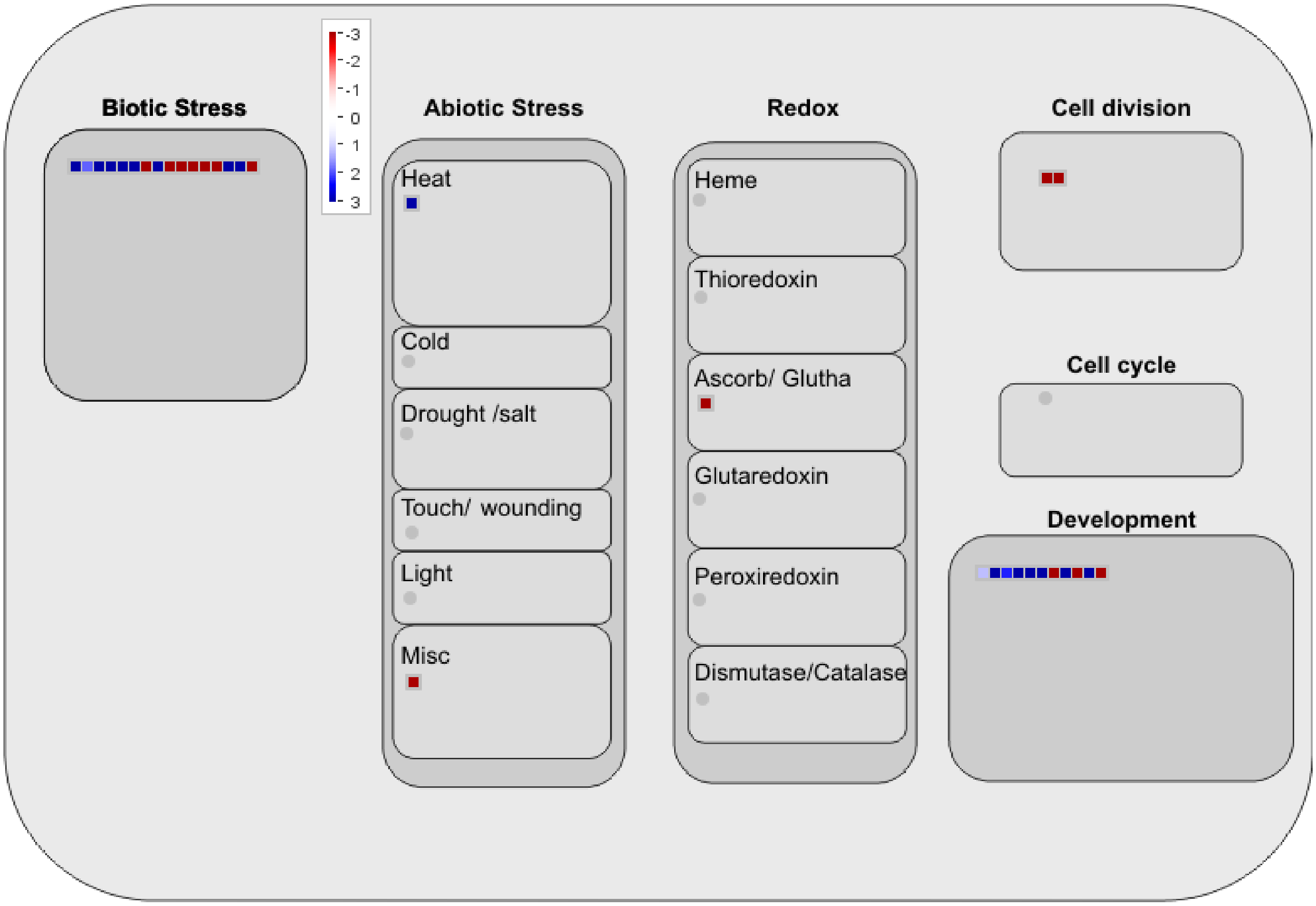

**Fig.S6 Cline with reference to variation in allele and genotype frequencies of SNPs in the *LOV1* gene in Norway spruce populations across Sweden.**

**(a) and (b): allele frequencies of R156K and D190N respectively**

**(c) and (d): genotype frequencies of R156K and D190N respectively, Tukey's *post-hoc* categorization is indicated above the bars.**

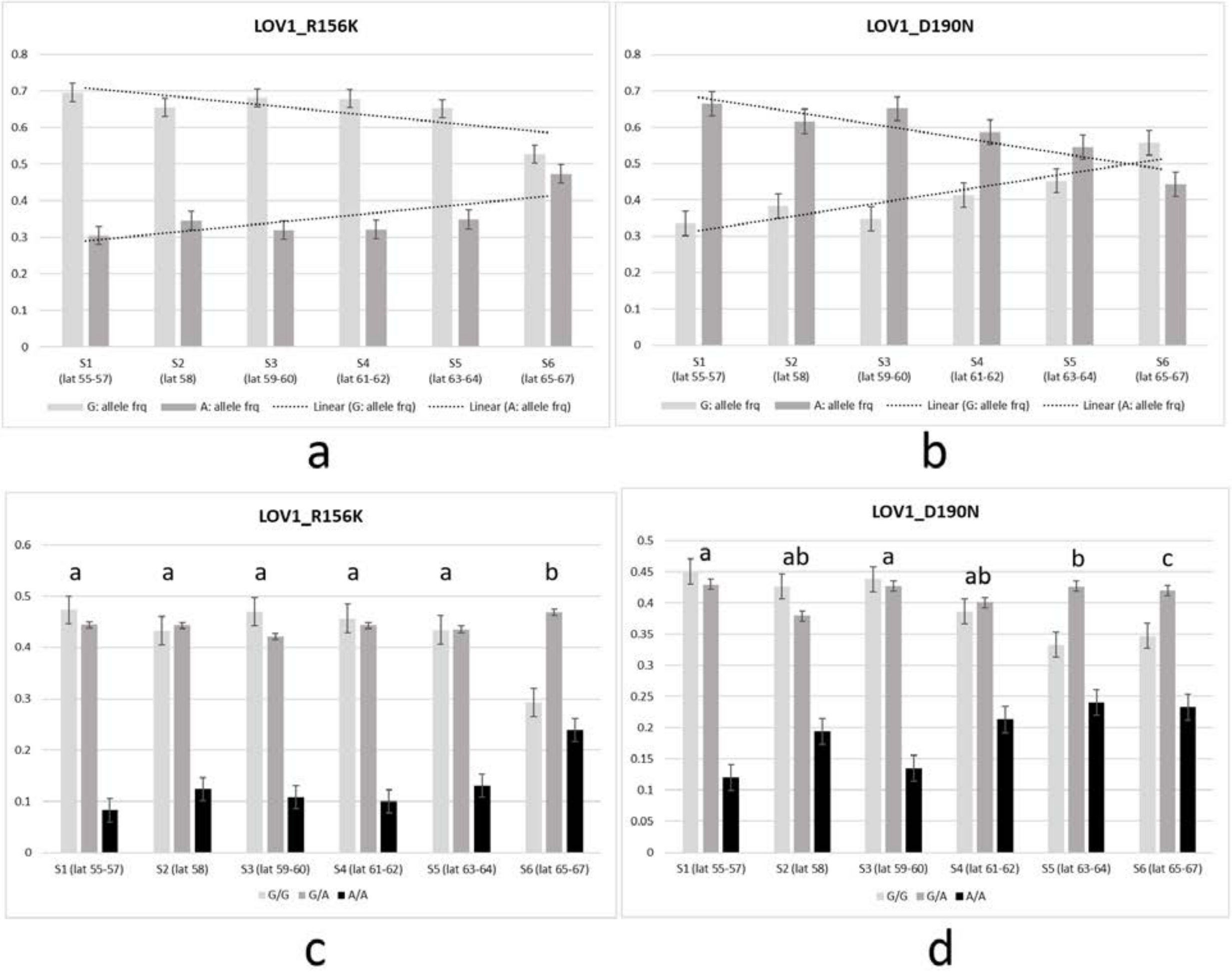

**Fig.S7 Cline with reference to variation in allele and genotype frequencies of SNPs in the *SCRM2* gene in Norway spruce populations across Sweden**

**(a) and (b): allele frequencies of T208M and S238N respectively**

**(c) and (d): genotype frequencies of T208M and S238N respectively, Tukey's *post-hoc* categorization is indicated above the bars.**

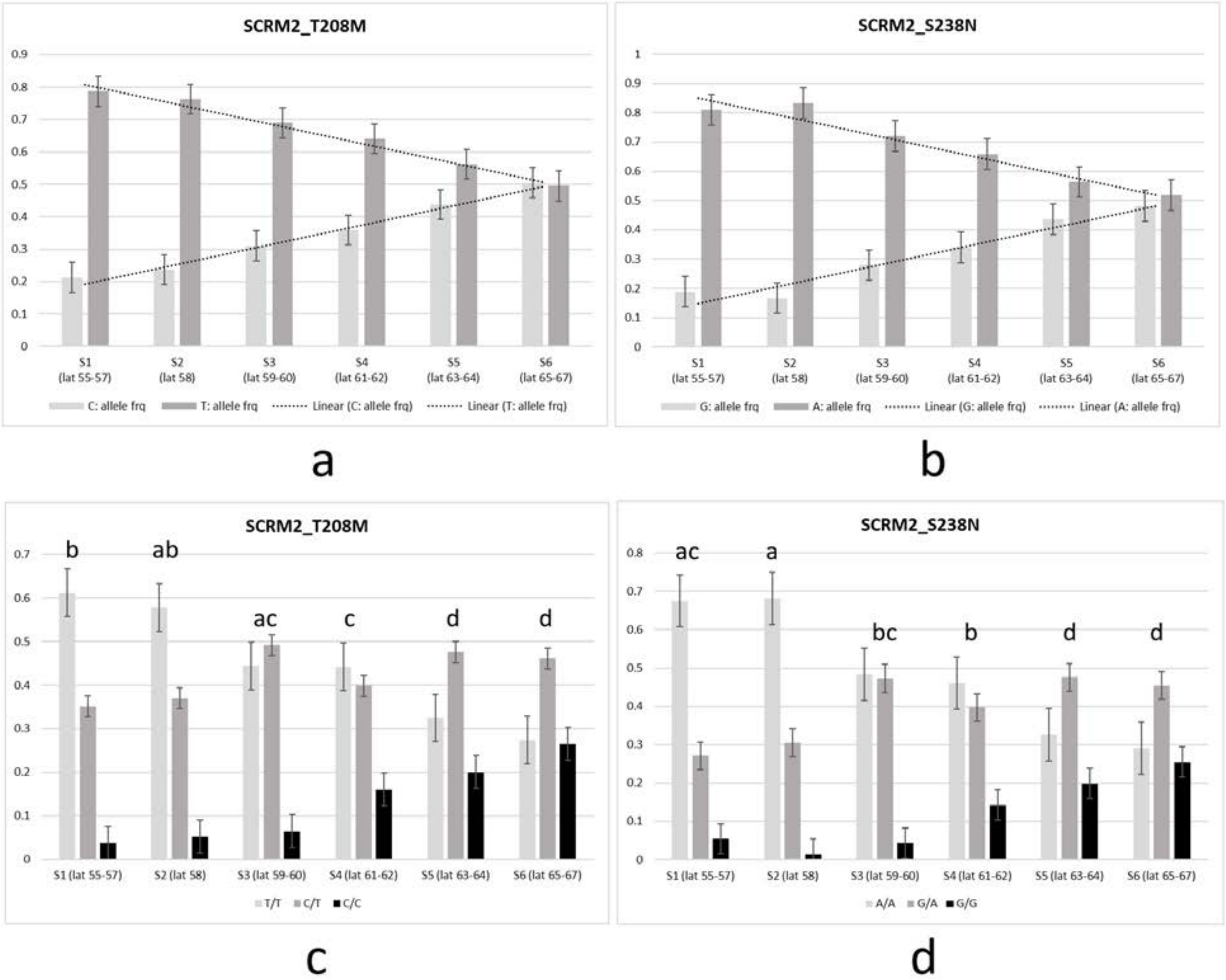

**Fig.S8 Cline with reference to variation in allele and genotype frequencies of SNPs in the *TCP2* gene in Norway spruce populations across Sweden**

**(a) (b) (c) (d) and (e): allele frequencies of F4S, V25A, E32G, E51K and I53F respectively**

**(f) (g) (h) (i) and (j): genotype frequencies of F4S, V25A, E32G, E51K and I53F respectively, Tukey's *post-hoc* categorization is indicated above the bars.**

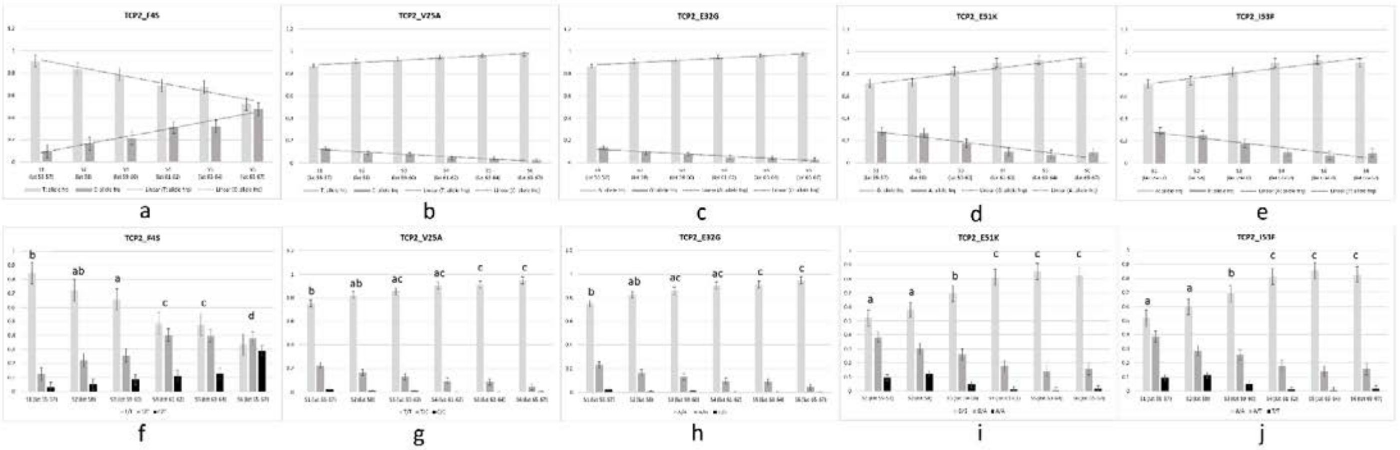

**Fig.S9 Cline with reference to variation in allele and genotype frequencies of SNPs in the *NAC036* gene in Norway spruce populations across Sweden**

**(a) and (b): allele frequencies of D206E and T237M respectively**

**(c) and (d): genotype frequencies of D206E and T237M respectively, Tukey's *post-hoc* categorization is indicated above the bars.**

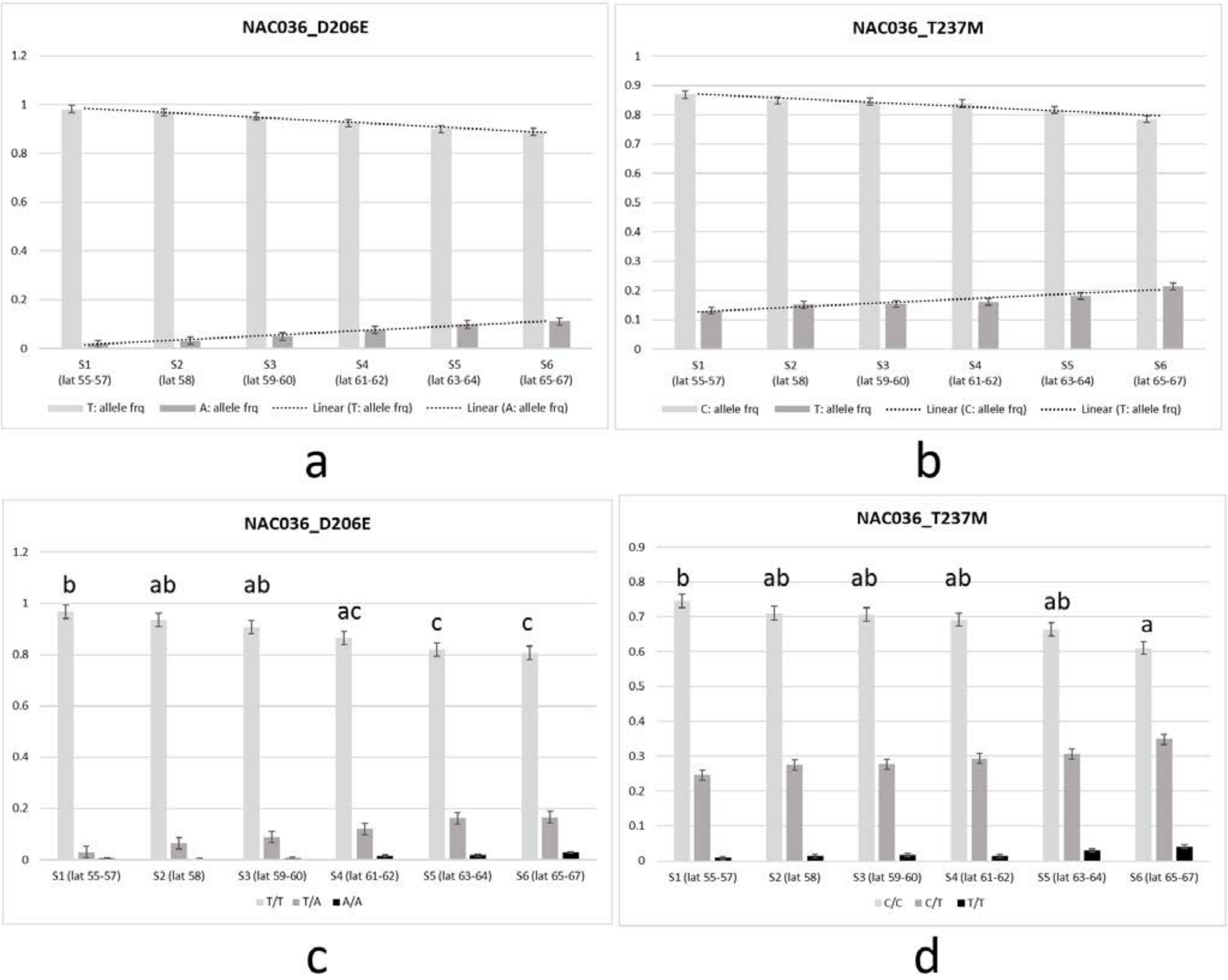

**Fig.S10 Cline with reference to variation in allele and genotype frequencies of SNPs in the *EXPB3* gene in Norway spruce populations across Sweden**

**(a) (b) and (c): allele frequencies of K3R, R6S and C32R respectively**

**(d) (e) and (f): genotype frequencies of K3R, R6S and C32R respectively, Tukey's *post-hoc* categorization is indicated above the bars.**

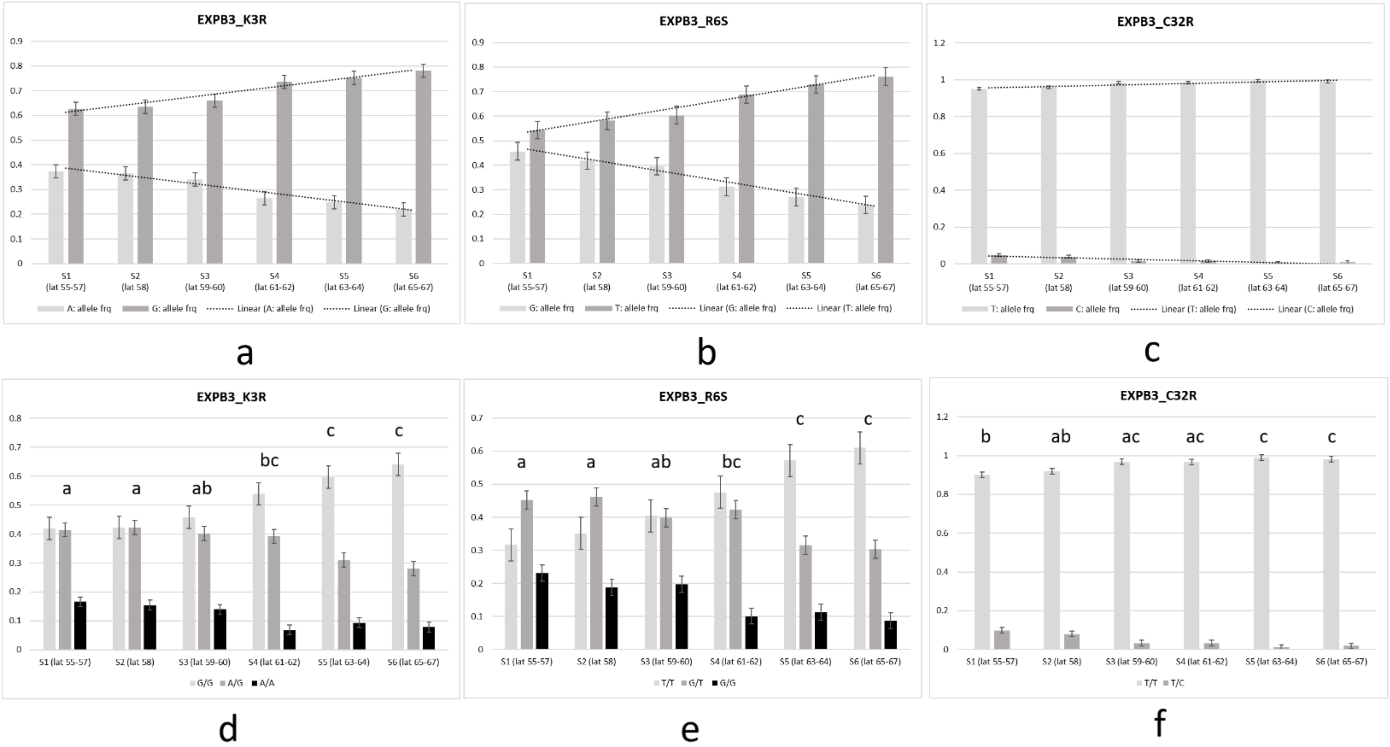

**Fig.S11 Cline with reference to variation in allele and genotype frequencies of SNP in the *FLZ6* gene in Norway spruce populations across Sweden**

**(a) allele frequencies of L88V**

**(b) genotype frequencies of L88V, Tukey's *post-hoc* categorization is indicated above the bars.**

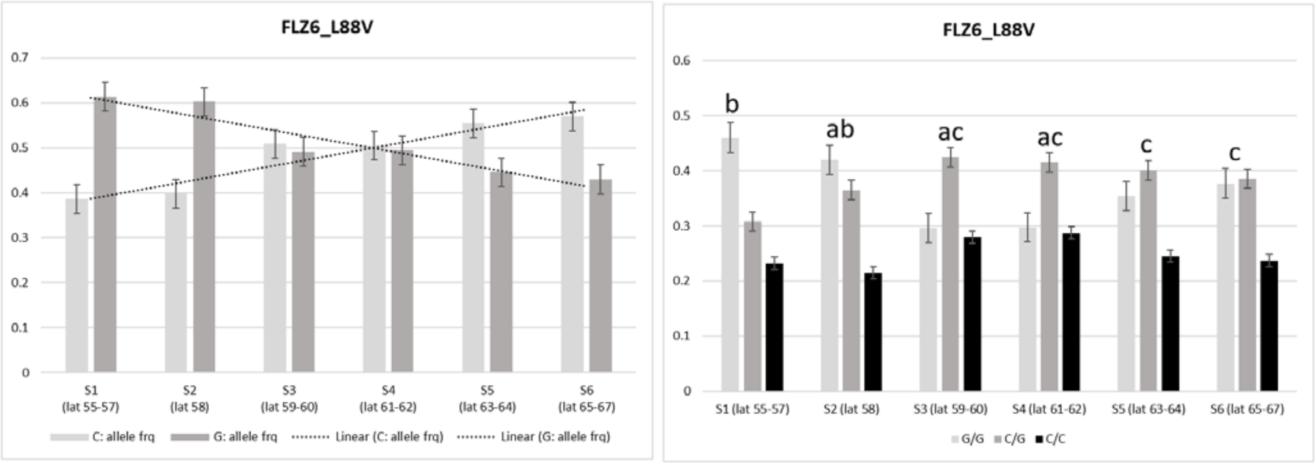


a b

**Fig.S12 Cline with reference to variation in allele and genotype frequencies of SNPs in the *VRLK1* gene in Norway spruce populations across Sweden**

**(a) and (b): allele frequencies of A265V and V283L respectively**

**(c) and (d): genotype frequencies of A265V and V283L respectively, Tukey's *post-hoc* categorization is indicated above the bars.**

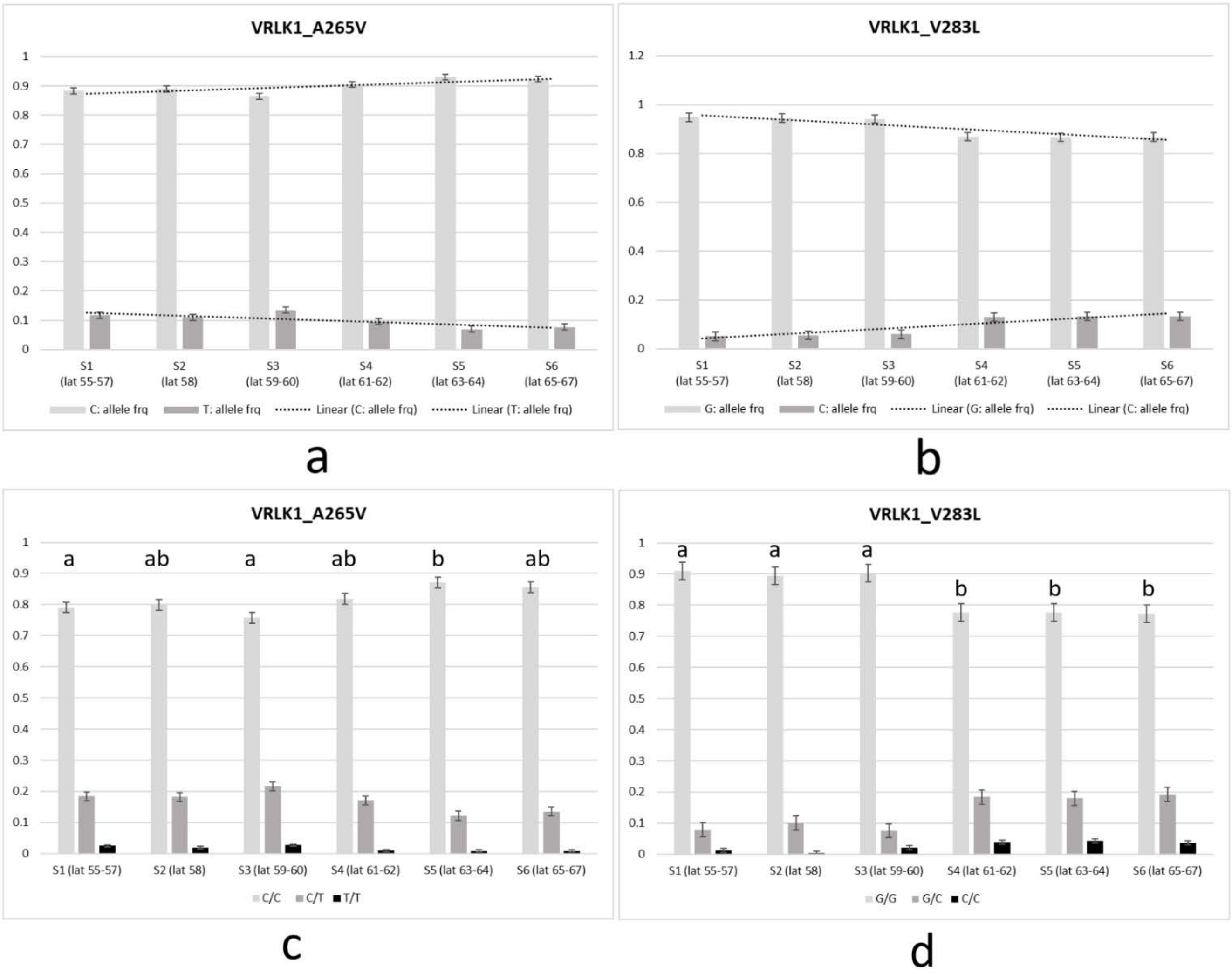

**Fig.S13 Cline with reference to variation in allele and genotype frequencies of SNPs in the *RPS2* gene in Norway spruce populations across Sweden**

**(a) and (b): allele frequencies of S123N and E166A respectively**

**(c) and (d): genotype frequencies of S123N and E166A respectively, Tukey's *post-hoc* categorization is indicated above the bars.**

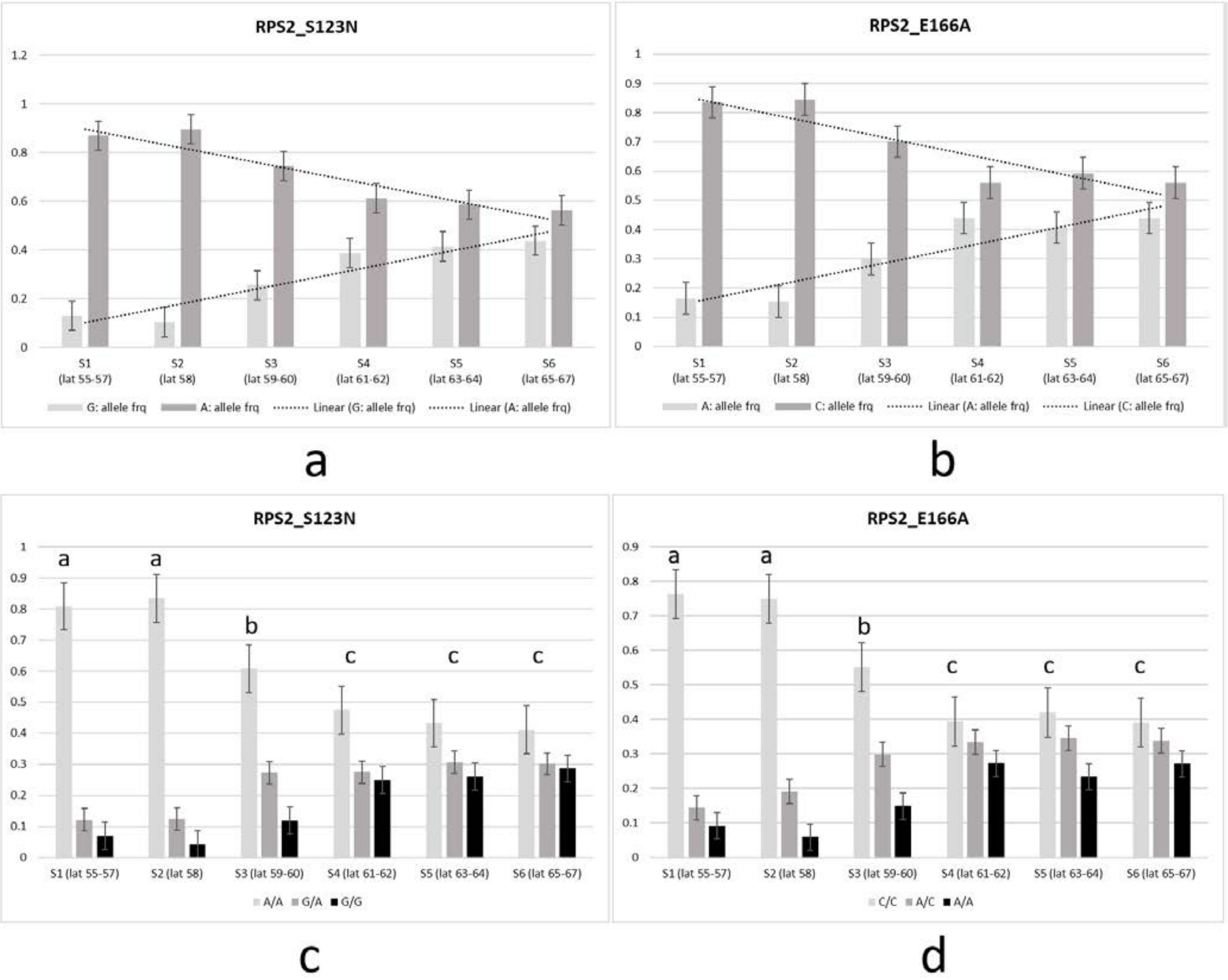

**Fig.S14 Domains of TEOSINTE BRANCHED 1, CYCLOIDEA AND PCF TRANSCRIPTION FACTOR 2 (TCP2)**
**Missense polymorphism sites are highlighted in green in the Norway spruce gene sequences**
**Well characterized domains of the genes are highlighted in other colors in the *Arabidopsis thaliana* gene sequences**
**MA_92659g0010** [551 residues] Norway spruce (http://congenie.org/)
**AT4G18390** [365 residues] *Arabidopsis thaliana* (https://www.arabidopsis.org)

```
CLUSTAL O(1.2.4) multiple sequence alignment


MA_92659p0010      MDSFEEDSDYLEEDNDHFGFNENAVPTEAKDENCKALYEGDLFSSIKPHSEYISVEQERK      60
AT4G18390          ------------------------------------------------------------      0


MA_92659p0010      YGHQGPTAYSYACCCYSSPFSVGHETTGCKKPTNDPAAYPLNGSHKASIADSSPRHMPPG      120
AT4G18390          ------------------------------------------------------------      0


MA_92659p0010      RPCSLIAQHIDPNPCEDEVPVNLMSDMFQFPDFLETYAQSAEWEKKNTLQRKQENNPKRS      180
AT4G18390          ----------------------MIGDLM------KNNNNG-------DVV-DNEVNNRLS      24
                                         ::.*::      :.  :.        :  .:* * : *

MA_92659p0010      RLQFQQPEKSSGNMNSSGFMEPQKIRRHSGNSDRHRKVVTSRGPRDRRMRLSIETAIKFF      240
AT4G18390          RW----------HHN---SSRIIRVSRASGGKDRHSKVLTSKGPRDRRVRLSVSTALQFY      71
                   *           : *     .  :: * **..*** **:**:******:***:.**::*:
42-100: TCP domain: Probably involved in DNA-binding and protein-protein interactions

MA_92659p0010      EIQDRLGFDQPSKAVEWLIQNSRSAIDQLPAISQSALDISSHLKSNSDSLSPFLHRDTDI      300
AT4G18390          DLQDRLGYDQPSKAVEWLIKAAEDSISELPSLNNTHFPTDDENHQNQT-LTTVA------      124
                   ::*****:***********: :..:*.:**::.:: :  ... :.*.  *: .

MA_92659p0010      SGADFVTGFEENMGIADFLSPSLFTEPSILKEQSRSKRKKYNSRKNSVKSALVNSTGRRR      360
AT4G18390          ---------------ANSLSKS----------------------------ACSSNSDTSK      141
                                  *: ** *                            *  ..:.  :

MA_92659p0010      HIAGLKENRS----KARERARERTKEKSCARVKATS-PSPHMNHTRAPEIKALSFQENTA      415
AT4G18390          NSSGLSLSRSELRDKARERARERTAKETKERDHNHTSFTDLLNSGSDPVNSNRQWMASAP      201
                   : :**. .**    ********** :::  * :  :  :  :*    *  .  .:  .:
      151-172: R domain: May mediate protein-protein interactions

MA_92659p0010      S------ASSGNTLNDRMEASISQLYCVAPTLKIGVQEKSLNRRFHHHGCSLTAQDSVKE      469
AT4G18390          SSSPMEYFSSGLILGSGQQTHFPISTNSHPF-SSISDHHHHHPHHQHQEFSFVPDHLI--      258
                   *       ***  *..  :: :       *  .   :.:  : :.:*:  *:. :. :

MA_92659p0010      LAENSHLSGHSNLSFNYSQSQIQSSVQTVAQRYKHQPEDRSMESTGSFTGLCSTTFPPAV      529
AT4G18390          -SPAESNGGAFNLDFNMSTPSGAG------------AAVSAASGGGFSGFNRGTLQSNS      304
                    :  .  .*  **.** *  .  .                *  * *.*:*:   *:

MA_92659p0010      ------------------------------PAMDVDLQSRISDLYVDYSLLQ--------      551
AT4G18390          TNQHQSFLANLQRFPTSESGGGPQFLFGALPAENHHHN-HQFQLYYENGCRNSSEHKGKG      363
                                                 ** : . : :  :** : .  :

MA_92659p0010      --     551
AT4G18390          KN     365
```

**Fig.S15 Domains of LONG VEGETATIVE PHASE 1 (LOV1)**
**Missense polymorphism sites are highlighted in green in the Norway spruce gene sequences**
**Well characterized domains of the genes are highlighted in other colors in the *Arabidopsis thaliana* gene sequences**
**MA_16619g0010** [288 residues] Norway spruce (http://congenie.org/)
**AT2G02450** [414 residues] *Arabidopsis thaliana* (https://www.arabidopsis.org)

```
CLUSTAL O(1.2.4) multiple sequence alignment


MA_16619p0010       -------------------------------------------------MDLPGFRFYPT      11
AT2G02450           MAIVSSTTSIIPMSNQVNNNEKGIEDNDHRGGQESHVQNEDEADDHDHDMVMPGFRFHPT      60
                                                                 *  :*****:**

MA_16619p0010       EEELVGFYLKKRIQGSHPMNFDIVIPTLDLYRYDPCELPSLAHDVEERQWFFFVPRDHKN      71
AT2G02450           EEELIEFYLRRKVEGKRFN--VELITFLDLYRYDPWELPAMAA-IGEKEWYFYVPRDRKY      117
                    ****: ***:::::*.:      :*   ******** ***::*  : *::*:*:****:*

MA_16619p0010       SS-GRPNRLTKSGYWKATGSDRAIRNELLQCIGLKKILVFYKGKAPYGQKTDWIMNEYRL      130
AT2G02450           RNGDRPNRVTTSGYWKATGADRMIRSETSRPIGLKKTLVFYSGKAPKGTRTSWIMNEYRL      177
                     . .****:*.********:** **.*   : ***** ****.**** *  :*.********
                          51-198: NAC domain: DNA-binding and dimerization domain

MA_16619p0010       PDLRSSSVKVRDIVLCRIYRKAASQRSMELQAKPD--HVVKEE-----TAVSGDEYEDTS      183
AT2G02450           PHHETEKYQKAEISLCRVYKRPGVEDHPSVPRSLSTRHHNHNSSTSSRLALRQQQHHSSS      237
                    *. .:.. :  :* ***:*:: . :    .:   . .   *   ::.      *:  :::..:*

MA_16619p0010       ---YSTTLNDGGQLSNPMTEIEVSSLYYQKRERVCS-S-----DQSKTNNGDSLYECL--      232
AT2G02450           SNHSDNNLNNNN----NINNLEKLSTEYSGDGSTTTTTTNSNSDVTIALANQNIYRPMPY      293
                        ...**:..      :.::*   *   *.     . : :      * : :  .:.:*. :

MA_16619p0010       ---E-PIMLSEETRVLKNPKACKTALLELPNMSL------------------------N      263
AT2G02450           DTSNNTLIVSTRNHQDDDETAIVDDLQRLVNYQISDGGNINHQYFQIAQQFHHTQQQNAN      353
                       :  :::* ..:   .: .*     * .* * .:                          *

MA_16619p0010       CPLSQPIATPPALLSTPPPFQPSL----I-------------------------------      288
AT2G02450           ANALQLVAAATTATTLMPQTQAALAMNMIPAGTIPNNALWDMWNPIVPDGNRDHYTNIPF      413
                    .    * :*:  :  :  *   *  :*     *

MA_16619p0010       -      288
AT2G02450           K      414
```

**Fig.S16 Domains of SCREAM 2 (SCRM2)**

**Missense polymorphism sites are highlighted in green in the Norway spruce gene sequences**

**Well characterized domains of the genes are highlighted in other colors in the *Arabidopsis thaliana* gene sequences**

**MA_10435231g0010** [327 residues] Norway spruce (http://congenie.org/)

**AT1G12860** [450 residues] *Arabidopsis thaliana* (https://www.arabidopsis.org)

```
CLUSTAL O(1.2.4) multiple sequence alignment


MA_10435231p0010      ------------------------------------------------------------   0
AT1G12860             MNSDGVWLDGSGESPEVNNGEAASWVRNPDEDWFNNPPPPQHTNQNDFRFNGGFPLNPSE   60


MA_10435231p0010      -------------------------------------------MEMEMAWTGLLDKNFSS   17
AT1G12860             NLLLLLQQSIDSSSSSSPLLHPFTLDAASQQQQQQQQQQEQSFLATKACIVSLLNVPTIN   120
                                                                 :  :  . ..**:    .

MA_10435231p0010      LRAMLDGGGEDGFYATMDANGWTLEALLGSSLSSPGGSVLSNVGGSI----NA--SHHLS   71
AT1G12860             NNTFDDFGFDSGFLGQQ---------FHGN-HQSPNSMNFTGLNHSVPDFLPAPENSSGS   170
                       .:: * * :.** .            : *.  .**..  ::.:. *:     *  .   *

MA_10435231p0010      CSMSTS----------VHTLEGPGGSDQLGLLHRRDVIESGIHKNGGNQKPNG----GVG   117
AT1G12860             CGLSPLFSNRAKVLKPLQVMASS-GS-QPTLFQKRAAMRQSSSSKMCNSESSSEMRKSSY   228
                      *.:*            ::.: .  ** *  *:::* .:...  .:  *.: ..    .

MA_10435231p0010      ERGA----DSVVAV-DSNNNKYVNAANAGRGKKNGLPAKNLMAERRRRKKLNDRLLMLRS   172
AT1G12860             EREIDDTSTGIIDISGLNYESDDHNTNNNKGKKKGMPAKNLMAERRRRKKLNDRLYMLRS   288
                      **       .:: : . * :.  : :* .:***:*:******************* ****

MA_10435231p0010      VVPKVSKMDRSSILGDAAEYLQELLQRIGDLHTELTSGSCNSK---PLADTLADFPCRLN   229
AT1G12860             VVPKISKMDRASILGDAIDYLKELLQRINDLHTELESTPPSSSSLHPLTPTPQTLSYRVK   348
                      ****:*****:****** :**:******.****** *   .*.   **: *   :  *::
                         263–312: BHLH domain

MA_10435231p0010      GELQASF---SSTVEPAKVEVRSREGKSLHINMFCSEKPGLLLSTIRALDELGLDVKQAV   286
AT1G12860             EELCPSSSLPSPKGQQPRVEVRLREGKAVNIHMFCGRRPGLLLSTMRALDNLGLDVQQAV   408
                       **  *    *  .  :   :**** ****:::*:***..:*******:****:*****:***
                         378–450: ACT domain: involved in the binding of small ligands

MA_10435231p0010      ISCLNGFALDVLQEKQSM-GGDVTAEEIKTLLLHTAGYQAGL    327
AT1G12860             ISCFNGFALDVFRAEQCQEDHDVLPEQIKAVLLDTAGYAGLV    450
                      ***:*******:: :*.   .**  *:**::**.**** . :
```

**Fig.S17 Domains of NAC DOMAIN CONTAINING PROTEIN 36 (NAC036)**
**Missense polymorphism sites are highlighted in green in the Norway spruce gene sequences**
**Well characterized domains of the genes are highlighted in other colors in the *Arabidopsis thaliana* gene sequences**
**MA_101849g0010** [306 residues] Norway spruce (http://congenie.org/)
**AT2G17040** [276 residues] *Arabidopsis thaliana* (https://www.arabidopsis.org)

```
CLUSTAL O(1.2.4) multiple sequence alignment


MA_101849p0010      -------------------------MAMPDKICN------DGVVQGELMDLPGFRFYPTE     29
AT2G17040           MNSDGVWLDGSGESPEVNNGEAASWVRNPDEDWFNNPPPPQHTNQNDFRFNGGFPLNPSE     60
                                             :  **:         : . *.::    ** : *:*

MA_101849p0010      EELLSFYMKKKIQAGHS-------LN------------------F---------DKIIPT     55
AT2G17040           NLLLLL--QQSIDSSSSSSPLLHPFTLDAASQQQQQQQQQQEQSFLATKACIVSLLNVPT     118
                    : ** :  ::.*::. *       :.                  *            :**
                         6-152: NAC domain: DNA-binding and dimerization domain

MA_101849p0010      MDLYQYDPWELPGFAHDAGEQQWFFFVPRYHNKRARPNRLAVSGYWKATGSDRAVRNELL     115
AT2G17040           INNNTFDD-----FGFDSG-----FLGQQFHGNHQSPNSMNFTGLNHSVPDFLPAPENSS     168
                    ::   :*      *..*:*     *:  ::*.::  ** : .:*  ::. .   . ::

MA_101849p0010      QCIGLKKILV-------FYKGKSPYGQKTDWIMNEYRMPEFSSSAPKKKMDMVLCRIYRK     168
AT2G17040           GSCGLSPLFSNRAKVLKPLQVMASSGSQPTLFQKRAAMRQ--------SSSSKMCNSESS     220
                     . **. ::          :  :  *.:   : :.  * :        . .  :*.   .

MA_101849p0010      TATRKSLEQRAKTNGVDKEEPVVSSDVYEDKSHSLT-----------------------V     205
AT2G17040           SEMRKSSYEREIDDT-STGIIDISGLNYESDDHNTNNNKGKKKGMPAKNLMAERRRRKKL     279
                    :  ***  :*   :  ..    :*.  **...*. .                       :

MA_101849p0010      DVRCLSN----PM-------------------------------------------IEAK     218
AT2G17040           NDRLYMLRSVVPKISKMDRASILGDAIDYLKELLQRINDLHTELESTPPSSSSLHPLTPT     339
                    : *        *                                            :  .

MA_101849p0010      VSSVQYLDDEGVCSSRSKTKDESS---------LFDCLEPIMFCEDEETTVLKTSK----     265
AT2G17040           PQTLSYRVKEELCPSSSLPSPKGGQPRVEVRLREGKAVNIHMFCGRRPGLLLSTMRALDN     399
                     .::.*  .* :* * *  . :..           ..::  ***  .   :*.* :

MA_101849p0010      ----------------------PKACKPPQLELPKLPMDFLFGQQITTPYALLATPSPFQ     303
AT2G17040           LGLDVQQAVISCFNGFALDVFRAEQCQEDHDVLPEQIKAVLLD---TAGYAGLV------     450
                                           : *:  :  **:    .*:.   *: ** *.

MA_101849p0010      IPL   306
AT2G17040           ---   450
```

**Fig.S18 Domains of EXPANSIN B3 (EXPB3)**

**Missense polymorphism sites are highlighted in green in the Norway spruce gene sequences**

**Well characterized domains of the genes are highlighted in other colors in the *Arabidopsis thaliana* gene sequences**

**MA_7354451g0010** [71 residues] Norway spruce (http://congenie.org/)

**AT4G28250** [264 residues] *Arabidopsis thaliana* (https://www.arabidopsis.org)

```
CLUSTAL O(1.2.4) multiple sequence alignment


MA_7354451p0010      MRKTVRLQWPLVPNFCLDVTAFCFVLLLLFSCKEPALGLEHTELNRDDLEWRPATATWYG    60
AT4G28250            -----------MQLFPVMLATLCIVLQLLIGSS----ALATTNRHVSNSHWLPAVATWYG    45
                                :  * : ::::*:** **:...    .*  *: : .: .* **.*****

MA_7354451p0010      SPDGDGSDGIL-------------------------------------------------    71
AT4G28250            SPNGDGSDGGACGYGTLVDVKPLHARVGAVNPILFKNGEGCGACYKVRCLDKSICSRRAV    105
                     **:******

MA_7354451p0010      ------------------------------------------------------------    71
AT4G28250            TVIITDECPGCSKTSTHFDLSGAVFGRLAIAGESGPLRNRGLIPVIYRRTACKYRGKNIA    165
```

**54-162: Domain: Expansin-like EG45**

```
MA_7354451p0010      ------------------------------------------------------------    71
AT4G28250            FHVNEGSTDFWLSLLVEFEDGEGDIGSMHIRQAGAREWLEMKHVWGANWCIIGGPLKGPF    225
```

**175-256: Domain: Expansin-like CBD**

```
MA_7354451p0010      ---------------------------------------        71
AT4G28250            SIKLTTLSAGKTLSATDVVPRNWAPKATYSSRLNFSPVL        264
```

**Fig.S19 Domains of FCS LIKE ZINC FINGER 6 (FLZ6)**
**Missense polymorphism sites are highlighted in green in the Norway spruce gene sequences**
**Well characterized domains of the genes are highlighted in other colors in the *Arabidopsis thaliana* gene sequences**
**MA_14341g0010** [180 residues] Norway spruce (http://congenie.org/)
**AT1G78020** [162 residues] *Arabidopsis thaliana* (https://www.arabidopsis.org)

```
CLUSTAL O(1.2.4) multiple sequence alignment


MA_14341p0010       MFEDHLRFDNLAAVPLDLEVELAFPEILSEEMMRYPGKRPCPRPRPAAAPIRRTSSTAML      60
AT1G78020           ---------------------------------MLLGKRQ-------RPPINRTTSLSEI      20
                                                    ***          **.**:* : :

MA_14341p0010       DSALNV-----------------------------DHDDRYKQSPEDGGASRKAAALTSH      91
AT1G78020           KFDLNLPSESEPSNQQKPTVASPYGSNGQAVTAAVDQNRGFLDQR-LLSMVTPRGNLRRH      79
                    .  **:                             *::  : :.    .     . *  *

MA_14341p0010       VHVFPEVAYFLQACFLCKRRLGPDTDIYMYRGDAAFCSAECRHEQIVIDERKEKCSAEVR      151
AT1G78020           SGDFSDAGHFLRSCALCERLLVPGRDIYMYRGDKAFCSSECRQEQMAQDERKEKGKSAAP      139
                       * :..:**::* **:* * *. ******** ****:***:**:. ****** .: .
```

**88-132: FLZ-type zinc finger: likely to be involved in protein-protein interaction.**

```
MA_14341p0010       KMKE-SSAPANNRQSSSTNQSVRAGTVAAA- 180
AT1G78020           AKEPAVTAPARAKPG--------KGRAAAAV 162
                      :   :***. : .         * .***
```

**Fig.S20 Domains of VASCULAR-RELATED RLK 1 (VRLK1)**

**Missense polymorphism sites are highlighted in green in the Norway spruce gene sequences**

**Well characterized domains of the genes are highlighted in other colors in the *Arabidopsis thaliana* gene sequences**

**MA_587505g0010** [635 residues] Norway spruce (http://congenie.org/)

**AT1G79620** [971 residues] *Arabidopsis thaliana* (https://www.arabidopsis.org)

```
CLUSTAL O(1.2.4) multiple sequence alignment


MA_587505p0010      ------------------------------------------------------------      0
AT1G79620           MVGSNTVTATSRLLLICFAYSFTVFSMISSVTDPRDAAALRSLMDQWDNTPPSWGGSDDP      60


MA_587505p0010      ------------------------------------------------------------      0
AT1G79620           CGTPWEGVSCNNSRITALGLSTMGLKGRLSGDIGELAELRSLDLSFNRGLTGSLTSRLGD      120


MA_587505p0010      ------------------------------------------------------------      0
AT1G79620           LQKLNILILAGCGFTGTIPNELGYLKDLSFLALNSNNFTGKIPASLGNLTKVYWLDLADN      180


MA_587505p0010      ---------------------------------------------------MGGNGFHLS      9
AT1G79620           QLTGPIPISSGSSPGLDLLLKAKHFHFNKNQLSGTIPPKLFSSEMILIHVLFDGNRFTGS      240
                                                                   :.** *  *

MA_587505p0010      LILPIAVIITIAG--VMATGMKGKDESKCPYDLTVASRMIP-------------------      48
AT1G79620           IPSTLGLIQTLEVLRLDRNTLTG----KVPENLSNLTNIIELNLAHNKLVGSLPDLSDMK      296
                    :   :.:* *:    :  . :.*    * * :*:  :.:*

MA_587505p0010      SSCYANNTDQTIE-KNTCCWS-VFASYVFSISLYANRTGTVLLDANGS------RHCTN-      99
AT1G79620           SMNYVDLSNNSFDPSESPLWFSTLPSLTTLVMEYGSLQGPLPNKLFGFPQLQQVRLKKNA      356
                    *  *.: :::::: .::  *  .: * .  :  *..  * :  .  *       *  .*

MA_587505p0010      -----------------------EFVRFVRSRGLVQPGFFSNNSSCAINPLIFTAHAGPC      136
AT1G79620           FNGTLSLGDTVGPELQLVDLQDNDISSVTLSSGYTNTLILEGNPVCTTA-------LSNT      409
                                           ::  .. * * .:  ::..*  *:          .

MA_587505p0010      RYATLGDISNVIDVGNASIACDGPSLN-----ESKACERC--QEK-VISKALELLNLTNS      188
AT1G79620           NYCQIQQQQVKRIYSTSLANCGGKSCPLDQKVSPQSCECAYPYEGTLYFRGPMFRDLSNV      469
                    .*. : : .     ..:   *.* *       . ::** .   *  :  :.  : :*:*

MA_587505p0010      KEFVPCGMAITMGIWGRDP---SEE-------RFGGYVGCVLQVLGNIG-----------      227
AT1G79620           NTY----HSLEMSLWVKLGLTPGSVSLQNPFFNNDDYLQIQLALFPPMGKYFNRTEVQRI      525
                    : :     :: *.:* :     ..        . ..*:   * ::  :*

MA_587505p0010      GLGTSSLIPSPPPP--PLPAS----HEPSGGNKK-GRGRVRIAVAAGIAATSVVLLVFAA      280
AT1G79620           GFDLSNQTYKPPPLFGPYYFIASPYTFPADGNGHSLSSRMVTGIITGCSALVLCLVALGI      585
                    *:. *.   .***   *          *:.** :   .*:  .: :* :*  : *:.:.

MA_587505p0010      LIVLKSKKTNGPSEESTASSSLQVLTRSLMGIDKRTETVAREPLPTEGLYLFTKRELQRA      340
AT1G79620           YAMWQKRRAEQA----------IGLSRPFVSWASSGKDS-GGAPQLKGARWFSYEELKKI      634
                      : :.::::              *:* ::.  .  :         :*   *: .**::

MA_587505p0010      TEDFSEARVLGTGGSGKVYLGKLPSGKQVAIKRIVQKRK--VDEFYKEVELLSRLRHPNL      398
AT1G79620           TNNFSVSSELGYGGYGKVYKGMLQDGHMVAIKRAQQGSTQGGLEFKTEIELLSRVHHKNL      694
                    *::** :  ** ** **** * * .*: *****  *  .    ** .*:*****::* **
                         638-912: Domain: Protein kinase

MA_587505p0010      ASLLGYCQHRREHILVYEYMPGGDLGCLLFRREDPVLPWDRRLRIALDCAEGLAHLHLFP      458
AT1G79620           VGLVGFCFEQGEQILVYEYMSNGSLKDSLTGRSGITLDWKRRLRVALGSARGLAYLHELA      754
                    ..*:*:* .: *:******* .*.*   *  *.. .* *.****:**..*.***:** :

MA_587505p0010      EGAIVHRDIKPTKLSDFGISKLIAPDISHASTEIKGTTGFLDPEYFFVGKLNILLNESGQ      518
AT1G79620           DPPIIHRDVKST---------------------------------------NILLDENLT      775
                    :  *:***:* *                                       ****:*.
```

```
MA_587505p0010      AKLSDFGVSKLIAP-DFSHASTEIKGTTGYVDPEYFTVGRLTDASDVYSFGVVLLQLISG     577
AT1G79620           AKVADFGLSKLVSDCTKGHVSTQVKGTLGYLDPEYYTTQKLTEKSDVYSFGVVMMELITA     835
                    **::***:***::    .*.**::*** **:****:*. :**: *********:::**:.

MA_587505p0010      QKAVISTPSGGADSVVYMAHVFMSGEDPDVRNLVDPRIADAMDARDLGSIK---MVLDIA     634
AT1G79620           KQPIEKG-----KYIVREIKLVMNKSDDDFYGLRDKMDR---SLRDVGTLPELGRYMELA     887
                    :: : .       . :*   ::.*. .* *. .* *       . **:*::      :::*

MA_587505p0010      Y-----------------------------------------------------------     635
AT1G79620           LKCVDETADERPTMSEVVKEIEIIIQNSGASSSSSASASSSATDFGEKLLYGGTLKKKEA     947


MA_587505p0010      ------------------------       635
AT1G79620           RDGDGGGAFDYSGGYSVPTKIEPK       971
```

**Fig.S21 Domains of MYB DOMAIN PROTEIN 3 (MYB3)**

**Missense polymorphism sites are highlighted in green in the Norway spruce gene sequences**

**Well characterized domains of the genes are highlighted in other colors in the *Arabidopsis thaliana* gene sequences**

**MA_7115g0010** [293 residues] Norway spruce (http://congenie.org/)

**AT1G22640** [257 residues] *Arabidopsis thaliana* (https://www.arabidopsis.org)

```
CLUSTAL O(1.2.4) multiple sequence alignment


MA_7115p0010      MG-RSCCSKEGLNRGAWSRKEDMILSEYIRIHGDGGWTNLPQRAGLKRCGKSCRLRWTNY      59
AT1G22640         MGRSPCCEKAHMNKGAWTKEEDQLLVDYIRKHGEGCWRSLPRAAGLQRCGKSCRLRWMNY      60
                  **   **.*  :*:***:::** :* :*** **:* * .**: ***:********** **
                          9-61: Myb-type1 HTH DNA-binding domain

MA_7115p0010      LRPDIKLGNISPDEDELIIRMHRLLGNRWSLIAGRLPGRTDNEIKNYWNTRLSKKLLSID      119
AT1G22640         LRPDLKRGNFTEEEDELIIKLHSLLGNKWSLIAGRLPGRTDNEIKNYWNTHIKRKLLSRG      120
                  ****:* **:: :******::* ****:**********************::..:**** .
                          62-116: Myb-type2 HTH DNA-binding domain

MA_7115p0010      DSQSKSTARNLETSSKSPPPPPNHVFKTTPIKITTAVKFSETVGPKRCNG-YG-RSNCSS      177
AT1G22640         IDPNS--HRLINESVVSPSSLQNDVVETIHL------DFSGPVKPEPVREEIGMVNNCES      172
                   . ..   * :: *  **    *.*.:*  :      .**  * *:  .   *  .**.*

MA_7115p0010      AEAIKLCNIKENTNSIPHDDSEHIVFDMT----------DVNLLETEAGNCTSAIWSLEE      227
AT1G22640         SGT---------TSEKDYGNEEDWVLNLELSVGPSYRYESTRKVSVVDSAESTRRWGSE-      222
                  : :         *..  :.:.*. *:::           ... :..  .  ::  *. *

MA_7115p0010      ERSPHSYYFAVDTATLDESLSELNVLSSPDCNLLYSASGSSTDFELEEFYREAATLATGI      287
AT1G22640         -------LFGAHESDAVCLCCRIGLFRNESCRNC----------RVSDVRTH--------      257
                          *... :      ..:.:: . .*.            .:.:.  .

MA_7115p0010      NDSVFT  293
AT1G22640         ------  257
```

**Fig.S22 Domains of RESISTANT TO P. SYRINGAE 2 (RPS2)**
**Missense polymorphism sites are highlighted in green in the Norway spruce gene sequences**
**Well characterized domains of the genes are highlighted in other colors in the *Arabidopsis thaliana* gene sequences**
**MA_475302g0010** [303 residues] Norway spruce (http://congenie.org/)
**AT4G26090** [910 residues] *Arabidopsis thaliana* (https://www.arabidopsis.org)

```
CLUSTAL O(1.2.4) multiple sequence alignment


MA_475302p0010      ------------------------------------------------------------     0
AT4G26090           MDFISSLIVGCAQVLCESMNMAERRGHKTDLRQAITDLETAIGDLKAIRDDLTLRIQQDG     60

                      29-58: Coiled coil domain - essential for the resistance to AvrRpt2
MA_475302p0010      ------------------------------------------------------------     0
AT4G26090           LEGRSCSNRAREWLSAVQVTETKTALLLVRFRRREQRTRMRRRYLSCFGCADYKLCKKVS     120


MA_475302p0010      ------------------------------------------------------------     0
AT4G26090           AILKSIGELRERSEAIKTDGGSIQVTCREIPIKSVVGNTTMMEQVLEFLSEEEERGIIGV     180

                         135-440: NB-ARC domain - nucleotide-binding domain
MA_475302p0010      ------------------------------------------------------------     0
AT4G26090           YGPGGVGKTTLMQSINNELITKGHQYDVLIWVQMSREFGECTIQQAVGARLGLSWDEKET     240


MA_475302p0010      ------------------------------------------------------------     0
AT4G26090           GENRALKIYRALRQKRFLLLLDDVWEEIDLEKTGVPRPDRENKCKVMFTTRSIALCNNMG     300


MA_475302p0010      ------------------------------------------------------------     0
AT4G26090           AEYKLRVEFLEKKHAWELFCSKVWRKDLLESSSIRRLAEIIVSKCGGLPLALITLGGAMA     360


MA_475302p0010      ------------------------------------------------------------     0
AT4G26090           HRETEEEWIHASEVLTRFPAEMKGMNYVFALLKFSYDNLESDLLRSCFLYCALFPEEHSI     420


MA_475302p0010      ------------------------------------------------------------     0
AT4G26090           EIEQLVEYWVGEGFLTSSHGVNTIYKGYFLIGDLKAACLLETGDEKTQVKMHNVVRSFAL     480


MA_475302p0010      ------------------------------------------------------------     0
AT4G26090           WMASEQGTYKELILVEPSMGHTEAPKAENWRQALVISLLDNRIQTLPEKLICPKLTTLML     540

                 512-533: Leucine-rich repeat 1 (LRR) - probably act as specificity determinant
of pathogen recognition
MA_475302p0010      ---MKLKEIP-----------ALDIGNQAAHGGFPMSNAKTMLQLQILKIQGCRVLKTFR     46
AT4G26090           QQNSSLKKIPTGFFMHMPVLRVLDLSFTS-ITEIPLSI-KYLVELYHLSMSGTKIS--VL     596
                        .**:**           .**:.  :    :*:*  * :::*  *.:.* ::  .
                         534-556: LRR 2            559-580: LRR 3
MA_475302p0010      VEELPNLRKLG---ISHCAELKELSMES-----------------------------GG     73
AT4G26090           PQELGNLRKLKHLDLQRTQFLQTIPRDAICWLSKLEVLNLYYSYAGWELQSFGEDEAEEL     656
                     :** *****    :.:   *: :  ::
                      582-604: LRR 4     605-627: LRR 5
MA_475302p0010      GLLRLERL--------DLWSLQRLESIVWNAKTMLQIQFFGISNCHVLKTFGVEELP---     122
AT4G26090           GFADLEYLENLTTLGITVLSLETLKTLFEFGALHKHIQHLHVEECNELLYFNLPSLTNHG     716
                    *:  ** *         : **: *:::.  .    :**.: :.:*: *  *.: .*

MA_475302p0010      -SLKEINISNCVELKESFM---ESGGSLPMLEILELYSLEIRSIVWNEKTM----LQLQS     174
AT4G26090           RNLRRLSIKSCHDLEYLVTPADFENDWLPSLEVLTLHSLHNLTRVWGNSVSQDCLRNIRC     776
                     .*:.:.*..* :*:  .     ... ** **:* *:**.  : **.:..      :::.

MA_475302p0010      FYISYCRVLKTF-RVEELPNLKEIEIKQCPELQVQIGS----------VPMLKKLTLEG     222
AT4G26090           INISHCNKLKNVSWVQKLPKLEVIELFDCREIEELISEHESPSVEDPTLFPSLKTLRTRD     836
```

```
                    : **:*. **..  *::**:*: **: :* *::  *..           .* **.*  ..

MA_475302p0010      LKKSESIARASSVWNKETMPKLEYINITGCPLLRRLPMEMDK-LPNLKKIVGEVGWWEGI     281
AT4G26090           LPELNSI-----LPSRFSFQKVETLVITNCPRVKKLPFQERRTQMNLPTVYCEEKWWKAL     891
                    * : :**     : .: :: *:* : **.** :::**::  :   ** .:  *  **:.:

MA_475302p0010      NWEIDNVKIKLSELFRDEDEDD 303
AT4G26090           EKDQPNEELCYLPRFVPN---- 909
                    : :  * ::     *  :
```
